# Xenoglycosylation of the HIV-1 Glycan Shield Promotes Cross-Clade Neutralization and Diverse Human Antibody Lineages

**DOI:** 10.64898/2026.10.08.757767

**Authors:** Linling He, Xiaohe Lin, Yi-Nan Zhang, Maddy L. Newby, Connor DesRoberts, Joel D. Allen, Robyn L. Stanfield, Max Crispin, Ian A. Wilson, Jiang Zhu

## Abstract

Despite advances in HIV-1 envelope glycoprotein (Env)-based vaccine design, eliciting broadly neutralizing antibodies (bNAbs) remains a central challenge. Here, we identify core α(1,3)-fucose as a key modulator of neutralizing antibody (NAb) induction and breadth. Uncleaved prefusion-optimized (UFO) trimers and self-assembling protein nanoparticles (SApNPs) produced in High Five (Hi5) insect cells incorporate core α(1,3)-fucose into the Env glycan shield. Crystal structures of Hi5-produced BG505 UFO trimers in complex with bNAbs confirm native-like Env architecture and epitope presentation. In wild-type mice, these trimers and SApNPs elicit robust autologous and heterologous NAb responses. We then engineer Chinese hamster ovary (CHO) cells to co-express three insect- and plant-derived core α(1,3)-fucosyltransferases. The resulting immunogens carry fucosylated mammalian glycans and elicit cross-clade NAb responses in mice, supporting an important role for core α(1,3)-fucose in broadened neutralization. Env-specific IgE responses suggest engagement of associated immune pathways during vaccination. In ATX-Gx humanized mice, core α(1,3)-fucosylated, glycan-trimmed Env elicits detectable cross-clade neutralization by week 11. Single-cell analysis identifies distinct human antibody lineages with cross-clade activity and noncanonical Env recognition. Repertoire and functional analyses uncover putatively related lineage members across two animals within the same group. Together, these findings support xenoglycosylation as a promising, germline-agnostic strategy for HIV-1 vaccine design.

**ONE-SENTENCE SUMMARY:** Engineering the HIV-1 glycan shield with core α(1,3)-fucose promotes cross-clade neutralization

## INTRODUCTION

As of 2024, an estimated 40.8 million people are living with HIV-1 worldwide, with approximately 1.3 million new infections annually (*1*). HIV-1 continues to impose a substantial global health burden, disproportionately affecting regions where poverty, social inequality, and limited healthcare infrastructure constrain prevention and treatment (*2–4*). Despite decades of research, no licensed vaccine is available (*5*). Meanwhile, long-acting antiretroviral agents have emerged as effective options for HIV-1 pre-exposure prophylaxis (PrEP). Notably, the first-in-class capsid inhibitor lenacapavir has demonstrated safety and high efficacy as a twice-yearly subcutaneous regimen in phase 3 trials across diverse populations (*6–8*), with once-yearly formulations under evaluation in phase 1 studies (*9*). Although these advances are promising, a safe, effective, and broadly protective HIV-1 vaccine remains an important long-term goal.

The HIV-1 envelope glycoprotein (Env) mediates viral entry and is the primary target of humoral immunity, making it central to vaccine development. Functional Env forms a homotrimer of noncovalently associated gp120–gp41 heterodimers generated by proteolytic cleavage of a gp160 precursor (*10*). HIV-1 evades antibody recognition through multiple mechanisms, including rapid immune escape driven by an error-prone reverse transcriptase (*11*), Env metastability that diverts responses toward non-neutralizing epitopes (*12*), and a dense host-derived glycan shield that masks much of the Env surface and limits antibody access (*13*). Nevertheless, up to 30% of people living with HIV-1 eventually develop serum neutralizing activity against diverse circulating strains. Since 2009, multiple families of broadly neutralizing antibodies (bNAbs) targeting distinct sites of vulnerability have been isolated (*14–17*), and structural analyses of Env–bNAb complexes have enabled antibody-guided rational vaccine design (*18–21*). Yet, a vaccine that reproducibly elicits bNAb responses in humans remains elusive.

Like other viruses that utilize a class I fusion mechanism, HIV-1 Env adopts a metastable prefusion conformation (*22*). This intrinsic Env instability can lead to misfolding, gp120 shedding, and exposure of gp41 stumps on the virion surface (*12*), generating immunodominant decoys that divert antibody responses away from functional Env spikes. A major milestone in HIV-1 vaccine research was the development of the cleaved, soluble BG505 SOSIP.664 trimer (*23*), which closely mimics native Env (*24–27*). Subsequent structural studies using X-ray crystallography and cryo-electron microscopy (cryo-EM) (*28, 29*) established SOSIP trimers as both a foundation and a catalyst for modern HIV-1 Env vaccine design (*30*). The natively flexibly linked (NFL) trimer (*31–33*) and the uncleaved prefusion-optimized (UFO) trimer (*34, 35*) further advanced the field (*36, 37*). Among these, the UFO design directly addresses a key determinant of Env metastability**―**the N-terminal bend in heptad repeat 1 (HR1_N_, residues 547-569)**―**through targeted optimization (*34*). Both UFO and UFO-BG designs, with the latter incorporating BG505 gp41, have since been applied to stabilize HIV-1 Envs from diverse strains and subtypes (*35, 38*).

Delivering Env trimers in multimeric formats that mimic virus-like particles (VLPs) has emerged as a promising strategy for HIV-1 vaccine design (*39–45*). VLPs elicit potent immune responses (*46–53*), owing to their particulate size, repetitive antigen display, efficient lymph node trafficking, and robust induction of germinal centers (GCs) (*42, 54*). Protein nanoparticles (NPs), which retain key VLP-like properties but are more amenable to engineering, provide versatile scaffolds for vaccine design. SpyTag/SpyCatcher-mediated conjugation (*55, 56*), two-component nanoparticles (*57*), and single-component self-assembling protein nanoparticles (1c-SApNPs) (*35, 58, 59*) have been evaluated for HIV-1 Env display (*60–65*). In our 1c-SApNP design, the Env trimer, NP scaffold, locking domain (LD), and PADRE helper T-cell epitope are encoded by a single gene and self-assemble into a multilayered particulate architecture (*65*). This platform has been applied to multiple viral targets, including filovirus glycoprotein (GP) (*59, 66*), SARS-CoV-2 spike (S) (*67, 68*), and influenza hemagglutinin (HA) and matrix protein 2 ectodomain (M2e) (*69–71*). BG505 UFO trimer-presenting 1c-SApNP vaccines have advanced to a phase 1 clinical trial (ClinicalTrials.gov: NCT06541093) evaluating safety and immunogenicity.

Despite advances in Env trimer stabilization and NP immunogen design, the glycan shield remains a major obstacle to HIV-1 vaccine development because of its multifaceted roles in Env– bNAb interactions (*72–76*). Glycans block immune recognition of neutralizing epitopes (*13*), and mannose-rich moieties may exert immunosuppressive effects (*77, 78*). At the same time, they form integral components of several bNAb epitopes, including those at the V1V2 apex and V3 stem (*79, 80*), and can promote lymph node targeting of HIV-1 NP immunogens (*81, 82*). Env glycans also shape strain-specific immunodominance: in rabbits, autologous tier 2 NAbs often target specific holes within the glycan shield (*83–89*), whereas in rodents and nonhuman primates (NHPs), BG505 Env-induced autologous NAbs tend to recognize glycan-rich C3/V4 and C3/V5 regions (*90–93*). To address this complexity, some strategies aim to recapitulate native Env glycosylation (*94*), whereas others use sequential immunization to remove and later restore glycans surrounding bNAb epitopes (*95–98*). We recently reported that enzymatic trimming of oligomannose-type glycans by endoglycosidase H (Endo H) enhanced tier 2 NAb elicitation and responder frequency in multiple animal models (*65*). However, glycan trimming had little effect on neutralization breadth, leaving the induction of breadth as a persistent challenge for HIV-1 vaccine development.

Here, we investigated an alternative strategy to overcome the glycan shield barrier to HIV-1 bNAb elicitation. We first optimized a plasmid-based transient expression system in High Five (Hi5) insect cells for efficient immunogen production. Hi5-produced BG505 UFO trimers and 1c-SApNPs exhibited high yield, purity, and stability and incorporated core α(1,3)-fucose into the glycan shield. They also displayed distinct antigenic profiles when tested against a broad panel of bNAbs, NAbs, and non-NAbs. Crystal structures of unmodified and glycan-trimmed Hi5-derived BG505 UFO trimers, determined at 4.46 and 4.09 Å resolution in complex with two bNAb pairs, revealed native-like conformations closely resembling mammalian cell-expressed Env. In BALB/c mice, these immunogens elicited NAbs against variants representing subtypes A, B, C, and G and recombinant forms CRF07_BC and CRF01_AE. We then transiently co-expressed BG505 Env immunogens with three insect- or plant-derived core α(1,3)-fucosyltransferases (FucTs) in Chinese hamster ovary (CHO) cells (termed CHO^FucT3^), introducing core α(1,3)-fucose into mammalian glycans. Whereas ExpiCHO-expressed BG505 UFO trimers failed to elicit detectable tier 2 NAb responses, their core α(1,3)-fucosylated counterparts induced robust autologous and heterologous NAb responses in wild-type mice. IgE reactivity, potentially associated with core α(1,3)-fucose in insect and plant allergens, was detected during immunization. Finally, ATX-Gx humanized mice were immunized with Hi5- and CHO^FucT3^-derived BG505 UFO trimers to assess the impact of core α(1,3)-fucosylation on human antibody responses. Env-specific single-cell sorting and antibody cloning, followed by functional validation, identified distinct human antibody lineages with cross-clade neutralizing activity and unconventional Env interactions. Repertoire analyses revealed NAb lineage expansion and diversification in one animal and the presence of a putatively related lineage in a second animal within the same group. This study suggests that xenoglycosylation may promote cross-clade NAb responses without requiring predefined bNAb developmental pathways.

## RESULTS

### Plasmid-based insect cell expression and characterization of HIV-1 BG505 Env immunogens

The insect cell–baculovirus expression vector system (IC-BEVS) is widely used for recombinant protein production (*99–101*) and supports industrial-scale vaccine manufacturing (*102, 103*), as exemplified by the licensed vaccines Flublok® and Cervarix® (*104–106*). However, IC-BEVS has several inherent limitations, including non-mammalian glycosylation, susceptibility to proteolytic degradation, suboptimal yields, and the risk of baculoviral contamination (*107–110*). Plasmid-based expression (PBE) systems have recently emerged as baculovirus-free alternatives for protein expression in insect cells (*111–115*), offering a simplified workflow and shorter production timelines (*116, 117*) (**Fig. 1a**). Insect cells typically produce paucimannose glycans rather than oligomannose-, hybrid-, and complex-type structures generated by mammalian cells (*118–120*) (**Fig. 1b**). Among commonly used insect cell lines, *Trichoplusia ni* (Hi5) cells achieve higher recombinant protein yields than *Spodoptera frugiperda* (Sf9) cells (*121*). Thus, an optimized PBE system in Hi5 cells may provide a streamlined platform for producing HIV-1 Env immunogens with shorter, paucimannose-type glycans that resemble trimmed mammalian glycans (*65*).

**Fig. 1.**
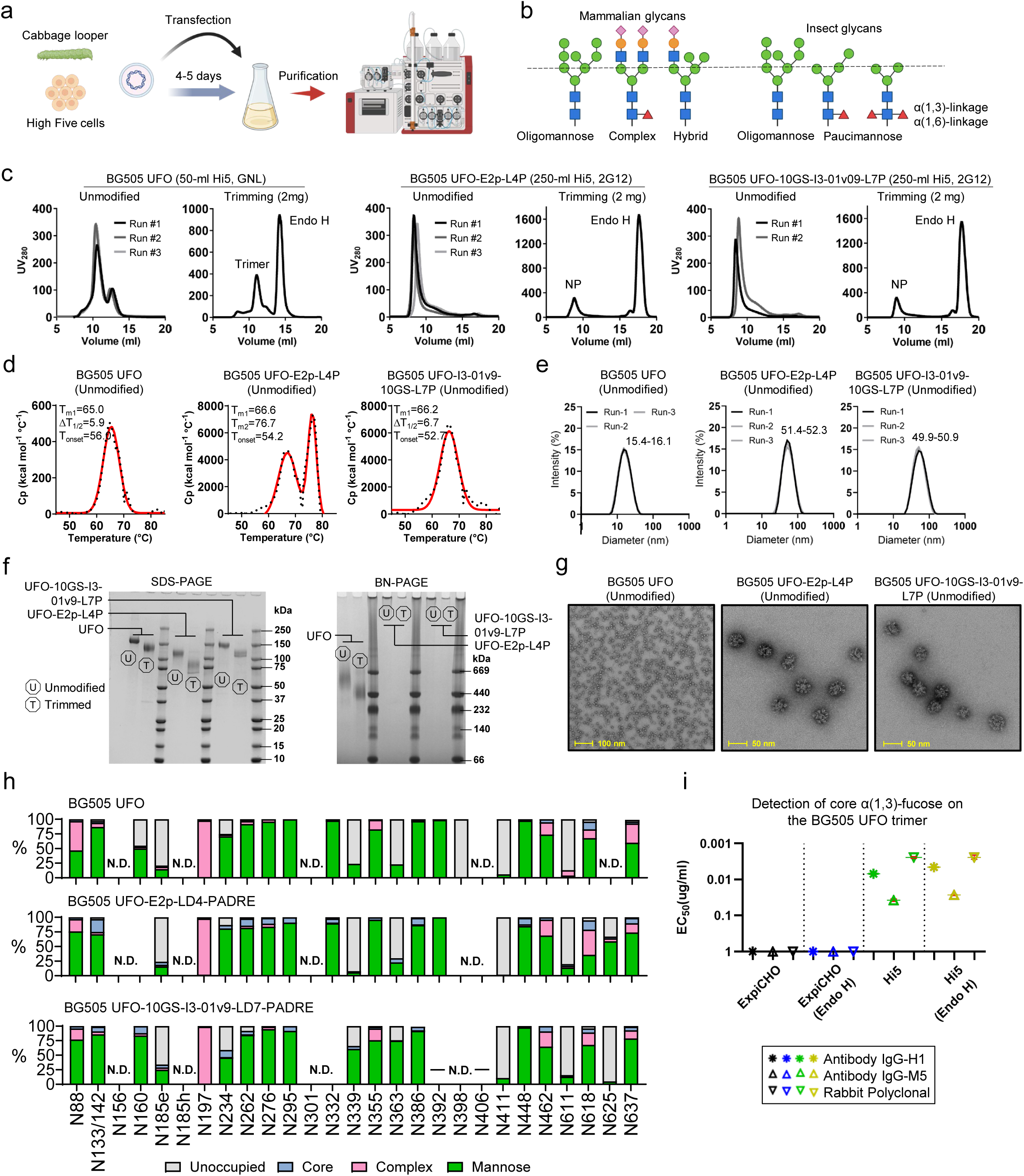
Hi5 insect cell expression and in vitro characterization of BG505 Env immunogens. (**a**) Schematic of the baculovirus-free, plasmid-based expression (PBE) platform for recombinant protein production in High Five (Hi5) insect cells. (**b**) Comparison of representative glycoforms generated by mammalian cells (oligomannose-, hybrid-, and complex-type glycans) versus insect cells (oligomannose-type and paucimannose-type glycans with core α(1,3)-fucose). (**c**) SEC profiles of Hi5-expressed BG505 UFO trimer (left), UFO-E2p-L4P SApNP (middle), and UFO-10GS-I3-01v9-L7P SApNP (right) in unmodified or glycan-trimmed forms. Data represent two to three independent Hi5 expression experiments. Trimers were purified by GNL chromatography, and SApNPs by immunoaffinity chromatography (IAC) using a 2G12 antibody column. For glycan trimming, 2 mg of BG505 Env protein was treated with Endo H before SEC analysis. (**d**) DSC thermograms of Hi5-expressed, unmodified BG505 Env immunogens. Experimental data and Gaussian fits are shown as black dots and red lines, respectively. Thermal denaturation midpoints (T_m_ or T_m1–2_), onset temperatures (T_onset_), and half-widths at half-maximum (ΔT_1/2_) are indicated. (**e**) Particle size distributions of Hi5-expressed, unmodified BG505 Env immunogens from three independent production batches. Hydrodynamic diameters (D_h_), measured by DLS, are indicated. (**f**) SDS-PAGE (reducing) and BN-PAGE analyses of Hi5-expressed BG505 Env immunogens. Both unmodified and glycan-trimmed samples were analyzed. (**g**) Representative nsEM images of Hi5-expressed, unmodified BG505 Env immunogens. Scale bars, 100 and 50 nm, as indicated. (**h**) Site-specific glycan analysis of Hi5-expressed BG505 Env immunogens. Glycan compositions are categorized by type: complex (pink), mannose (green), core (blue), and unoccupied (gray). Unresolved glycan sites are denoted as “N.D.” (**i**) ELISA-derived EC_50_ (µg/mL) values for ExpiCHO- and Hi5-expressed BG505 UFO trimers (unmodified or glycan-trimmed) binding to core α(1,3)-fucose-specific monoclonal antibodies (IgG-H1 and IgG-M5) and a rabbit polyclonal antibody. For samples with absorbance at 450 nm (A_450_) < 0.5 at the maximum antibody concentration (1 μg/mL), binding was considered negligible, and EC_50_ was set to 1 μg/mL. The insect cell expression and glycan schematics were created with BioRender.com.

Here, we adopted a recently reported PBE system (*114, 116, 117*) to transiently express three BG505 Env immunogens in Hi5 insect cells: a UFO trimer and two multilayered SApNPs based on the E2p and I3-01v9 scaffolds (*65*). Trimers were purified by *Galanthus nivalis* lectin (GNL) affinity chromatography, whereas SApNPs were isolated by 2G12-based immunoaffinity chromatography (IAC). The immunogens were analyzed with or without Endo H treatment to trim oligomannose-type glycans (*65*). Size-exclusion chromatography (SEC) revealed distinct elution profiles (**Fig. 1c**). Compared with ExpiCHO (*35, 65*), PBE/Hi5 produced ∼fivefold more trimer, whereas SApNP yields were 55–65% lower, based on ultraviolet absorbance at 280 nm (UV_280_). The Hi5 trimers contained an additional monomer peak (∼12.7 mL), whereas ExpiCHO trimers eluted predominantly as a single species (*35*). Hi5-produced SApNPs exhibited broader SEC peaks than their ExpiCHO counterparts. These elution profiles were unaffected by Endo H treatment, indicating that the differences primarily reflected intrinsic properties of the expression systems rather than glycan trimming. Thermostability was largely preserved between Hi5 and ExpiCHO, as indicated by closely matched differential scanning calorimetry (DSC) profiles (**Fig. 1d**). The BG505 UFO trimer exhibited a lower melting temperature in Hi5 (T_m_ = 65.0 °C) than in ExpiCHO (68.3 °C), whereas the difference in T_m_ was minimal for SApNPs (ΔT_m_ < 1 °C). Endo H treatment slightly increased trimer stability but modestly reduced that of SApNPs, with minor effects overall (**Fig. S1a**). Dynamic light scattering (DLS) showed a hydrodynamic diameter (Dh) of ∼16.0 nm for Hi5 trimers, with SApNP size distributions comparable to those of ExpiCHO material (*65*) (**Fig. 1e**). Glycan trimming resulted in a small decrease in Dh values (**Fig. S1b**). All immunogens were analyzed by sodium dodecyl sulfate–polyacrylamide gel electrophoresis (SDS-PAGE) under reducing conditions and by blue native–polyacrylamide gel electrophoresis (BN-PAGE) (**Fig. 1f**). In SDS-PAGE, glycan trimming caused a greater reduction in apparent molecular weight for Hi5-derived than ExpiCHO-derived immunogens, suggesting a higher abundance of Endo H-susceptible glycans in the former. In BN-PAGE, E2p and I3-01v9 SApNPs were retained in the loading wells due to their large size, supporting intact particle assembly. Finally, SEC-purified Hi5 immunogens were visualized by negative-stain EM (nsEM), revealing triangular, propeller-like trimers and well-formed particles with ordered surface arrays (**Fig. 1g**; **Fig. S1c**), consistent with previous observations for ExpiCHO-produced immunogens (*65*).

Together, these results establish PBE/Hi5 as a rapid and efficient platform for producing HIV-1 Env immunogens. Compared with IC-BEVS (∼4 weeks) and ExpiCHO (∼2 weeks), the PBE/Hi5 system developed here enables production within 4–5 days while achieving higher trimer yields than ExpiCHO, with lower SApNP yields but comparable assembly efficiency (*65*).

### Glycan analysis of Hi5 insect cell–produced HIV-1 BG505 Env immunogens

Liquid chromatography–mass spectrometry (LC–MS) has been widely used to characterize *N-* linked glycosylation profiles of native-like Env trimers produced in mammalian cells (*122–124*). Previously, we analyzed ExpiCHO-produced BG505 Env with or without Endo H treatment (*35, 65*). Because the glycosylation machinery differs between insect and mammalian cells (*118, 119*), we determined site-specific glycosylation and glycan occupancy of BG505 immunogens produced in Hi5 cells. Three aliquots of each sample were digested separately with trypsin, chymotrypsin, and α-lytic protease to generate fragments predominantly carrying a single *N*-linked glycosylation site, followed by glycopeptide analysis using established workflows (*35, 65*). Where possible, the same peptide sequence was used for each site across all samples to minimize bias arising from differences in ionization efficiency. In some cases, incomplete peptide coverage and unresolved or averaged sites limit site-specific assignment accuracy.

Overall, the distribution of mannose- and complex-type glycans across Hi5-produced UFO trimers largely resembled that of mammalian cell-expressed material, although mannose species were more abundant. Sites typically occupied by complex-type glycans, such as N88, N197, N462, and, to a lesser extent, N355, exhibited varying degrees of processing, with N197 showing full complex-type occupancy (**Fig. 1h**). The similar site-specific processing patterns may reflect greater accessibility of these glycosylation sites to glycan-processing enzymes in both expression systems. Glycan sites within the trimer-associated mannose patches (TAMPs) and the intrinsic mannose patch (IMP) retained high levels of mannose-type glycans (M4–M9), similar to mammalian cell-expressed trimers, whereas paucimannose species [(±Fuc)Man_3–4_GlcNAc_2_] were enriched at solvent-accessible mixed or complex-type sites (**Fig. S1d**). This shift is consistent with increased exposure of these positions to insect-derived mannosidases during the later stages of glycan processing. Reduced glycan occupancy was observed in the V2 and V3 regions (e.g., N156 and, to a lesser extent, N339) and at gp41 sites N611 and N625. Underoccupancy in and around the V3 region may increase access of V3-directed bNAbs to underlying peptide epitopes while preserving surrounding glycan architecture, whereas glycan holes in gp41 may promote engagement by non-NAbs. For N133 and N142, glycan composition was reported as an average due to detection of only double-occupied peptides. Site-specific data for N156, N301, and N406 were not fully resolved across samples and were only partially reported. Among insect cell lines, Hi5 adds core α(1,3)-fucose, a carbohydrate motif implicated in allergic responses to insect- and plant-derived glycoallergens (*125–128*), whereas Sf9 largely lacks this modification (*129–131*). To assess core α(1,3)-fucosylation of the Env glycan shield produced by the PBE system in Hi5 cells, we quantified glycans bearing one or two fucose residues (Fuc(1) or Fuc(2)) at each *N*-linked glycan site (**Fig. S1e**). The site-specific glycan compositions are summarized in **Fig. S1f–h**, but the LC–MS data do not directly resolve monosaccharide linkages (e.g., α(1,3) vs. α(1,6)); therefore, linkage assignments can be inferred only for select glycoforms. For example, α(1,3)-linked core fucose can be inferred for oligomannose-type glycans from M5 (HexNAc(2)Hex(5)) to core-trimmed structures (e.g., HexNAc(1)) when two fucose residues are present (Fuc(2)). In contrast, fucose linkage cannot be reliably assigned for glycoforms containing extended GlcNAc branches [e.g., HexNAc(3)Hex(x) and above]. Overall, Fuc(2)-containing glycans were detected at low levels across all Hi5 immunogens in both soluble trimer and SApNP formats.

Core α(1,3)-fucose is a cross-reactive carbohydrate determinant (CCD) recognized by allergy-associated IgE antibodies (*125–128*). To confirm its presence within the glycan shield of Hi5-produced BG505 Env, we used IgG versions of two CCD-specific IgE antibodies, H1 and M5 (*132*), together with a rabbit polyclonal antibody (RabbitPoly). The 1.8 Å crystal structure of M5 bound to a CCD-derived disaccharide (PDB: 5I8O) provides atomic-level insight into recognition of this motif by IgE (*132*). In enzyme-linked immunosorbent assays (ELISA), ExpiCHO- and Hi5-produced BG505 UFO trimers, with or without glycan trimming, were tested for binding to IgG-H1, IgG-M5, and RabbitPoly (**Fig. 1i**; **Fig. S1i**). As expected, ExpiCHO-produced trimers showed no detectable reactivity, consistent with the absence of core α(1,3)-fucose, whereas Hi5-derived trimers bound all three reagents with half-maximal effective concentration (EC_50_) values of 0.002– 0.04 μg/mL. Of the three, RabbitPoly exhibited the strongest binding, and IgG-H1 bound BG505 Env 5–6-fold more strongly than IgG-M5. Glycan trimming slightly enhanced H1 and M5 binding, suggesting that local glycan context and epitope accessibility influence antibody recognition of the core α(1,3)-fucose motif. Together, glycan profiling and CCD-specific antibody binding supported the presence of core α(1,3)-fucose in Hi5-produced HIV-1 Env immunogens.

### Antigenic characterization of Hi5 insect cell–produced BG505 Env immunogens

In our previous study, ExpiCHO-expressed BG505 UFO trimers and SApNPs—bearing wild-type, oligomannose-enriched, or Endo H–trimmed glycans—were evaluated against a broad panel of 23 bNAbs, NAbs, and non-NAbs (*65*). Here, we assessed Hi5-produced BG505 UFO trimers and two 60-mer SApNPs (unmodified or Endo H-trimmed) against the same antibody panel using bio-layer interferometry (BLI). Quantitation sensors were used to enable direct comparison between trimers and SApNPs while accounting for differences in avidity. Peak binding responses are summarized in **Fig. 2a**, with full sensorgrams across six antibody concentrations shown in **Fig. S2a–f**.

**Fig. 2.**
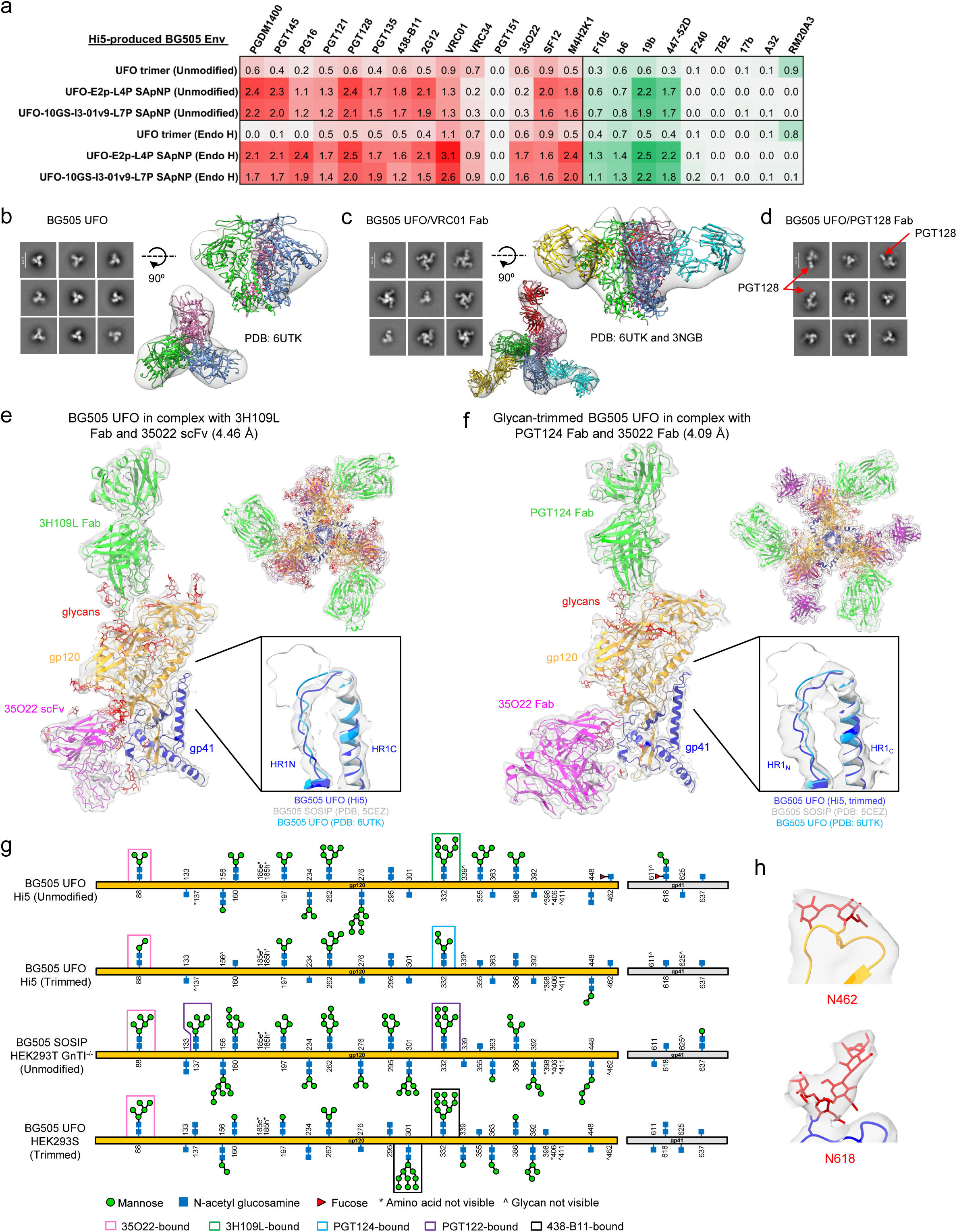
Antigenic and structural characterization of Hi5-expressed BG505 Env immunogens. (**a**) Antigenic profiles of Hi5-expressed BG505 UFO trimer and SApNPs in unmodified or glycan-trimmed forms. Fourteen bNAbs/NAbs and nine non-NAbs were evaluated by BLI on an Octet RED96 using six antigen concentrations (starting at 266 nM for the trimer and 5.6 nM for SApNPs, followed by 2-fold serial dilutions; sensorgrams in **Figs. S2a–f**). Peak responses at the highest concentration are shown in a color-coded matrix (green to red: weak to strong). (**b–d**) nsEM analysis of Hi5-expressed BG505 UFO trimer in the unbound state (**b**), in complex with VRC01 (**c**), and in complex with PGT128 (**d**). Representative 2D class averages are shown. For the unbound trimer, the BG505 UFO crystal structure (PDB: 6UTK) was fitted into the 3D reconstruction. For the VRC01 complex, the gp120/VRC01 structure (PDB: 3NGB) was docked onto BG505 UFO trimer (PDB: 6UTK) and fitted into the nsEM density. For the PGT128 complex, 3D reconstruction was not performed due to flexible binding and incomplete Fab occupancy. (**e– f**) Crystal structures of Hi5-expressed BG505 UFO trimers bound to bNAbs. Unmodified BG505 UFO was crystallized with 3H109L Fab and 35O22 scFv (**e**), and Endo H-treated BG505 UFO was crystallized with PGT124 and 35O22 Fabs (**f**). Side views show gp120 (gold), gp41 (blue), modeled glycans (red), and antibodies (green/magenta). Top views illustrate antibody occupancy and trimer arrangement. Insets highlight the shortened HR1 region characteristic of the UFO design compared with BG505 SOSIP (PDB: 5CEZ) and mammalian-expressed BG505 UFO (PDB: 6UTK). (**g**) Residue-specific schematic of N-linked glycans observed in crystal structures of BG505 Env expressed in insect and mammalian cells. Glycans modeled from electron density are mapped onto linear gp120 and gp41 sequences for Hi5-expressed BG505 UFO trimers in unmodified or glycan-trimmed forms. For comparison, glycan schematics are shown for BG505 SOSIP trimers from HEK293T GnTI-/- cells (PDB: 5FYL) and BG505 UFO trimers from HEK293S cells treated with Endo H after complex formation with Fabs (PDB: 6UTK). (**h**) Fucosylated N-linked glycans at N618 and N462 in unmodified Hi5 BG505 UFO, shown with electron density (gray surface) and protein backbone (gp120, gold; gp41, blue).

The Hi5 trimer recapitulated the overall antigenicity of the ExpiCHO material, showing robust reactivity to bNAbs and NAbs but minimal recognition by non-NAbs, except for the V3 antibodies 19b and 447-52D (*133–135*) and the base-binder RM20A3 (*136*) (**Fig. 2a**, top). Relative to the soluble trimer, SApNPs showed enhanced binding to bNAbs targeting the V2 apex (×3.6– 6.0), N332 supersite (×2.4–4.3), CD4bs (×1.4), and silent face (SF; ×1.8–2.2), but reduced binding to bNAbs recognizing the fusion peptide (FP; ×0.3–0.4) and gp120/gp41 interface (×0.3–0.5). For non-NAbs, NP display amplified signals at the CD4bs (×1.2–2.3) and V3 tip (×3.2–5.7), with little or no reactivity toward CD4-induced (CD4i), gp41, or base epitopes. The absence of RM20A3 binding to SApNPs was consistent with occlusion of the trimer base upon fusion to the NP scaffold. Across both expression systems, NP display enhanced recognition of apex-proximal epitopes but limited access to base-proximal sites occluded by neighboring trimers. Despite these shared trends, host-dependent effects were evident. Most bNAbs directed to the V2 apex and V3 stem (*137–139*) bound Hi5-derived BG505 Env less efficiently, consistent with reduced occupancy at N156 and N301 (**Fig. 1h**). PG16, which prefers α-2,6-sialylated hybrid- or complex-type glycans at N156 or N173 (*140*), showed reduced binding to the paucimannose-rich Hi5 trimer, whereas particulate display restored and enhanced recognition, with SApNPs yielding up to sixfold higher signals. The gp120 SF is enriched in high-mannose glycans, with N262, N295, and N448 as key contributors (*141*). At these sites, ExpiCHO and Hi5 Env exhibited broadly similar M8/M9 levels (**Fig. S1f– h**): 52–86%, 69–86%, and 70–87% for ExpiCHO versus 50–65%, 80–86%, and 65–88% for Hi5. Unexpectedly, NP display reduced SF12 binding to ExpiCHO-derived Env (×0.6) but enhanced recognition of Hi5-derived Env (×1.8–2.2), suggesting that reduced V2/V3 glycan occupancy in a paucimannose-rich environment may favor recognition by SF-directed bNAbs such as SF12 and VRC-PG05 (*141, 142*). PGT151, which targets the gp120–gp41 interface, prefers cleaved, native-like Env and requires complex-type glycans at N611 and N637 (*143, 144*). It bound weakly to the ExpiCHO UFO trimer but showed little or no reactivity toward SApNPs, likely due to steric hindrance on the NP surface (*65*). In Hi5-produced Env, near-complete loss of N611 occupancy was associated with loss of PGT151 binding across formats (**Fig. S1f–h**). Finally, bNAb VRC01 (*145, 146*) bound Hi5- and ExpiCHO-derived Env at comparable levels, whereas early CD4bs antibodies F105 and b6 showed stronger reactivity with Hi5 immunogens (**Fig. 2a**). The CD4-binding loop, V5 loop, and loop D form the recessed CD4bs rim, with V5 restricting access to the site (*146–148*). In ExpiCHO Env, N462 on V5 was occupied by complex-type and Man5 glycans, whereas in Hi5 Env it was dominated by paucimannose species (HexNAc(2)Hex(3)Fuc(1)), likely favoring b6 and F105 binding. N363 and N386 flank the CD4bs (*146–148*), and N363 was largely underoccupied in Hi5 immunogens, potentially increasing access for F105. Furthermore, reduced occupancy at distal N156 may promote transient Env opening and increase access for CD4bs-directed non-NAbs and NAbs with alternative angles of approach.

Endo H treatment reduced trimer binding to V2 apex bNAbs to negligible levels but had distinct effects on SApNP antigenicity. A similar effect was observed in the previous studies of ExpiCHO-produced immunogens (*65*). After glycan trimming, the most pronounced enhancement was observed for VRC01 (×2.4–2.8 in Hi5 versus ×2.2–2.3 in ExpiCHO). For PG16, which targets a Man5 glycan at N160 and prefers a complex- or hybrid-type glycan at N156 or N173 (*140*), unmodified SApNPs bound more strongly (×3.4–4.6) than the soluble trimer. Upon trimming, trimer reactivity was abolished, whereas SApNPs showed markedly increased PG16 signals (1.6– 2.2-fold higher than untrimmed particles), likely reflecting altered glycan recognition combined with avidity effects. For interface-directed bNAbs, PGT151 binding remained undetectable, but glycan trimming enhanced Env recognition by 35O22 (*149*), with SApNPs showing 2.7–2.8-fold stronger binding than the trimer. A similar effect was observed for the mouse NAb M4H2K1, which targets a dense C3/V4 glycan patch (*91*). Among non-NAbs, the V3 antibodies 19b and 447-52D recognized glycan-trimmed SApNPs more strongly than unmodified particles. Increased binding of the CD4bs-directed antibodies F105 and b6 to these SApNPs likely reflects reduced glycan shielding at the CD4bs together with NP-mediated avidity.

### Structural characterization of Hi5 insect cell–produced BG505 UFO trimer

Structural biology techniques, particularly X-ray crystallography and electron microscopy (EM), have been instrumental in the rational design of HIV-1 vaccines (*37*). We previously used nsEM to validate HIV-1 Env trimers with HR1_N_ or UFO designs (*34, 65, 150, 151*), and subsequently applied this approach to diverse viral antigens, including pneumovirus fusion protein (F), filovirus GP, influenza hemagglutinin (HA), and hepatitis C virus (HCV) E1E2 heterodimers (*66, 71, 152, 153*). Crystal structures of HR1_N_- or UFO-stabilized HIV-1 Env trimers produced in mammalian expression systems have been determined at 3.4–6.6 Å resolution (*34, 35, 91, 154*). However, structural information for insect cell–produced HIV-1 Env trimers has remained limited.

We first assessed the structural integrity of Hi5-produced BG505 UFO trimers using an established nsEM workflow (**Fig. S2g**). Two-dimensional (2D) class averages revealed propeller-like projections characteristic of native-like Env, with minor populations of dissociated species observed in excluded classes (**Fig. 2b**; **Fig. S2h**). Three-dimensional (3D) reconstruction and rigid-body fitting of the BG505 UFO trimer (PDB: 6UTK) (*154*) into the density map showed excellent agreement, confirming a native-like, prefusion-closed conformation. In complex with the bNAb VRC01, the trimers displayed near-complete threefold Fab occupancy, with no unbound species detected (**Fig. 2c**; **Fig. S2h**), consistent with comparable VRC01 binding to ExpiCHO- and Hi5-derived Env (**Fig. 2a**). A trimer–VRC01 model generated by docking the gp120/VRC01 complex (PDB: 3NGB) (*145*) onto the BG505 UFO trimer (PDB: 6UTK) (*154*) closely matched the density map. In contrast, complexes with the V3 glycan–directed bNAb PGT128 frequently showed only one or two bound Fabs per trimer (**Fig. 2d**; **Fig. S2h**). This pattern resembles that of glycan-trimmed, ExpiCHO-produced trimers (*65*), suggesting that shorter paucimannose glycans may have similar effects on antibody recognition of the V3 glycan patch as Endo H-mediated trimming.

To obtain higher-resolution insight into native-like Env expressed in insect cells, we determined two crystal structures of BG505 UFO trimers in complex with bNAbs. Trimers were expressed using PBE/Hi5 and purified by GNL affinity chromatography followed by SEC. In the first complex, an unmodified trimer (without Endo H treatment) was co-crystallized with the Fab of 3H109L—an inferred PGT122 precursor (*155*)—and the single-chain variable fragment (scFv) of the gp120/gp41 interface–targeting bNAb 35O22 (*149*), yielding a structure at 4.46 Å resolution (**Fig. 2e**). In the second complex, the trimer was treated with Endo H and crystallized with the N332 supersite-targeting PGT124 Fab (*156*) and 35O22 Fab, yielding a structure refined to 4.09 Å resolution (**Fig. 2f**). Both bNAb-bound BG505 UFO structures revealed compact, prefusion-closed Env conformations closely matching the mammalian-derived reference (PDB: 6UTK) (*154*). Notably, both complexes featured the shortened gp41 HR1_N_ characteristic of the UFO trimer design (*34*). The Cα root-mean-square deviation (RMSD) relative to 6UTK was 1.17 Å and 1.19 Å for the untrimmed and Endo H–treated trimers, respectively, demonstrating that native-like Env folding and quaternary organization were preserved in Hi5-produced trimers.

We next examined site-specific glycan features in the two Hi5-derived Env structures (**Fig. 2g**; **Figs. S2i–j**). *N*-linked glycan sites that retained more than a single GlcNAc residue in the Endo H–treated structure were interpreted as less accessible to the enzyme in the native trimer. However, the absence of modeled glycan density does not exclude the presence of extended glycan chains, which may be conformationally flexible and therefore invisible at this resolution. In the untrimmed Hi5 trimer, glycans could be modeled at most sites observed in the mammalian BG505 SOSIP reference, consistent with broadly conserved glycan positioning across systems (*157*). In the Endo H–treated Hi5 trimer, most sites appeared to be reduced to single-GlcNAc stubs, with only seven sites retaining more than one GlcNAc residue. By comparison, the mammalian Endo H–treated BG505 UFO reference retains fifteen sites with glycans extending beyond a single GlcNAc; in that case, Endo H was added after Fab complex formation, potentially limiting enzyme access (*154*). The predominance of single-GlcNAc stubs in the Hi5-derived, Endo H–treated Env structure indicates efficient glycan trimming prior to complex formation, although some glycans may be incompletely modeled at this resolution. At the V3 glycan supersite, extended density was observed for N295, N332, and N386 in both structures, whereas N301 and N392 were modeled as single GlcNAc residues. At the V1/V2 apex, N133 appeared as a single GlcNAc in both structures, and reduced density at N156 and N160 in the trimmed structure was consistent with Endo H treatment. Additional glycan sites, including N197, N262, and N448, retained relatively extended density after Endo H treatment, consistent with limited accessibility. In contrast, sites surrounding the CD4bs, including N234, N276, N355, and N462, appeared to be efficiently trimmed to GlcNAc cores. At the 35O22–Env interface, N88 remained relatively ordered in both structures; N618 shifted from an extended glycan to a single GlcNAc; N625 was reduced from a minimal stub to no modelable density; and N637 remained a single GlcNAc in each structure. Importantly, core fucosylation was directly visualized at two sites (N462 and N618) in the untrimmed structure (**Fig. 2h**), consistent with the fucosylated mannose-type structures of Hi5 insect cell expression (**Fig. S1f-h**). Both sites rank among the most heavily fucosylated positions by LC-MS (**Fig. S1e-h**), providing independent structural support for the compositional assignment.

The two crystal structures provide the first structural view of HIV-1 Env trimers expressed in insect cells and their recognition by bNAbs in a distinct glycan context. Together with the nsEM analysis, these data confirm that the PBE/Hi5 system produces native-like, prefusion-closed Env immunogens comparable to those from mammalian systems. The preserved structural integrity, combined with well-defined glycan profiles and a core α(1,3)-fucose signature, supports further evaluation of Hi5-produced HIV-1 Env immunogens in animal models and translational studies.

### Murine NAb responses induced by Hi5-produced BG505 Env immunogens

In our previous studies, BG505 trimer-presenting SApNPs elicited tier 2 murine NAbs targeting the C3/V4 glycan patch (*35, 91*), and glycan trimming increased responder frequency (*65*). Here, we assessed the immunogenicity of Hi5-produced BG505 trimers and SApNPs, with or without glycan trimming, in mice (**Fig. 3a**). Immunogens were formulated with aluminum hydroxide (AH) alone or with the TLR9 agonist CpG ODN 1826 adsorbed to AH. Each dose contained 10 µg of antigen with 100 µL of AH or AH/CpG (200 µL total). CpG was selected based on its performance in our previous vaccine studies (*67, 71, 152*). Following our established protocol, groups of eight BALB/c mice received four intraperitoneal (i.p.) immunizations at weeks 0, 3, 6, and 9, with sera collected after each immunization (*65*). Mouse IgG was purified from week 11 sera for TZM-bl neutralization assays (**Fig. 3a-d**; **Figs. S3a-b**). The pseudovirus panel comprised the autologous tier 2 clade A BG505.T332N, tier 1 clade B SF162, and six heterologous tier 2 strains: clade A 398F1, clade B X2278, clade G X1632, clade C 25710, CRF07_BC BJOX2000, and CRF01_AE CNE8. This panel enabled rapid assessment of vaccine-induced cross-clade neutralization and NAb potency. Pseudotyped murine leukemia virus (MLV) served as a negative control. Notably, all pseudoviruses were produced in HEK293T cells and therefore carried mammalian glycans.

**Fig. 3.**
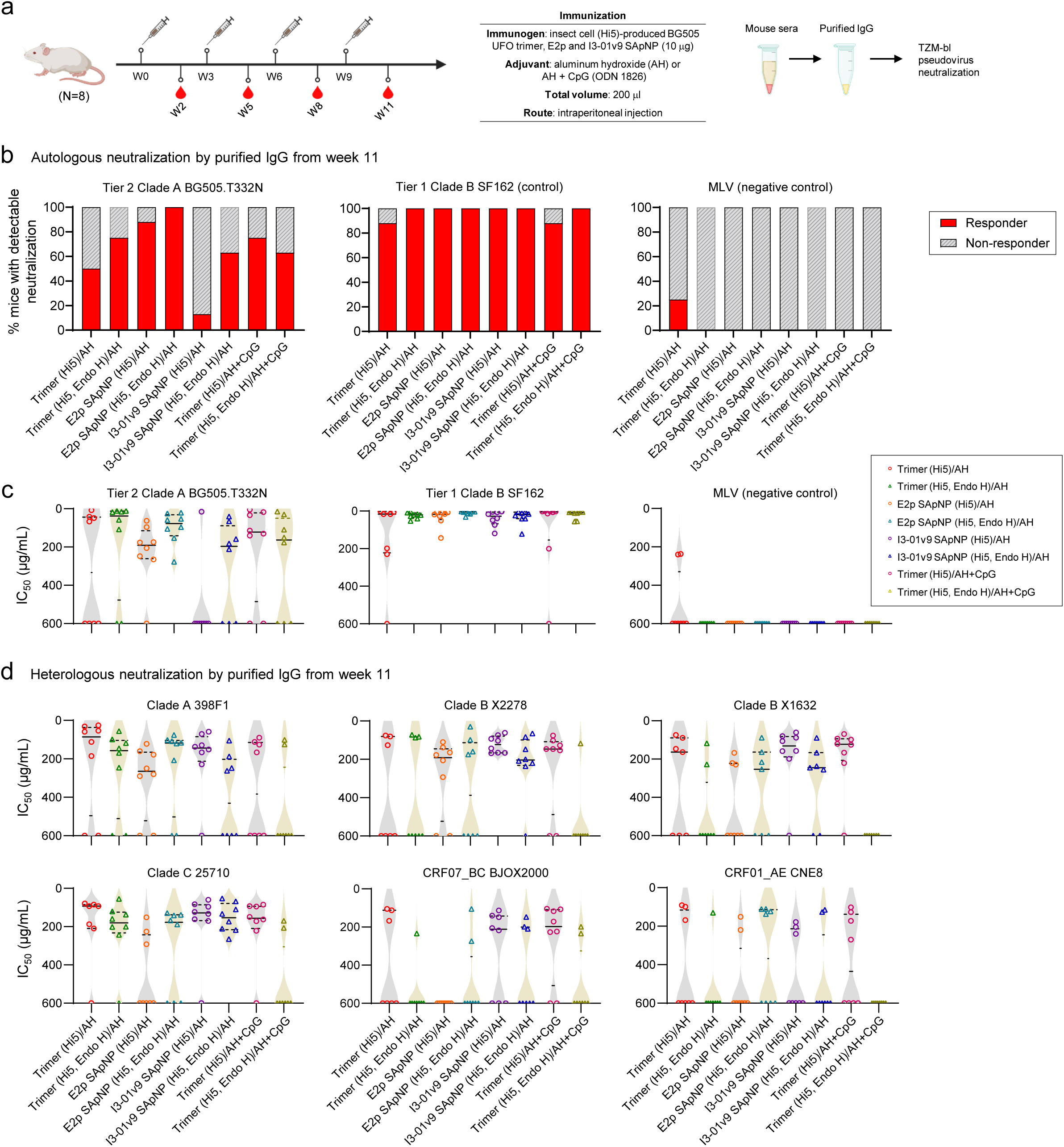
Cross-clade neutralizing responses elicited by Hi5-expressed BG505 Env immunogens in wild-type mice. (**a**) Schematic of the mouse immunization regimen for Hi5-expressed BG505 Env vaccines (n = 8 mice per group). Mice were immunized intraperitoneally (i.p.) at weeks 0, 3, 6, and 9 with 10 μg of protein antigen formulated in 200 μl of aluminum hydroxide (AH) alone or AH in combination with the TLR9 agonist CpG ODN 1826. A 3-week interval was used for rapid evaluation of vaccine formulations. Sera were collected after each immunization; IgG was purified from week-11 samples for TZM-bl neutralization assays. (**b**) Vaccine responder rates (RR) calculated for all groups using purified week-11 IgG against autologous tier 2 BG505.T332N (left), tier 1 clade B SF162 (middle), and pseudotyped murine leukemia virus (MLV) as a negative control (right). RR is defined as the percentage of animals per group with measurable half-maximal inhibitory dilution or concentration (ID_50_ or IC_50_) values against a given virus. (**c**) Neutralization potency (IC_50_) determined for all groups using purified week-11 IgG against the three viruses described in (b). (**d**) Vaccine-induced IgG neutralization against six heterologous tier 2 strains: 398F1 (clade A), X2278 (clade B), X1632 (clade G), 25710 (clade C), BJOX2000 (CRF07_BC), and CNE8 (CRF01_AE). IC_50_ values were calculated over a 0–100.0% neutralization range. The mouse immunization schematic was created with BioRender.com.

Using purified week 11 IgG, we calculated the autologous responder rate, defined as the proportion of mice with a measurable NAb response against BG505.T332N. Overall, all groups except the AH-formulated I3-01v9 SApNP group exhibited high responder rates (≥50%) against BG505.T332N. Glycan trimming substantially increased responder rates for the AH-adjuvanted Hi5-produced trimer, E2p SApNP, and I3-01v9 SApNP to 75%, 100%, and 63%, respectively, compared with their untrimmed counterparts (**Fig. 3b**, left). All groups elicited potent NAb responses against the tier 1 clade B SF162, except for one mouse in each untrimmed trimer group formulated with either AH alone or AH/CpG (**Fig. 3b**, middle). MLV reactivity was negligible, except for two samples in the AH-adjuvanted trimer group (**Fig. 3b**, right). Neutralization potency was quantified using the 50% inhibitory concentration (IC_50_) (**Fig. 3c**). Glycan trimming also markedly enhanced autologous neutralization potency in AH-adjuvanted groups. Among vaccine groups, the trimmed trimer group exhibited the highest potency, with a mean IC_50_ of 54.2 µg/mL (**Fig. 3c**, left). Most groups showed strong neutralization of tier 1 SF162, with mean IC_50_ values of 6.7–20.0 µg/mL (**Fig. 3c**, middle). MLV neutralization remained negligible (**Fig. 3c**, right).

We next assessed heterologous NAb responses against the 6-virus panel (**Fig. 3d**). Notably, measurable neutralization was observed across all six heterologous tier 2 isolates, indicating cross-clade activity in wild-type mice, although responses varied across strains and clades. Against clade A 398F1, the untrimmed trimer, untrimmed E2p SApNP, and trimmed E2p SApNP groups each showed a 75% responder rate. Among these, the untrimmed trimer elicited the highest potency (IC_50_ = 108.3 µg/mL), followed by the trimmed E2p SApNP (174.9 µg/mL). Against clade B X2278 and clade G X1632, the untrimmed I3-01v9 SApNP group showed the highest responder rates (100% and 88%, respectively) and the strongest neutralization, with IC_50_ values of 114.1 and 138.2 µg/mL. For clade C 25710, several groups—including the AH-adjuvanted trimmed trimer, untrimmed and trimmed I3-01v9 SApNPs, and AH/CpG-adjuvanted untrimmed trimer—showed 88% responder rates and comparable IC_50_ values (137.7–174.2 µg/mL). Against CRF07_BC BJOX2000, the AH/CpG-adjuvanted untrimmed trimer showed the highest responder rate (75%) and an IC_50_ of 211.9 µg/mL. Finally, against CRF01_AE CNE8, the AH-adjuvanted trimmed E2p SApNP group showed the highest responder rate (50%) and an IC_50_ of 267.9 µg/mL.

In summary, Hi5-produced BG505 Env immunogens elicited robust tier 2 NAb responses in mice, with or without glycan trimming. Two key observations emerged from these data. First, in a notable departure from CHO-derived counterparts (*65*), untrimmed Hi5-derived trimers induced robust autologous tier 2 NAbs. Second, the Hi5 immunogen groups collectively elicited measurable NAb responses against a panel of diverse heterologous tier 2 isolates. Together, these findings indicate that Hi5 production confers a distinct immunogenic signature on an Env that otherwise elicits limited tier 2 neutralization when produced in CHO cells (*65*). Immunogen format, glycan state, and adjuvant each modulated the magnitude and cross-reactivity of NAb responses.

### Glycoengineering a mammalian cell line to produce core α(1,3)-fucosylated HIV-1 Env

Mammalian- and Hi5-produced BG505 Env trimers share nearly identical protein backbones but differ in glycan composition, most notably in core α(1,3)-fucose, as well as in their ability to elicit cross-clade NAb responses. Therefore, we hypothesized that core α(1,3)-fucose enhances immune recognition of an otherwise evasive HIV-1 Env (**Fig. 4a**). For key neutralizing epitopes flanked by *N*-linked glycans, mammalian cells generate self-like glycans that may hide these epitopes from immune recognition and promote immune evasion, whereas Hi5 cells produce glycans containing core α(1,3)-fucose that may serve as target cues for adjacent epitopes, thereby enhancing antibody responses to them. To test this hypothesis, we supplemented cultures with fucose and co-expressed exogenous core α(1,3)-fucosyltransferases (FucTs) from plants (*Arabidopsis thaliana*, AT) and insects (*Bombyx mori*, BM; *Drosophila melanogaster*, DM) during ExpiCHO expression, creating a mammalian platform for producing core α(1,3)-fucosylated immunogens (**Fig. 4b**).

**Fig. 4.**
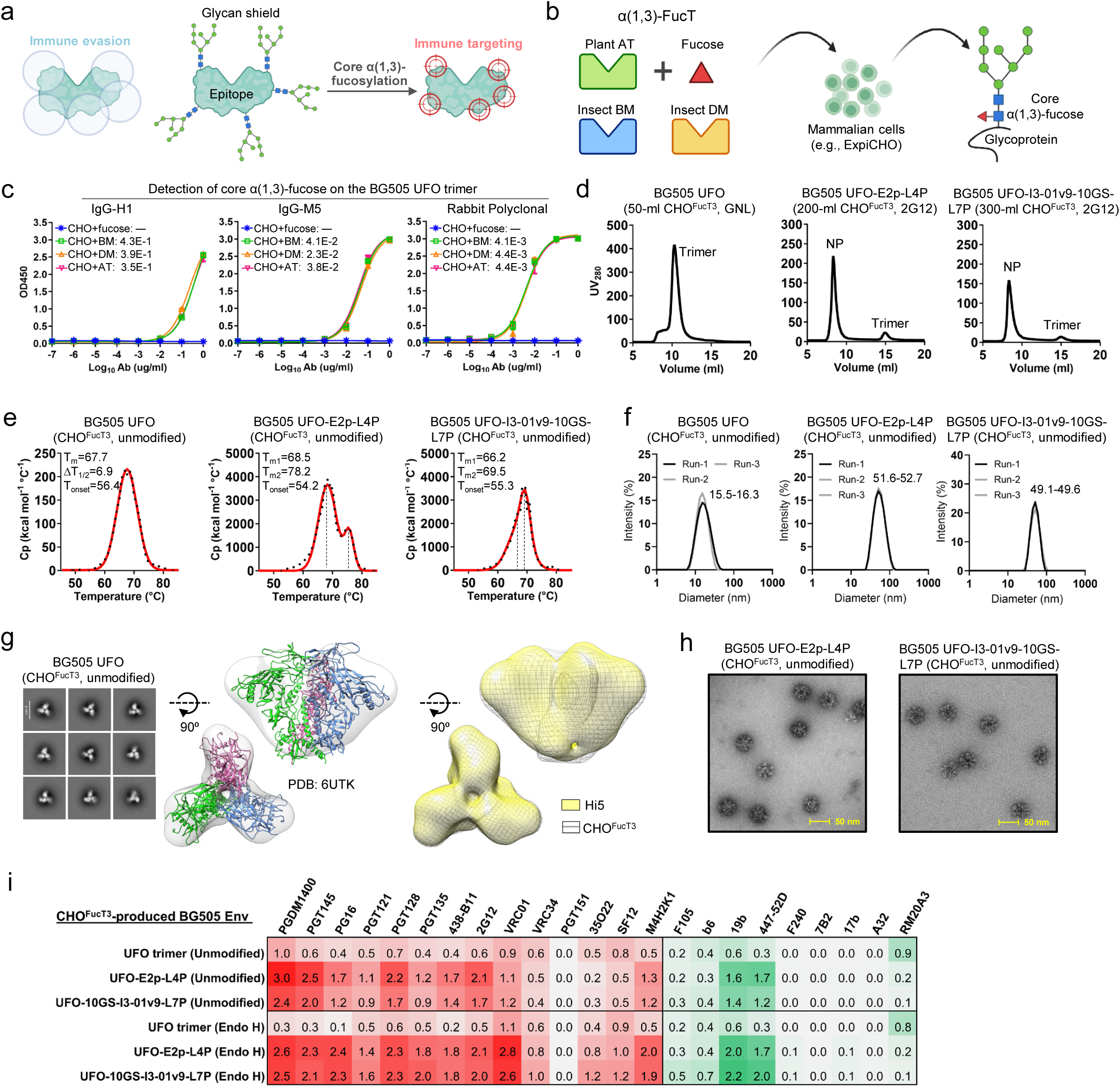
Glycoengineered mammalian expression and in vitro characterization of core α(1,3)-fucosylated BG505 Env immunogens. (**a**) Schematic of the core α(1,3)-fucose hypothesis: while mammalian-expressed “self-glycans” are immune-evasive, core α(1,3)-fucosylation of N-linked glycans surrounding a bNAb epitope may serve as “target cues” that help direct immune responses toward the epitope. (**b**) Schematic of the glycoengineering strategy. L-fucose and one or more core α(1,3)-fucosyltransferases from plants (*Arabidopsis thaliana*, AT) and insects (*Bombyx mori*, BM; *Drosophila melanogaster*, DM) are incorporated into a mammalian expression system to generate HIV-1 Env immunogens bearing mammalian-type glycans with core α(1,3)-fucose. (**c**) ELISA curves of glycoengineered ExpiCHO-expressed BG505 UFO trimers binding to core α(1,3)-fucose-specific monoclonal antibodies (IgG-H1 and IgG-M5) and a rabbit polyclonal antibody. Four conditions were tested: fucose alone or with individual core α(1,3)-fucosyltransferase (AT, BM, or DM). EC_50_ values are indicated. The system containing fucose plus all three core α(1,3)-fucosyltransferases was termed CHO^FucT3^. (**d**) SEC profiles of CHO^FucT3^-expressed, unmodified BG505 UFO trimer (left), UFO-E2p-L4P SApNP (middle), and UFO-10GS-I3-01v9-L7P SApNP (right). Trimers were purified by GNL chromatography, and SApNPs by IAC using a 2G12 column. (**e**) DSC thermograms of CHO^FucT3^-expressed, unmodified BG505 Env immunogens. Experimental data and Gaussian fits are shown, with thermal parameters indicated. (**f**) Particle size distributions of CHO^FucT3^-expressed, unmodified BG505 Env immunogens from three independent batches. Hydrodynamic diameter (Dh) ranges are indicated. (**g**) nsEM analysis of CHO^FucT3^-expressed, unmodified BG505 UFO trimer. Left: representative 2D class averages. Middle: side and top views of the 3D reconstruction with BG505 UFO (PDB: 6UTK) fitted into the density. Right: comparison of 3D reconstructions from Hi5 and CHO^FucT3^ systems. (**h**) Representative nsEM images of CHO^FucT3^-expressed, unmodified BG505 UFO trimer-presenting E2p and I3-01v9 SApNPs. (**i**) Antigenic profiles of CHO^FucT3^-expressed BG505 UFO trimer and SApNPs (E2p and I3-01v9) in unmodified or glycan-trimmed forms. Fourteen bNAbs/NAbs and nine non-NAbs were evaluated by BLI on an Octet RED96 using six antigen concentrations (starting at 266 nM for the trimer and 5.6 nM for SApNPs, followed by 2-fold serial dilutions; sensorgrams in **Figs. S4g–l**). Peak responses at the highest concentration are shown in a color-coded matrix (green to red). The schematics in (a) and (b) were created with BioRender.com.

We first expressed the BG505 UFO trimer in 25-mL ExpiCHO cultures supplemented with fucose, with each of the three FucTs co-expressed individually. All four conditions yielded nearly identical SEC profiles, with a trimer peak at ∼10 mL, indicating minimal effects on Env folding (**Fig. S4a**). BN-PAGE analysis of SEC fractions showed a single band, further confirming trimer integrity (**Fig. S4b**). ELISA was performed using IgG-H1, IgG-M5 (*132*), and RabbitPoly as antibody probes (**Fig. 4c**). BG505 trimers co-expressed with each FucT in the presence of fucose, but not those expressed with fucose alone, yielded EC_50_ values of 0.35–0.43 μg/mL (H1), 0.023– 0.041 μg/mL (M5), and 0.0041–0.0044 μg/mL (RabbitPoly), confirming incorporation of core α(1,3)-fucose into ExpiCHO-produced glycans. We then expressed BG505 immunogens in 50-mL (trimer), 200-mL (E2p), and 300-mL (I3-01v9) ExpiCHO cultures supplemented with fucose and co-transfected with AT, BM, and DM FucT plasmids at a 1:1:1 ratio. This glycoengineered expression system was designated CHO^FucT3^. The BG505 UFO trimer exhibited an SEC profile comparable to that observed with fucose alone or with fucose and individual FucTs, whereas the two 60-mer SApNPs produced well-defined particle peaks at ∼8.5 mL with a minor tailing peak at 14.8–15 mL corresponding to unassembled subunits (**Fig. 4d**). CHO^FucT3^-produced BG505 Env immunogens exhibited thermal profiles largely similar to their Hi5-produced counterparts (**Fig. 4e**). Compared with standard ExpiCHO expression (*65*), the trimer and I3-01v9 SApNP showed slight decreases in T_m_ and T_onset_, whereas the E2p SApNP exhibited slightly higher T_m1_/T_m2_ values, with all differences within 1 °C. DLS confirmed that core α(1,3)-fucosylation had minimal impact on size distribution for both SApNPs, with the BG505 UFO trimer displaying a Dh of ∼16.0 nm (**Fig. 4f**). We next examined whether the core α(1,3)-fucosylated glycan shield was susceptible to Endo H treatment. BN-PAGE showed a clear downward shift for the trimer band after treatment, whereas both SApNPs, regardless of glycan state, remained in the loading well (**Fig. S4c**). The trimer and SApNPs exhibited slightly reduced Dh values after trimming (**Fig. S4d**).

Given that core α(1,3)-fucosylation had no adverse effect on Env biophysical properties, we evaluated the structural integrity and antigenicity of CHO^FucT3^-produced immunogens prior to *in vivo* studies. BG505 UFO trimers bearing core α(1,3)-fucosylated mammalian glycans were characterized by nsEM (**Fig. 4g**). The 2D class averages displayed a propeller-like morphology typical of native-like Env trimers, and rigid-body fitting of the BG505 UFO trimer structure (PDB: 6UTK) into the 3D density map showed excellent agreement, supporting structural integrity (**Fig. 4g**, left). The 3D reconstructions of CHO^FucT3^- and Hi5-produced trimers overlapped closely, indicating a native-like Env conformation across the two expression systems and glycan contexts (**Fig. 4g**, right). The native-like Env architecture was preserved after Endo H treatment, as shown by nsEM 2D and 3D analyses (**Fig. S4e**). The CHO^FucT3^-produced SApNPs yielded well-formed particles with visible surface projections, and this morphology remained unaffected by Endo H treatment (**Fig. 4h**; **Fig. S4f**). Antigenicity was assessed by BLI using the same panel of antibodies (**Fig. 4i**; **Figs. S4g–l**). Overall, CHO^FucT3^-produced immunogens exhibited antigenic profiles broadly similar to their non-fucosylated counterparts, although their binding signals were slightly reduced for most bNAbs and NAbs. The effect of glycan trimming was consistent with findings in our previous ExpiCHO study (*65*): although trimming reduced bNAb recognition of the V2 apex, it led to a more balanced antigenic profile for SApNPs, with strong binding to bNAbs targeting the V2 apex, N332-glycan supersite, and CD4bs. Together, these data demonstrate that core α(1,3)-fucosylation and enzymatic trimming can be integrated into a mammalian expression platform to generate structurally intact, antigenically competent HIV-1 Env immunogens.

Our glycoengineering strategy successfully incorporated core α(1,3)-fucose into the glycan shield of HIV-1 Env expressed in ExpiCHO cells. By introducing this non-mammalian motif into a system that normally produces immunologically “silent” self-glycans, we created a controlled platform to evaluate whether this glycan modification enhances NAb elicitation. This platform not only provides a direct experimental test of our hypothesis but also offers an engineered mammalian expression system for producing Env immunogens with defined xenoglycan modifications.

### Murine NAb responses induced by CHO^FucT3^-produced BG505 Env immunogens

We assessed the immunogenicity of CHO^FucT3^-produced BG505 UFO trimers and SApNPs, along with an ExpiCHO-derived BG505 UFO trimer, in BALB/c mice (**Fig. 5a**). The immunization regimen followed our previous protocol (*65*). Each dose contained 10 µg of antigen formulated with 100 µL of AH in a final volume of 200 µL. Sera were collected two weeks after each vaccine dose. Env-specific binding antibody (bAb) responses were evaluated throughout the immunization course, whereas vaccine-induced NAb responses were assessed at the final time point (week 11).

**Fig. 5.**
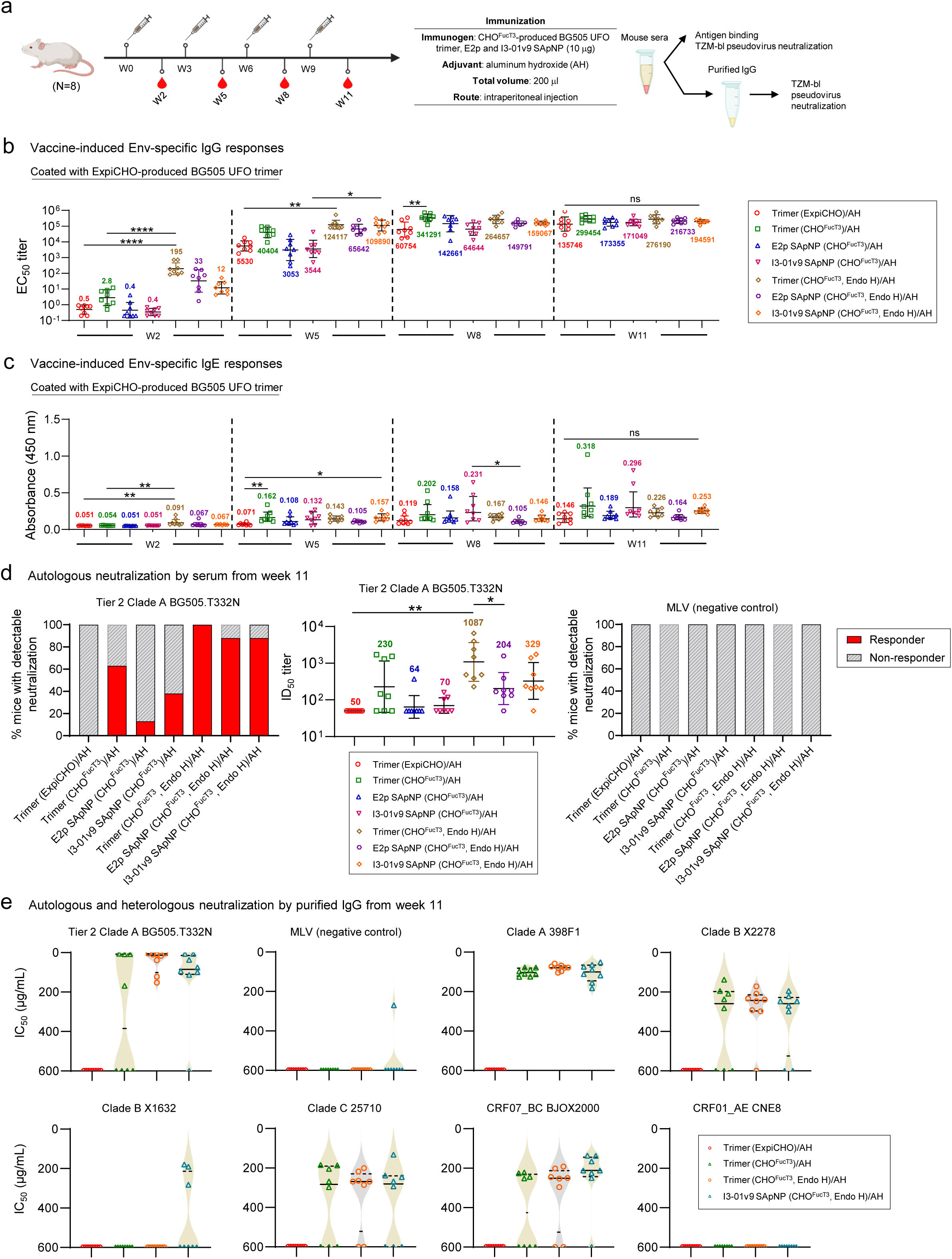
Binding and neutralizing antibody responses elicited by CHO^FucT3^-expressed BG505 Env immunogens in wild-type mice. (**a**) Schematic of the mouse immunization regimen for CHO^FucT3^-expressed BG505 Env vaccines (n = 8 mice per group). Mice were immunized intraperitoneally (i.p.) at weeks 0, 3, 6, and 9 with 10 μg of protein antigen formulated in 200 μl of aluminum hydroxide (AH). Sera were collected after each immunization for binding and TZM-bl neutralization assays. For selected groups, IgG was purified from week-11 samples for TZM-bl neutralization assays. (**b**–**c**) Longitudinal Env-specific IgG (**b**) and IgE (**c**) responses at weeks 2, 5, 8, and 11 during immunization, using the ExpiCHO-produced BG505 UFO trimer as the coating antigen in serum ELISA. EC_50_ titers are plotted for IgG, and absorbance at 450 nm (A_450_) values are plotted for IgE. Data points are shown as geometric mean ± geometric SD. Statistical analysis was performed using one-way ANOVA followed by Tukey’s multiple-comparisons post hoc test at each time point. Statistical significance is indicated as follows: ns (not significant), \**p* < 0.05, \*\**p* < 0.01, and \*\*\*\**p* < 0.0001. (**d**) Autologous neutralization by week 11 serum. Left: vaccine responder rates (RR) against autologous tier 2 BG505.T332N. Middle: NAb responses (ID_50_ titers) against autologous tier 2 BG505.T332N. Right: RR measured against MLV (negative control). (**e**) Autologous and heterologous neutralization by purified week 11 IgG. In addition to autologous tier 2 BG505.T332N and the negative control MLV, six heterologous tier 2 strains were tested: 398F1 (clade A), X2278 (clade B), X1632 (clade G), 25710 (clade C), BJOX2000 (CRF07_BC), and CNE8 (CRF01_AE). IC_50_ values were calculated over a 0–100.0% neutralization range. The mouse immunization schematic was created with BioRender.com.

Serum IgG and IgE responses were measured by ELISA against an ExpiCHO-produced BG505 UFO trimer (**Fig. 5b-c**; **Figs. S5a-c**). The CHO^FucT3^-produced, glycan-trimmed trimer elicited the highest IgG titers at weeks 2 and 5, with EC_50_ values 390- and 22.4-fold higher than those of the ExpiCHO trimer and 69.6- and 3.1-fold higher than those of the untrimmed CHO^FucT3^ trimer, respectively (**Fig. 5b**). At weeks 8 and 11, the glycan-trimmed CHO^FucT3^ trimer maintained the highest IgG titers, with EC_50_ values 5.6- and 2.2-fold higher than those of the ExpiCHO trimer. For core α(1,3)-fucosylated SApNPs, glycan trimming consistently enhanced EC_50_ titers at all time points. At week 5, Endo H-treated E2p and I3-01v9 SApNPs achieved EC_50_ titers of 65,642 and 109,890, 21.5- and 31-fold higher, respectively, than their untrimmed counterparts. We also examined IgE responses previously associated with core α(1,3)-fucose in insect and plant allergens (*127, 132, 158*), as such responses may affect vaccine-induced immunity. Both CHO^FucT3^ trimer groups exhibited higher IgE absorbance (A_450_) than the ExpiCHO trimer across all time points (**Fig. 5c**). However, the difference in IgE levels between the untrimmed and trimmed groups diminished after two vaccine doses. Among the CHO^FucT3^-produced SApNPs, I3-01v9 elicited the highest IgE responses at weeks 8 and 11, with A_450_ values of 0.231 and 0.296, 1.9- and 2.0-fold higher, respectively, than the ExpiCHO trimer. After four doses, IgE levels remained several orders of magnitude lower than the corresponding IgG levels, with an estimated IgG-to-IgE ratio exceeding 6,700:1, indicating a strong bias toward IgG, with relatively low IgE induction.

We evaluated serum neutralization using week 11 samples (**Fig. 5d**; **Fig. S3d**). Responder rates and 50% inhibitory dilution (ID_50_) titers were determined against autologous BG505.T332N. Overall, CHO^FucT3^ trimers and SApNPs elicited robust NAb responses, whereas the ExpiCHO trimer showed no detectable neutralization. Glycan trimming substantially increased responder rates for the CHO^FucT3^ trimer (100%), E2p SApNP (88%), and I3-01v9 SApNP (88%) relative to their untrimmed counterparts (**Fig. 5d**, left). Notably, the glycan-trimmed CHO^FucT3^ trimer elicited the strongest NAb response, with an ID_50_ titer of 1,087, 4.7-fold higher than that of the untrimmed trimer (**Fig. 5d**, middle). Similarly, the glycan-trimmed CHO^FucT3^ E2p and I3-01v9 SApNPs yielded ID_50_ titers of 204 and 329, representing 3.1- and 4.7-fold increases over their untrimmed counterparts, respectively. MLV reactivity was negligible in endpoint sera (**Fig. 5d**, right). To confirm these results, IgG was purified from week 11 sera of four groups—the ExpiCHO trimer, untrimmed and glycan-trimmed CHO^FucT3^ trimers, and glycan-trimmed CHO^FucT3^ I3-01v9 SApNP—and tested against autologous BG505.T332N, six heterologous strains, and MLV (**Fig. 5e**; **Figs. S3e-f**). Against BG505.T332N, both CHO^FucT3^ trimer groups showed responder rates of ≥50%, whereas the ExpiCHO trimer showed no detectable neutralization. Consistent with the serum data, glycan trimming increased the trimer responder rate from 50% to 100%, while the glycan-trimmed I3-01v9 SApNP achieved an 88% responder rate. Neutralization potency followed a similar trend, with the glycan-trimmed CHO^FucT3^ trimer exhibiting the greatest potency (IC_50_ = 19.3 µg/mL). Nonspecific MLV reactivity was detected in only one animal in the I3-01v9 SApNP group. Cross-clade neutralization was then assessed against six heterologous tier 2 pseudoviruses. The ExpiCHO trimer showed no cross-neutralization, whereas all CHO^FucT3^ groups exhibited measurable activity against multiple strains. Against clade A 398F1, all CHO^FucT3^ groups showed a 100% responder rate with IC_50_ values of 78.3–100 µg/mL. Against clade B X2278, responder rates ranged from 63 to 88%, with IC_50_ values of 266.5–307.3 µg/mL. Against clade G X1632, only the CHO^FucT3^ I3-01v9 SApNP group showed neutralization (38% responder rate; IC_50_ = 406.6 µg/mL). Against clade C 25710 and CRF07_BC BJOX2000, CHO^FucT3^ groups showed responder rates of 50–88% with IC_50_ values of 215.4–375.9 µg/mL. Unlike the Hi5 immunogens (**Fig. 3d**), no neutralization was observed against CRF01_AE CNE8.

These findings demonstrate that core α(1,3)-fucosylation enhances Env-specific NAb responses, while immunogen format (trimer *vs.* SApNP) and glycan state (unmodified *vs.* Endo H-trimmed) further modulate the magnitude and cross-reactivity of neutralization in wild-type mice. Consistent with our prior observations for Hi5 immunogens, CRF01_AE CNE8 remained the most resistant strain in the panel, whereas heterologous responses were more frequently observed against clade A, B, and C isolates and CRF07_BC.

### Responses induced by core α(1,3)-fucosylated Env within a human antibody repertoire

To evaluate the impact of core α(1,3)-fucosylation on HIV-1 Env vaccine immunogenicity within a human antibody repertoire, we assessed selected BG505 UFO trimers in Alloy transgenic mouse models (*159–161*). Equipped with human immunoglobulin loci, these mice provide a relevant platform for characterizing human antibody responses to glycoengineered Env immunogens. This is particularly important because core α(1,3)-fucose is a non-human xenoglycan motif that may differentially influence B-cell selection and maturation in human versus murine repertoires.

Here, we evaluated core α(1,3)-fucosylated BG505 UFO trimers in Alloy ATX-GK and ATX-GKH mice, with the latter exhibiting a hyperimmune phenotype. The Hi5 trimer was tested in ATX-GK mice (Group 1) and ATX-GKH mice (Group 3), whereas the glycan-trimmed CHO^FucT3^ trimer was tested in ATX-GKH mice (Group 2). Each group comprised two Alloy mice and followed the same immunization and sampling protocol used for wild-type mice (**Fig. 6a**). To maximize the likelihood of generating Env-specific human NAb responses in this limited cohort, mice were immunized with 50 μg of trimer formulated with an AH/STING agonist mixture (**Fig. 6a**). Env-specific serum IgG and IgE responses were quantified by ELISA against an ExpiCHO-produced BG505 UFO trimer. Given the small sample size (*n* = 2 per group), data are shown as individual mouse values with group means indicated (**Fig. 6b**; **Figs. S6**). Overall, Group 1 reached peak IgG titers at week 11, whereas both ATX-GKH cohorts (Groups 2 and 3) plateaued by week 8. Group 2 exhibited the highest IgG titers throughout the study, with mean titers 962- and 144-fold higher than those of Groups 1 and 3 at week 2; 105- and 2-fold higher at week 5; 8- and 3-fold higher at week 8; and 4.4- and 2.8-fold higher at week 11, respectively. Notably, both Group 2 animals showed detectable IgG titers after a single dose, compared with titers at or near the limit of detection in the other two groups. This pattern was unlikely to result solely from the ATX-GKH phenotype, as Group 3 shared the same genetic background as Group 2. Instead, the combination of core α(1,3)-fucosylation and glycan trimming was associated with a more rapid IgG response, consistent with early B-cell activation. Group 2 achieved a peak mean EC_50_ titer of 2,220,303 at week 8, ∼20-fold higher than that observed in wild-type mice immunized with the same construct. Serum IgE responses were assessed using the same ExpiCHO trimer probe to measure Env-specific, rather than xenoglycan-specific, reactivity (**Fig. 6c, Fig. S6**). Across both mouse strains and all three immunogens, IgE signals remained low throughout immunization and were generally lower than those observed in wild-type mice (**Fig. 5c**). At week 11, Group 2 exhibited a modest mean A_450_ of 0.124, whereas one Group 3 animal reached an A_450_ of 1.013. IgE responses remained several orders of magnitude lower than the corresponding IgG responses, yielding high IgG-to-IgE ratios throughout the study. These data suggest a strong bias toward Env-specific IgG responses, with minimal IgE-associated reactivity in Alloy mice.

**Fig. 6.**
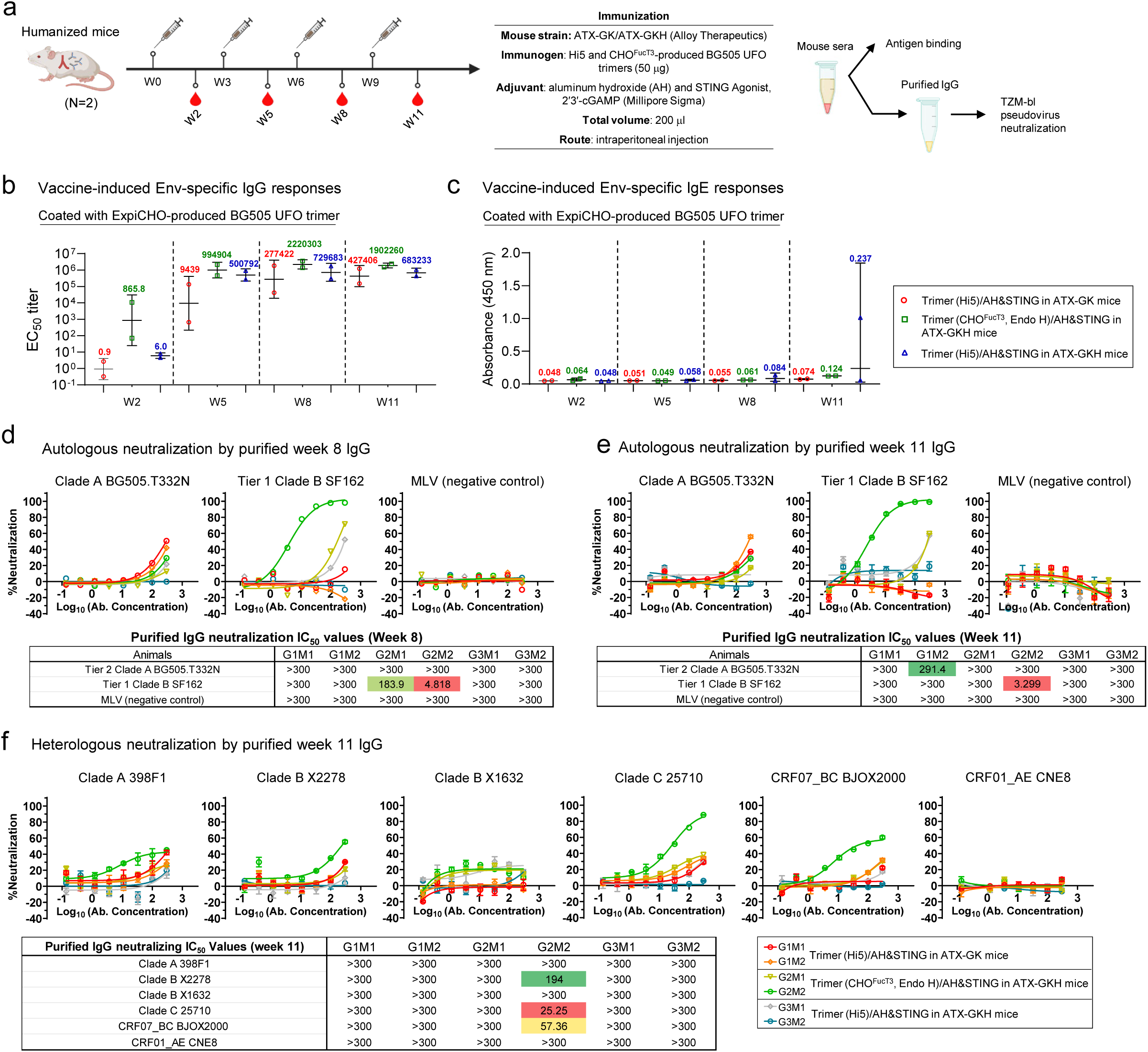
Antibody responses elicited by core α(1,3)-fucosylated BG505 UFO trimers in Alloy humanized mice. (**a**) Schematic of the humanized mouse immunization regimen for core α(1,3)-fucosylated BG505 UFO trimers (n = 2 mice per group). Two ATX-GK and four ATX-GKH (Hyperimmune) mice were immunized intraperitoneally (i.p.) at weeks 0, 3, 6, and 9 with 50 μg of protein antigen formulated in 100 μl of aluminum hydroxide (AH) and a STING agonist. Sera were collected after each immunization for binding and TZM-bl neutralization assays. IgG was purified from week-8 and week-11 sera from all six mice for TZM-bl neutralization assays. (**b–c**) Longitudinal Env-specific IgG (**b**) and IgE (**c**) responses at weeks 2, 5, 8, and 11 measured by serum ELISA using ExpiCHO-produced BG505 UFO trimer as the coating antigen. EC_50_ titers are plotted for IgG, and absorbance at 450 nm (A450) values for IgE. Data points are shown as geometric mean ± geometric SD. (**d–e**) Autologous neutralization by purified week-8 (**d**) and week-11 (**e**) IgG. Top: percent neutralization curves against autologous tier 2 BG505.T332N, tier 1 clade B SF162, and MLV as a negative control. Bottom: IC_50_ values against these three viruses. (**f**) Heterologous neutralization by purified week-11 IgG against six tier 2 isolates from the global panel. Top: percent neutralization curves against clade A 398F1, clade B X2278, clade G X1632, clade C 25710, CRF07_BC BJOX2000, and CRF01_AE CNE8. Bottom: IC_50_ values against these six viruses. In (**d–f**), IC_50_ values are color-coded based on neutralization potency.

To assess vaccine-induced NAb responses, we purified IgG from week 8 and 11 sera of all six Alloy mice. Purified IgG samples were evaluated in TZM-bl assays against autologous tier 2 BG505.T332N, tier 1 clade B SF162, and MLV. Overall, week 8 (**Fig. 6d**) and week 11 (**Fig. 6e**) samples showed similar neutralization profiles, with limited autologous activity, variable tier 1 responses, and no nonspecific MLV signals. The strongest tier 1 response was observed for mouse 2 in Group 2, with IC_50_ values of 4.8 and 3.3 μg/mL against SF162 at weeks 8 and 11, respectively. Autologous BG505.T332N neutralization was limited, with only mouse 2 in Group 1 reaching a detectable IC_50_ of 291.4 μg/mL at week 11. To assess cross-clade neutralization, we next tested a panel of six heterologous tier 2 pseudoviruses spanning clades A, B, C, B/C, and A/E (**Fig. 6f**). Whereas most samples showed little or no heterologous activity, mouse 2 in Group 2 exhibited detectable cross-clade neutralization against three tier 2 viruses: clade B X2278, clade C 25710, and CRF07_BC BJOX2000, with the greatest potency against the clade C isolate, 25710 (IC_50_ = 25.3 μg/mL). Notably, the same mouse lacked detectable autologous neutralization against BG505.T332N. CRF01_AE CNE8 was also not neutralized, consistent with the absence of a response to this strain in wild-type mice immunized with CHO^FucT3^ trimers and SApNPs (**Fig. 5e**). Given the small cohort, these cross-clade neutralizing responses are encouraging but remain observational and warrant cautious interpretation.

In summary, our findings indicate that core α(1,3)-fucosylated trimers elicited robust Env-specific IgG responses with minimal IgE-associated reactivity, and the highest IgG titers were observed in the glycan-trimmed CHO^FucT3^ trimer group. The neutralization data suggest that the combination of core α(1,3)-fucosylation and glycan trimming may elicit detectable cross-clade NAb responses within a human antibody repertoire following a short immunization regimen. We therefore next examined whether the nascent NAb responses arose from antibody lineages related to known bNAb pathways, targeted established sites of Env vulnerability, or represented distinct modes of recognition influenced by core α(1,3)-fucosylation.

### Single-cell analysis of human NAb responses induced by core α(1,3)-fucosylated Env

To investigate the molecular basis of early NAb responses, we performed antigen-specific single B-cell sorting on splenocytes from Group 2 mouse 2 (G2M2), which exhibited detectable cross-clade neutralization, to isolate Env-specific B cells for human monoclonal antibody (mAb) cloning (**Fig. 7a**). For sorting, we used a biotinylated, Avi-tagged BG505 gp140 trimer probe containing an optimized HR1_N_ fragment (*34, 35*) to capture B cells recognizing diverse Env epitopes. This probe was previously used to identify PGT121 intermediates from a phage-displayed Donor 517 antibody library (*151*) and to isolate vaccine-induced NAbs from mice and rabbits immunized with BG505 Env trimers and SApNPs (*91*). Trimer-binding B cells were isolated by fluorescence-activated cell sorting (FACS), enabling recovery of paired heavy- and light-chain sequences from individual cells. From a single sorting plate, we cloned 55 antibodies and recovered 18 paired full-length heavy-chain (HC) and κ-light-chain (KC) sequences (**Figs. S7a,b**).

**Fig. 7.**
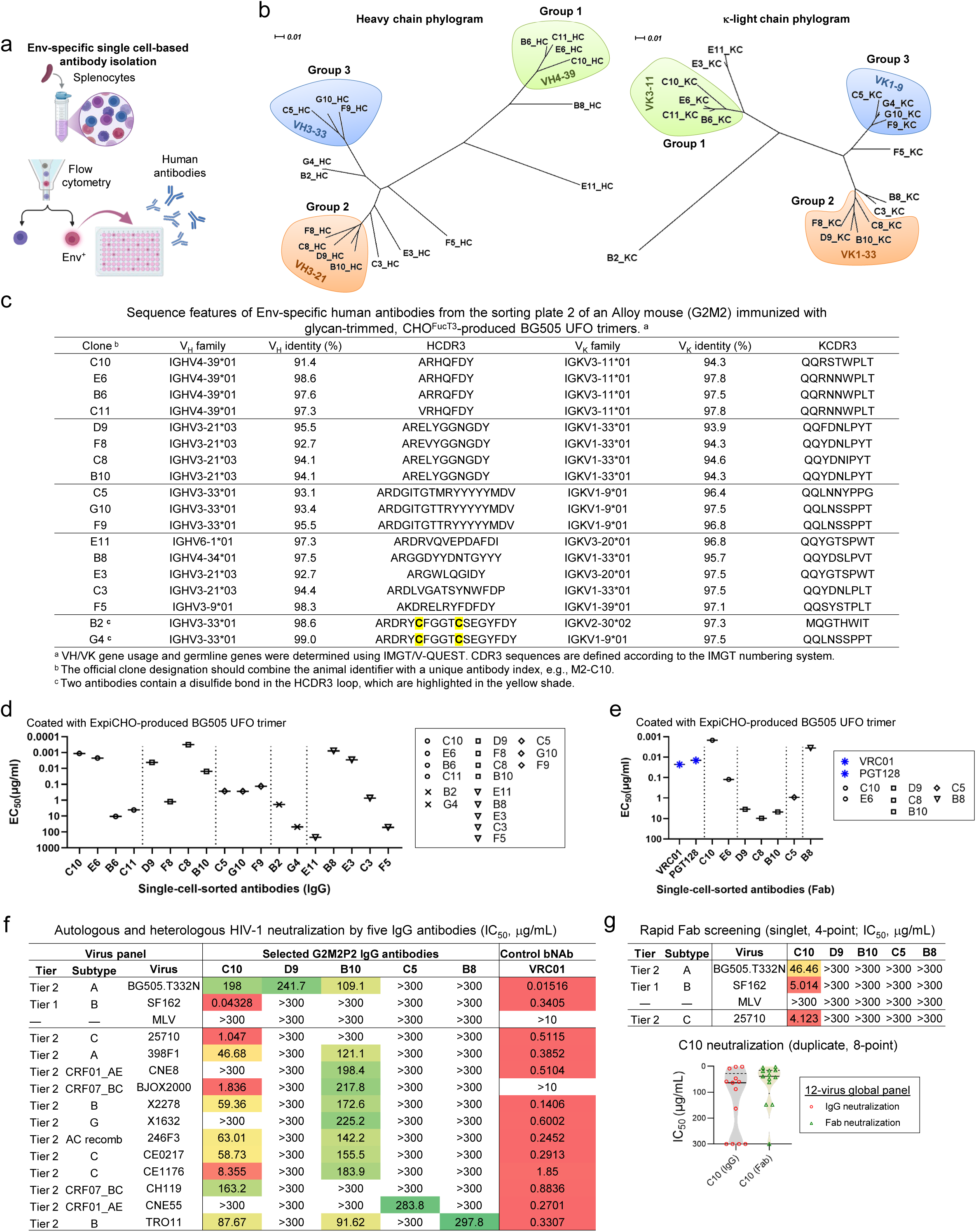
Diverse binding and neutralization profiles of human antibodies isolated by Env-specific single-cell sorting. (**a**) Schematic workflow of splenocyte isolation, Env-specific single B-cell sorting, and antibody cloning. (**b**) Heavy- and light-chain phylograms of 18 selected human antibodies for experimental validation. Three groups are color-coded based on shared germline V genes and heavy- and light-chain pairing: group 1 (green, VH4-39/Vκ3-11), group 2 (orange, VH3-21/Vκ1-33), and group 3 (blue, VH3-33/Vκ1-9). (**c**) Sequence features of 18 selected human antibodies, including germline V gene family, V gene identity, and CDR3 sequences. (**d**) ELISA binding (EC_50_) of 18 human antibodies as IgGs to ExpiCHO-produced BG505 UFO trimer. (**e**) ELISA binding (EC_50_) of seven human antibodies as Fabs to ExpiCHO-produced BG505 UFO trimer, with Fabs of VRC01 and PGT128 included for comparison. (**f**) Autologous and heterologous HIV-1 neutralization by five IgG antibodies, C10, D9, B10, C5, and B8 (IC_50_, μg/mL). Autologous tier 2 BG505.T332N, tier 1 clade B SF162, and MLV were tested initially, with MLV serving as a negative control. Twelve heterologous tier 2 isolates in the global panel were tested to assess neutralization breadth, with VRC01 IgG included as a bNAb control. (**g**) Top: estimated IC_50_ values from rapid Fab screening performed in singlet with a four-point dilution series; bottom: comparison of C10 IgG and Fab neutralization against the 12-virus global panel. In (**f–g**), IC_50_ values are color-coded according to neutralization potency.

Genetic analysis by IMGT/V-QUEST (*162*) revealed diverse germline gene usage, with multiple clonal lineages arising from distinct human V-gene families (**Fig. 7b**). These mAbs were predominantly derived from the V_H_3/V_H_4 HC and Vκ1/Vκ3 KC families, with a single V_H_6-derived HC and one Vκ2-derived KC. This distribution was reflected in the unrooted HC and KC phylograms, which clustered sequences by their germline gene usage: Group 1 (V_H_4-39*01/Vκ3-11*01), Group 2 (V_H_3-21*01/Vκ1-33*01), and Group 3 (V_H_3-33*01/Vκ1-9*01) (**Fig. 7b**). V_H_ and Vκ sequence identities relative to germline were 91.4–99.0% and 93.9–97.8%, respectively, indicating low levels of somatic hypermutation (SHM), consistent with the rapid immunization regimen (**Fig. 7c**). HCDR3 analysis revealed several notable features associated with human bNAbs (**Fig. 7c**). In particular, one mAb lineage contained five consecutive tyrosine residues (YYYYY) within an 18-amino-acid HCDR3 loop. Long, tyrosine-rich HCDR3s are often observed in human bNAbs targeting glycan-shielded epitopes on native Env, including V2 apex-directed bNAbs such as PG9 (*79*), PG16 (*163, 164*), and CAP256-VRC26 (*165*), as well as the N332 supersite bNAb BG18 (*166*). These features may facilitate penetration of the dense Env glycan shield and access to underlying protein surfaces. Similar YYYY motifs are also found in gp120/gp41 interface bNAbs, including PGT151 and ACS202 (*167, 168*), where tyrosine-rich HCDR3s engage the flexible gp41 fusion peptide and form an intermolecular antiparallel β-sheet. Two mAbs contained 18-amino-acid HCDR3s with predicted intra-HCDR3 disulfide bonds (C-X4-C). Such constrained conformations may reduce the entropic penalty of binding and facilitate access to sterically restricted epitopes, as observed for the CD4bs and N332 supersite recognized by VRC08 (*169*) and 438-B11 (*154*), respectively. Evidence of clonal expansion and somatic variation was also observed in lineages with short HCDR3s of 7 and 11 amino acids (**Fig. 7c**), corresponding to Groups 1 and 2, respectively (**Fig. 7b**). These findings indicate that core α(1,3)-fucosylated, glycan-trimmed Env elicited diverse human antibody lineages, including antibodies with sequence characteristics observed in known human bNAbs.

These antibodies bound the ExpiCHO-produced BG505 UFO trimer with a wide range of apparent binding strengths as measured by ELISA. In the IgG format (**Fig. 7d**; **Fig. S7c**), C10 and E6 were the strongest binders in Group 1 (EC_50_ = 0.0011–0.0022 μg/mL), whereas C8 from Group 2 exhibited the strongest Env binding overall, with an EC_50_ of 0.00032 μg/mL, 13- and 48-fold stronger than D9 and B10, respectively. In contrast, C5 and F9 bound Env weakly, with EC_50_ values of 0.13–0.27 μg/mL. Seven antibodies were evaluated as Fabs, along with bNAb controls VRC01 and PGT128 (**Fig. 7e**; **Fig. S7d**). C10 and B8 Fabs retained strong Env binding, with EC_50_ values of 0.0015 μg/mL and 0.0036 μg/mL, respectively, up to 10-fold lower than those of the two bNAb controls (0.014–0.023 μg/mL). In contrast, D9 and B10 were highly sensitive to IgG-to-Fab conversion, with a ∼three-order-of-magnitude reduction in binding. Five IgGs, C10, D9, B10, C5, and B8, were tested by BLI using AHC analytical sensors and ExpiCHO-produced BG505 UFO trimer as the analyte (**Fig. S7e**). C10 and B8 exhibited the strongest binding signals with minimal dissociation, whereas C5 showed substantially weaker binding, and B10 and D9 exhibited clear dissociation. These five IgGs were tested in TZM-bl assays against autologous BG505.T332N, tier 1 clade B SF162, 12 heterologous tier 2 isolates, and MLV (**Fig. 7f**; **Fig. S7f**). Although C10 showed weak autologous neutralization, it potently neutralized tier 1 SF162 (IC_50_ = 0.043 μg/mL) and neutralized 9 of the 12 heterologous tier 2 isolates, with IC_50_ values ranging from 1.05 to 163.2 μg/mL. Thus, C10 displayed substantial cross-clade breadth despite weak autologous activity. B10 exhibited weaker cross-clade neutralizing activity, with IC_50_ values of 91.6–225.2 μg/mL. D9, a member of the B10 clonal lineage, as well as C5 and B8, displayed negligible neutralizing activity. As expected, bNAb VRC01 exhibited exceptional potency and breadth, except against CRF07_BC BJOX2000, consistent with previous reports (*170*). In a rapid screening assay against three HIV-1 isolates, all five antibodies except C10 lost neutralizing activity upon conversion to Fab (**Fig. 7g**, top; **Fig. S7g**). A more detailed assay revealed that the C10 Fab retained neutralizing activity and, against several isolates in the 12-virus panel, showed enhanced neutralization relative to IgG, suggesting effective Env recognition in a monovalent form (**Fig. 7g**, bottom; **Fig. S7h**).

In summary, single-cell analysis identified multiple Env-specific human antibodies with diverse germline gene usage and sequence features associated with known bNAbs. Functional validation identified C10, a human NAb of V_H_4-39 and Vκ3-11 germline origins that neutralized 75% of tier 2 isolates in the global panel and retained neutralizing activity as a Fab. These findings provide a basis for epitope mapping and B-cell repertoire analysis of selected antibodies.

### Epitope probing, repertoire profiling, and lineage analysis of selected human antibodies

Given the diverse neutralization profiles of C10, D9, B10, C5, and B8, we sought to gain structural insight into their Env recognition and trace their evolution within the B-cell repertoire. To this end, Fabs of selected antibodies were complexed with different Env trimers for nsEM-based epitope mapping. Because Alloy-derived antibodies may approach Env differently from known NAbs and bNAbs or recognize epitopes influenced by core α(1,3)-fucosylation, their binding modes may not readily match established structural references. To facilitate epitope assignment, we first generated a panel of simulated low-resolution (30 Å) density maps for bNAbs targeting the V2 apex (PG9, PG16 (*139*), PGT145 (*138*)), the N332 supersite (PGT121, PGT124, PGT128, PGT135 (*138*)), the CD4bs (VRC01 (*146*), 8ANC134 (*171*), HJ16 (*172*)), the FP (VRC34 (*173*), ACS202 (*174*)), the SF (SF12 (*141*), VRC-PG05 (*142*)), and the gp120/gp41 interface (35O22 (*149*), 3BC315 (*175*)), along with the base-directed non-NAb RM20A3 (*136*) (**Fig. S8a**). These maps served as references for the nsEM analysis of Alloy-derived human antibodies.

For C10, the ExpiCHO-produced BG505 UFO trimer was complexed with Fab overnight, followed by SEC purification for nsEM analysis. 2D class averages revealed classes containing two or three apparent Fab densities per trimer (**Fig. 8a**, panel #1; **Fig. S8b**, left). C10 appeared to approach Env laterally in the 2D views. The characteristic triangular shape of the native-like Env trimer remained visible, although the propeller tips appeared blunted and density around the putative interaction site was poorly resolved, consistent with conformational and/or Fab flexibility. In the 3D reconstruction generated without C3 symmetry, Fab density was less well resolved, limiting reliable assignment of binding stoichiometry and orientation, as reflected by comparison with simulated maps of VRC01-, VRC34-, and SF12-Env complexes (**Fig. S8b**, right and bottom). Together, these low-resolution data indicate heterogeneous C10–Env recognition but do not permit reliable assignment of its epitope or binding mode. For B10, given its weak binding to unmodified Env, we used the immunization antigen, the CHO^FucT3^-produced, glycan-trimmed BG505 UFO trimer, for complex formation and nsEM analysis (**Fig. 8a**, panel #2; **Fig. S8c**). A characteristic Fab-like shape with a “hole” was clearly visible in some 2D classes, potentially representing top views of the B10-Env complex (**Fig. 8a**, panel #2). Despite additional nsEM analyses and 3D reconstructions, conformational heterogeneity and limited resolution precluded reliable epitope assignment for this antibody as well (**Fig. S8c**). C5 was analyzed similarly but yielded separate 2D classes of Env trimers and Fabs (**Fig. 8a**, panel #3), consistent with its weak binding (**Fig. 7d-e**; **Fig. S7e**). B8 appeared to dissociate Env trimers during overnight incubation and was therefore complexed for only 1 h prior to nsEM analysis (**Fig. 8a**, panel #4). 2D class averages revealed a clear binding mode, with one Fab bound to the trimer base in most classes and two Fabs in occasional classes. Fitting the BG505 UFO Env/RM20A3 model (PDB: 6UTK, 6NF5) into the 3D reconstruction supported assignment of B8 as a base-directed antibody with an angle of approach resembling RM20A3. Higher-resolution structures by crystallography or cryo-EM will be needed to clarify how these antibodies recognize their epitopes on native-like Env and how their binding modes relate to their distinct neutralizing profiles.

**Fig. 8.**
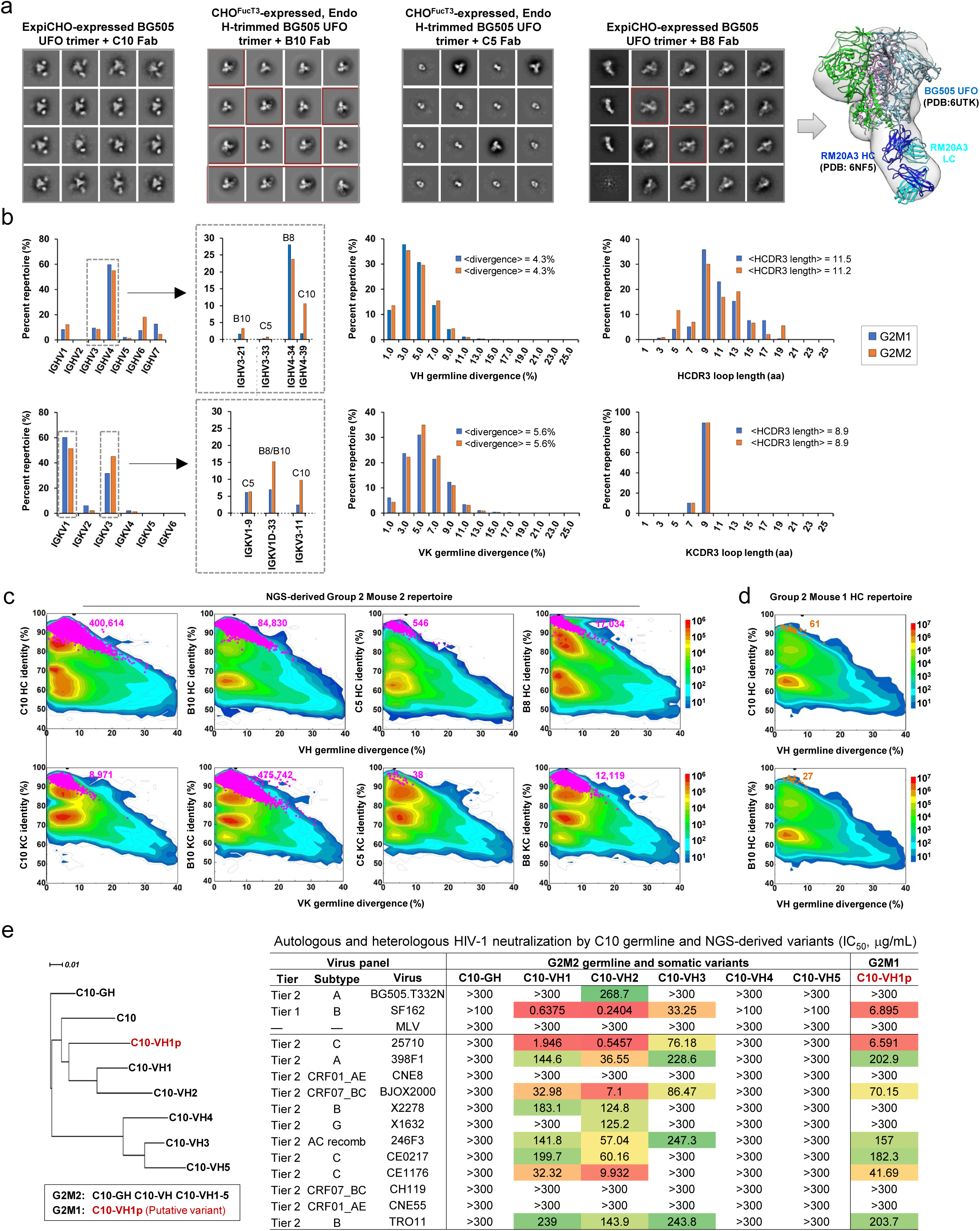
Epitope mapping and repertoire-level lineage analysis of select human antibodies elicited by α(1,3)-fucosylated HIV-1 Env. (**a**) Epitope mapping of C10, B10, C5, and B8 by negative-stain EM analysis, with representative 2D class averages shown for BG505 UFO trimer, either unmodified or glycan-trimmed, in complex with each Fab. For B10 and B8, complexes with two Fabs bound are highlighted with dark red boxes. For B8, a 3D reconstruction fitted with structures of the BG505 UFO trimer (PDB: 6UTK) and RM20A3 (PDB: 6NF5) is shown because of the unambiguous epitope assignment. (**b**) Quantitative antibody repertoire profiles of G2M1 and G2M2 showing germline gene usage (left; inset showing subfamily-level usage of germline genes associated with the four antibodies of interest), SHM (germline divergence, middle), and HCDR3/KCDR3 loop lengths (right). (**c**) Divergence-identity analysis of C10, B10, C5, and B8 lineages within their respective G2M2 heavy-chain (top) and light-chain (bottom) repertoires. Heavy- and κ-light-chain sequences are plotted as a function of sequence identity to the template and divergence from their putative germline genes. Color-coding denotes sequence density. Templates and putative lineage variants identified based on shared germline V genes and a CDR3 identity cutoff of 95% are shown as black and pink dots, respectively, with the corresponding sequence numbers indicated. (**d**) Identification of C10- and B10-like lineage sequences within the G2M1 heavy-chain repertoires. Templates and putative lineage variants are shown as black and orange dots, respectively, with the corresponding sequence numbers indicated. (**e**) Functional validation of putative heavy-chain germline precursors and lineage variants spanning different degrees of SHM for C10. Left: germline-rooted phylogenetic tree of C10 heavy-chain variants identified in G2M2 and in a second animal, G2M1. Right: neutralization (IC_50_, μg/mL) against the autologous tier 2 clade A virus BG505.T332N, tier 1 clade B virus SF162, the 12-virus global panel, and MLV as a negative control. C10-GH: inferred germline precursor; C10-VH1-5: G2M2-derived somatic variants; C10-VH1p: putative C10 variant identified in the G2M1 repertoire. A cutoff of 300 μg/mL was used for all tier 2 isolates, whereas 100 μg/mL was used for tier 1 SF162.

Repertoire profiling and lineage analysis were performed on splenic B cells from the two mice (G2M1 and G2M2) immunized with glycan-trimmed CHO^FucT3^ trimer. HC and KC variable domain libraries were sequenced using the Aviti platform, and raw data were processed using a modified Antibodyomics pipeline (*154, 176–178*) (**Fig. S8d**). Aviti sequencing generated ∼8.1– 12.2 million reads per library, yielding 7.2–9.6 million high-quality, error-free antibody chains.

The two mice exhibited similar germline gene usage: the IGHV4 family predominated among V_H_ genes (55.1-59.8%), while IGKV1 and IGKV3 accounted for 51.4-60.1% and 31.7-45.1%, respectively, of the Vκ repertoire (**Fig. 8b**, left). Despite this overall similarity, genes associated with specific NAb lineages were selectively enriched in G2M2 (**Fig. 8b**, left; **Fig. S8f**). IGHV4-39, the C10 HC germline gene, accounted for 10.6% of the G2M2 HC repertoire compared with only 1.8% in G2M1. Similarly, IGKV3-11, the C10 KC germline gene, constituted 9.9% of the repertoire in G2M2 versus 2.5% in G2M1. By contrast, somatic hypermutation (SHM) and CDR3 length distributions were nearly identical between the two mice. HCs were less mutated than KCs overall (**Fig. 8b**, middle), and the average HCDR3 length (11.5 aa) was shorter than that previously reported for healthy human donors (12.7-13.0 aa) (*179*) (**Fig. 8b**, right). These shared characteristics likely reflect similar baseline repertoires and responses to the same immunogen and regimen. We next examined the prevalence of these antibody lineages within the G2M2 B-cell repertoire. Using a CDR3 identity cutoff of 95%, putative somatic variants of the HC and KC for each antibody were identified within their respective germline gene families (**Fig. S8g**, left) and visualized on the divergence-identity plots (**Fig. 8c**). Despite variations in lineage size, expanded lineages spanning a broad range of SHM were identified for all four antibodies. Notably, C10 and B10 HCs accounted for 39.3% and 26.6% of the IGHV4-39 and IGHV3-21 germline gene families, respectively, indicating substantial expansion of these two NAb lineages. Finally, we investigated whether G2M1, the Alloy mouse lacking detectable cross-clade serum neutralization, contained similar antibody lineages. Querying the G2M1 HC and KC repertoires using germline V-gene and H/KCDR3 criteria revealed a small population of putative somatic variants for C10 and B10 (**Fig. 8d**; **Fig. S8g**, right), suggesting that related antibody lineages may have been initiated in G2M1 but remained at low abundance. However, because HC and KC repertoires were independently analyzed, the putative HC and KC variants could not be confirmed as paired chains from the same B cells. Even if correctly paired, whether their limited expansion accounts for the absence of detectable cross-clade serum neutralization in G2M1 remains uncertain.

Finally, we characterized the functional properties of putative C10 and B10 HC variants identified in the G2M2 and G2M1 repertoires, along with their inferred precursors (**Fig. S8h**). Twelve HC sequences were selected based on germline divergence and paired with the parental LCs for expression and functional testing. Although some reconstituted IgGs were expressed at low yields, both inferred HC germline precursors and several somatic variants were tested for neutralization (**Fig. 8e**; **Fig. S8i-j**). As expected, neither germline precursor showed detectable neutralizing activity. Among the five C10 variants from G2M2, C10-VH1, C10-VH2, and C10-VH3, carrying 6, 11, and 18 more V-gene nucleotide mutations than C10, respectively, neutralized 8, 9, and 5 of 12 global-panel isolates, with IC_50_ values as low as 1.95, 0.55, and 76.18 µg/mL. By contrast, the other two variants with more V-gene mutations showed no detectable neutralization. Because all HC variants were paired with the parental LC, the observed bell-shaped relationship between HC divergence and neutralizing activity should be interpreted with caution, as highly diverged HCs may not pair well with a less mature LC, resulting in functional loss. Notably, a putative C10-related HC variant from G2M1, C10-VH1p, exhibited a similar heterologous neutralization profile, neutralizing 7 of 12 isolates with an IC_50_ as low as 6.6 µg/mL. C10-VH1, C10-VH2, and C10-VH1p all clustered with parental C10 within the same phylogenetic branch (**Fig. 8e**, left), consistent with the presence of a putatively related HC lineage in a second animal, albeit with more limited expansion than in G2M2. For B10, two variants identified from G2M2, B10-VH6 and B10-VH8, exhibited weak autologous neutralization (**Fig. S8i-j**).

Our analyses revealed distinct features of human NAbs elicited by core α(1,3)-fucosylated Env. nsEM suggested heterogeneous Env recognition by C10 and B10, whereas B8 targeted the trimer base. Repertoire profiling showed expansion and diversification of C10 and B10 lineages in G2M2, with putatively related variants detected in G2M1. Multiple C10 variants retained cross-clade neutralization, including a putative C10 HC variant from G2M1, supporting the presence of putatively related HC variants across independently immunized animals.

## DISCUSSION

The glycan shield, once viewed as a passive mask that conceals conserved protein epitopes (*13*), is now considered an integral part of HIV-1 Env structure and immune recognition (*72, 180*). The discovery of glycan-reactive bNAbs has contributed to this shift in vaccine design by revealing that the glycan shield itself contains conserved, targetable glycopeptide epitopes. Contemporary vaccine strategies therefore aim not simply to evade the glycan shield, but to engage and modulate it to guide protective immune responses (*72–76*).

While the preservation of authentic glycans has been an important principle in HIV-1 Env vaccine design (*76, 94, 181*), various strategies have been evaluated to improve NAb elicitation (*182*). Among these strategies, glycan masking (*183, 184*) and immunofocusing (*185*) seek to redirect antibody responses through selective glycan modification. Adding specific glycans can mask immunodominant decoy regions from antibody recognition, whereas deleting others can enhance exposure of targeted bNAb epitopes, such as the CD4bs (*96, 97*) and the V2 apex (*186*). Deleted glycans can subsequently be reintroduced through sequential immunogens to restore the wild-type glycan shield (*96, 97*). The dense glycan canopy can also pose a substantial barrier to naïve bNAb precursors, which often have low initial affinity for heavily glycosylated epitopes and may require extensive affinity maturation to accommodate the native glycan shield or establish specific interactions with glycans within it. Germline-targeting strategies are designed to activate defined B-cell precursors and guide their maturation toward bNAbs (*187, 188*), and have been applied to the CD4bs (*189–192*), the N332 supersite (*193–196*), and the V2 apex (*197–200*). In sequential immunization regimens, priming immunogens present a simplified glycan landscape to activate rare precursor B cells, followed by booster immunogens with progressively restored native glycan complexity to guide maturation toward bNAbs. This stepwise approach aims to recapitulate key features of virus-antibody co-evolution observed in bNAb donors (*201, 202*).

Glycan trimming provides a direct approach to reducing the steric barrier imposed by the Env glycan shield. Conversion of oligomannose glycans to single GlcNAc residues using Endo H enhanced epitope accessibility while preserving native-like Env folding and trimer structure. This strategy builds on previous influenza HA and SARS-CoV-2 spike vaccine studies (203, 204) and has recently been extended to HIV-1 Env vaccines (65). Despite the reduction in glycan mass, glycan-trimmed BG505 UFO trimers displayed on SApNPs retained binding to V2- and V3-glycan-dependent bNAbs, suggesting that GlcNAc remnants, when presented in a multivalent format, can maintain key glycan–protein interactions involved in Env recognition. The tier 2 NAb responses elicited by the trimmed trimer further suggest that an intact glycan shield is not required for tier 2 NAb induction. Mechanistically, glycan trimming may balance structural integrity with epitope accessibility. By reducing bulky peripheral oligomannose glycans while preserving key epitope-associated glycans, such as N332 at the supersite, this approach alleviates steric hindrance without disrupting the Env antigenic landscape. However, the immunological effects of glycan trimming appear to be viral antigen-dependent. In our recent studies, oligomannose enrichment, rather than trimming, improved NAb elicitation for filovirus GP and soluble HCV E1E2 (*66, 152*), indicating distinct effects of glycan remodeling across viral glycoproteins.

Beyond enzymatic trimming, host-cell glycosylation pathways may be manipulated to alter the immunological properties of the glycan shield. Hi5 cells produce paucimannose-rich glycans with limited terminal processing, thereby reducing glycan shield complexity while preserving a native-like Env conformation (*205, 206*). A distinctive feature of Hi5-derived glycoproteins is the presence of core α(1,3)-fucose. Unlike mammalian core α(1,6)-fucose, core α(1,3)-fucose is a non-mammalian carbohydrate motif recognized by human antibodies, including IgE, and can engage allergy-associated immune pathways (*126–129*). We hypothesized that core α(1,3)-fucose could modulate immune responses when incorporated into vaccine antigens. Indeed, Hi5-produced, core α(1,3)-fucosylated BG505 Env immunogens elicited cross-clade NAb responses in mice. We then engineered the CHO^FucT3^ system to incorporate core α(1,3)-fucose into mammalian glycans. When carrying unmodified mammalian glycans, native-like BG505 trimers failed to elicit tier 2 NAb responses in mice (*207*), whereas CHO^FucT3^-produced immunogens elicited robust autologous and heterologous murine NAb responses. Notably, the glycan-trimmed CHO^FucT3^ trimer also induced detectable cross-clade neutralization in Alloy humanized mice. Although the precise mechanism remains unclear, engagement of IgE-associated immune pathways may have contributed to these responses. Core α(1,3)-fucosylated Env did not produce observable adverse reactions in animals, and IgE levels remained low relative to Env-specific IgG. Despite its low serum abundance, IgE can engage Fcε receptor-bearing immune cells and influence downstream immune responses (*208–210*). Conceptually, core α(1,3)-fucosylation enables antigen-encoded immunostimulation, with core α(1,3)-fucose acting as a covalently embedded signal for localized activation. By enhancing recognition of adjacent protein and glycopeptide epitopes, such xenoglycan motifs may reshape antibody responses to the Env glycan shield and overcome its barriers to NAb elicitation.

The Alloy mouse findings raise the possibility that glycan reprogramming may provide an alternative route to human NAb responses with cross-clade breadth. Germline-targeting strategies (*187*) are designed to activate rare bNAb precursors and guide their maturation through sequential immunogens that progressively restore the native glycan shield (*95–98*). In contrast, four doses of a core α(1,3)-fucosylated, glycan-trimmed trimer elicited multiple human antibody lineages within an 11-week regimen in Alloy mice. Single-cell analysis identified mAbs with diverse sequence features, while B-cell repertoire sequencing revealed variants across expanded lineages. Notably, C10 showed moderate neutralization with substantial cross-clade breadth, and multiple C10 variants retained similar breadth and potency. A putative C10 HC variant identified in a second animal also exhibited cross-clade neutralization when paired with the parental C10 KC, suggesting that related antibody responses may have arisen in multiple animals within the same group. These findings suggest that xenoglycosylation may support the development of multiple human antibody lineages with cross-neutralizing activity, rather than relying on the maturation of a predefined lineage through multistage immunization designed to recapitulate prolonged bNAb development pathways (*211, 212*). Although the Alloy study is exploratory and requires further validation, cross-clade neutralization was observed in two distinct lineages, suggesting glycan reprogramming may provide an alternative strategy for broadening human antibody responses to Env.

An important unresolved question is how human NAbs elicited by core α(1,3)-fucosylated Env recognize the native glycan shield. Initial nsEM analyses of several antibody–Env complexes showed conformational and binding heterogeneity that limited reliable epitope assignment. Further epitope mapping, together with higher-resolution structural studies where feasible, will be needed to determine whether these responses converge on known bNAb epitopes, access established sites of vulnerability through alternative binding modes, or recognize other features of Env structure. Such studies may also clarify how core α(1,3)-fucose and glycan trimming alter antibody access to Env and shape the development and breadth of NAb lineages. In particular, defining the precise epitopes and binding modes of C10 and B10 will require additional structural and functional investigation. Some of these Alloy-derived human NAbs may recognize Env through structurally flexible or noncanonical binding modes distinct from those of well-characterized bNAbs isolated from human donors. It remains possible that NAb elicitation in a humanized mouse model differs from that in natural human infection. More broadly, the strategy described here may represent an example of AllergoVaccinology—an intersection of allergology and vaccinology in which allergy-associated immune recognition mechanisms are repurposed to enhance vaccine-induced protective antibody responses against pathogens. Incorporation of allergic xenoglycan motifs, such as α(1,3)-fucose and β(1,2)-xylose (*125*), or other non-self sugars (*213*), into glycoproteins extends vaccine design beyond antigen stabilization and nanoparticle display to include glycan composition as an additional variable in immunogen design across diverse viral glycoproteins.

## METHODS

### Expression and purification of BG505 Env immunogens

For transient expression of BG505 Env immunogens in the BTI-Tn-5B1-4 (High Five™, or “Hi5”) cell line derived from the cabbage looper *Trichoplusia ni* (Expression Systems; cat. no. 94-002S), a plasmid expression vector was constructed as previously described (*116, 117*). The vector backbone was obtained from Addgene (plasmid #139772), and the Orgyia pseudotsugata immediate early 2 (OpIE2) promoter was amplified from an Addgene template (plasmid #50559) by polymerase chain reaction (PCR). The baculoviral immediate early 1 terminator (IE1Ter) was incorporated into the vector (*116, 117*), which was further optimized for protein secretion and yield. Genes encoding the BG505 UFO trimer, UFO-E2p-LD4-PADRE SApNP, and UFO-10GS-I3-01v9-LD7-PADRE SApNP were excised from their corresponding mammalian vectors and ligated into the digested Hi5 vector backbone. All expression plasmids were sequence-verified. Hi5 cells were thawed and cultured in ESF921 medium (Expression Systems; cat. no. 96-001-01) at 27°C and 135 rpm in a shaker incubator. When cells reached 2.0 × 10^6^ cells/mL, cultures were concentrated to 4.0 × 10^6^ cells/mL and transfected with polyethyleneimine (PEI-MAX; Polysciences, Inc.). For a 100-mL culture, 400 μg of plasmid in 5 mL ESF921 medium was mixed with 3.2 mL PEI-MAX (1.0 mg/mL) in 1.8 mL ESF921 medium. The DNA–PEI-MAX complex was then added to 100 mL of Hi5 cells. Five hours post-transfection, cultures were diluted with ESF921 medium to 1.0 × 10^6^ cells/mL in a final volume of 400 mL. Two days later, cultures were further diluted to 1.0–2.0 × 10^6^ cells/mL by adding 400 mL of ESF921 medium. On day 5, culture supernatants were harvested, clarified by centrifugation at 4,000 rpm for 10 min, and filtered through a 0.45 μm membrane (Thermo Scientific).

To incorporate core α(1,3)-fucose into mammalian-expressed BG505 Env immunogens, three core α(1,3)-fucosyltransferases (FucTs) were co-expressed individually or in combination in ExpiCHO™ cells (Thermo Fisher; cat. no. A29133) supplemented with L(-)-fucose (Chem-Impex; cat. no. 00736). Plasmids encoding full-length FucTs were generated from the plant *Arabidopsis thaliana* (accession no. BAB02969) and the insects *Drosophila melanogaster* (AJ302045) and *Bombyx mori* (AB551654). Previously described ExpiCHO™ expression protocols (*65*) were adapted for glycoengineering. ExpiCHO™ cells were cultured in ExpiCHO^TM^ Expression Medium (Thermo Fisher) at 37°C, 135 rpm, and 8% CO_2_. At 10 × 10^6^ cells/mL, cultures were diluted with fresh medium to 6 × 10^6^ cells/mL before transfection. ExpiFectamine^TM^ CHO/DNA complexes were prepared according to the manufacturer’s instructions. L(-)-fucose was supplemented to 5 mM on day 0 before transfection and every other day thereafter until harvest. For initial screening, BG505 UFO was expressed with L(-)-fucose alone or with each FucT individually in the presence of L(-)-fucose. For CHO^FucT3^ production, the three FucT plasmids were co-expressed at a 1:1:1 ratio. For a 100-mL transfection, 90 μg of antigen plasmid, 23 μg of one FucT plasmid or 23 μg of each of the three FucT plasmids (69 μg total), and 320 μl of ExpiFectamine™ CHO reagent were mixed in 7.7 mL of cold OptiPRO™ medium. After the first feed on day 1, cells were cultured at 32°C, 120 rpm, and 8% CO_2_ following the Max Titer protocol, with an additional feed on day 5. Culture supernatants were harvested on days 13–14, clarified by centrifugation at 1,467 × g for 20 min, and filtered through a 0.45 μm membrane (Thermo Fisher).

The same purification protocols were used for each immunogen format regardless of the expression system used. BG505 UFO trimers were purified from culture supernatants using a *Galanthus nivalis* lectin (GNL) column (Vector Labs), whereas SApNPs were purified by immunoaffinity chromatography (IAC) using a 2G12 antibody column. Trimers and SApNPs were further purified by size-exclusion chromatography (SEC) on a Superdex 200 Increase 10/300 GL column and a Superose 6 Increase 10/300 GL column, respectively. Protein concentrations were determined by UV absorbance at 280 nm (UV_280_) using theoretical extinction coefficients.

### Glycan trimming by Endo H treatment and enzyme removal

The protocol for Endo H treatment and removal was described previously (*65*). Briefly, N-linked glycans of purified BG505 UFO trimers and SApNPs were trimmed using endoglycosidase H (Endo Hf), a fusion of Endo H and maltose-binding protein (MBP) (NEB; cat. no. P0703L). In each reaction, 1 mg of antigen was mixed with 200 μl of 10× GlycoBuffer 3, 250 μl of Endo Hf, and H_2_O as needed to a final volume of 2 ml. The mixture was incubated at room temperature (25°C) for 5 h to allow enzymatic digestion of Env glycans. After incubation, reaction mixtures were purified by SEC using a Superdex 200 Increase 10/300 GL column (for trimers) or a Superose 6 Increase 10/300 GL column (for SApNPs) to remove the MBP-tagged enzyme. In our previous study (*65*), amylose resin (NEB; cat. no. E8021S) was used to remove residual Endo Hf following SEC purification. In the present study, this additional step was omitted to maximize antigen recovery for mouse immunization. SEC-purified fractions were pooled, aliquoted, and stored in liquid nitrogen or at −80°C until use.

### SDS-PAGE and BN-PAGE

To assess molecular weight and integrity, Hi5- and ExpiCHO-produced BG505 Env immunogens were analyzed by sodium dodecyl sulfate–polyacrylamide gel electrophoresis (SDS-PAGE) and blue native polyacrylamide gel electrophoresis (BN-PAGE). For SDS-PAGE under reducing conditions, proteins were mixed with 6× Laemmli SDS sample buffer (Thermo Scientific), boiled for 5 min at 100°C, and loaded onto a 10% Tris-glycine gel (Bio-Rad). Gels were run at 250 V for 25 min using SDS running buffer (Bio-Rad) and stained with InstantBlue (Abcam). For BN-PAGE, proteins were mixed with 4× native loading dye and loaded onto a 4–12% Bis-Tris NativePAGE gel (Life Technologies). Gels were run at 150 V for 2–2.5 h using NativePAGE running buffer (Life Technologies). BN-PAGE gels were stained with Coomassie Brilliant Blue R-250 (Bio-Rad) and destained with 6% ethanol and 3% glacial acetic acid.

### Differential scanning calorimetry (DSC)

Melting temperatures (T_m_ or T_m1_/T_m2_) and associated parameters were measured for BG505 Env immunogens produced in the Hi5 and glycoengineered CHO^FucT3^ expression systems and purified using GNL affinity chromatography for trimers or 2G12 immunoaffinity chromatography (IAC) for SApNPs, followed by SEC. DSC was performed using a MicroCal PEAQ-DSC instrument (Malvern). Purified proteins in PBS were diluted to 0.1–5 μM. Thermal denaturation was assessed from 20 to 100°C at a scan rate of 60°C/h. Data processing, including buffer correction, normalization, and baseline subtraction, was performed using MicroCal PEAQ-DSC software. Gaussian fitting was carried out in GraphPad Prism (version 10.3.1).

### Dynamic Light Scattering (DLS)

Particle size distributions of BG505 UFO trimers and SApNPs were measured using a Zetasizer Ultra instrument (Malvern). Purified BG505 Env immunogens produced in the Hi5 and ExpiCHO expression systems were diluted to 0.2 mg/mL in PBS, and 30 μL of each sample was loaded into a quartz batch cuvette (Malvern, cat. no. ZEN2112). Measurements were performed at 25°C in backscatter mode. Data were processed using Zetasizer Ultra software. Particle size distributions were plotted in GraphPad Prism (version 10.3.1).

### Site-specific glycan analysis

100 µg aliquots of each sample were denatured for 1h in 50 mM Tris/HCl, pH 8.0 containing 6 M of urea and 5 mM dithiothreitol (DTT). Next, Env samples were reduced and alkylated by adding 20 mM iodoacetamide (IAA) and incubated for 1h in the dark, followed by a 1h incubation with 20 mM DTT to eliminate residual IAA. The alkylated Env samples were buffer exchanged into 50 mM Tris/HCl, pH 8.0 using Vivaspin columns (3 kDa) and three of the aliquots were digested separately overnight using chymotrypsin (Mass Spectrometry Grade, Promega) or alpha lytic protease (Sigma Aldrich) at a ratio of 1:30 (w/w). The next day, the peptides were dried and extracted using an Oasis HLB µElution Plate (Waters).

The peptides were dried again, re-suspended in 0.1% formic acid, and analyzed by nanoLC-ESI MS with an Easy-nLC 1200 (Thermo Fisher Scientific) system coupled to a Fusion mass spectrometer (Thermo Fisher Scientific) using stepped higher energy collision-induced dissociation (HCD) fragmentation. Peptides were separated using an EasySpray PepMap RSLC C18 column (75 µm × 75 cm). A trapping column (PepMap 100 C18 3μM 75μM × 2cm) was used in line with the LC prior to separation with the analytical column. The LC conditions were as follows: 275-minute linear gradient consisting of 0-32% acetonitrile in 0.1% formic acid over 240 minutes followed by 35 minutes of 80% acetonitrile in 0.1% formic acid. The flow rate was set to 300 nL/min. The spray voltage was set to 2.7 kV and the temperature of the heated capillary was set to 55 °C. The ion transfer tube temperature was set to 275 °C. The scan range was 375−1500 m/z. The stepped HCD collision energies were set to 15, 25 and 45% and the MS2 for each energy was combined. Precursor and fragment detection were performed using an Orbitrap at a resolution MS1 = 100,000. MS2 = 30,000. The AGC target for MS1 =4e5 and MS2 =5e4 and injection time: MS1 =50ms MS2 =54ms.

Glycopeptide fragmentation data were extracted from the raw file using Byos (Version 4.6; Protein Metrics Inc.). The glycopeptide fragmentation data were evaluated manually for each glycopeptide; the peptide was scored as true-positive when the correct b and y fragment ions were observed along with oxonium ions corresponding to the glycan identified. The MS data where searched using the Protein Metrics 38 insect N-glycan library. All charge states for a single glycopeptide were summed. The precursor mass tolerance was set at 4 ppm and 10 ppm for fragments. A 1% false discovery rate (FDR) was applied. The relative amounts of each glycan at each site as well as the unoccupied proportion were determined by comparing the extracted ion chromatographic areas for different glycopeptides with an identical peptide sequence. Glycans were categorized according to the composition detected.

HexNAc(2)Hex(9−3) was classified as M9 to M3. Any of these structures containing a fucose were categorized as FM (fucosylated mannose). Complex-type glycans were classified according to the number of HexNAc subunits and the presence or absence of fucose. Core glycans refer to truncated structures smaller than M3. Paucimannose glycans refer to structures HexNAc(2)Hex(1-3) and include fucosylated structures. As this fragmentation method does not provide linkage information, compositional isomers are grouped.

### Biolayer interferometry

Antigenic profiles of Hi5- and CHO^FucT3^-expressed BG505 Env immunogens, in unmodified and glycan-trimmed forms, were measured on an Octet RED96 (FortéBio, Pall Life Sciences) using a previously described protocol and antibody panel (*65*). All assays were performed at 30 °C with agitation at 1,000 rpm in FortéBio 1× kinetic buffer, with a final volume of 200 μL per well, using solid black 96-well plates (Greiner Bio-One). For all antigens, 5 μg/mL antibody in 1× kinetic buffer was loaded onto anti-human IgG Quantitation (AHQ) biosensors for 300 s. For Alloy mouse-derived human IgGs C10, D9, B10, C5, and B8, anti-human IgG Analytical (AHC) biosensors were used instead. A 2-fold concentration gradient of antigen, starting at 266 nM for trimers and 5.6 nM for SApNPs, was prepared in a six-point dilution series. After a 60-s baseline step, antibody-loaded biosensors were immersed in antigen solutions for a 200-s association step, followed by a 300-s dissociation step. Baseline drift was corrected by subtracting the mean signal shifts from an antibody-loaded sensor not exposed to antigen and an unloaded sensor exposed to antigen. Octet data were processed using FortéBio’s Data Analysis software (version 8.1). Peak response signals at the highest antigen concentration were summarized in a matrix and color-coded for comparison. Experimental data for each antigen-antibody pair were fitted to a 1:1 binding model for visualization and qualitative comparison. Binding curves were plotted using GraphPad Prism (version 10.3.1).

### Enzyme-linked immunosorbent assay (ELISA)

For detection of core α(1,3)-fucose in the Env glycan shield, Costar 96-well high-binding flat-bottom half-area plates (Corning) were coated with 50 µL of PBS containing 0.1 μg of unmodified or glycan-trimmed BG505 UFO trimer produced in the Hi5 or CHO^FucT3^ expression systems. Plates were incubated overnight at 4°C and washed five times with PBST (PBS containing 0.05% (v/v) Tween 20). Wells were blocked for 1 h at room temperature with 150 µL of blocking buffer consisting of 4% (w/v) blotting-grade blocker (Bio-Rad) in PBS, followed by five washes with PBST. Three core α(1,3)-fucose-specific antibody reagents were used: IgG versions of HHH1 (IgG-H1) and HMM5 (IgG-M5) (*132*), and a rabbit polyclonal antibody (Agrisera; cat. no. AS07 268). For binding analysis of single-cell-derived Alloy antibodies, plates were coated with ExpiCHO-produced BG505 UFO trimer under the same conditions. All 18 cloned IgGs were tested, followed by seven selected Fabs, with VRC01 and PGT128 included as reference antibodies. Antibodies were diluted in blocking buffer to a maximum concentration of 1 µg/mL, followed by a 10-fold serial dilution. For each dilution, 50 µL was added to wells and incubated for 1 h at room temperature. Plates were washed five times with PBST. For secondary detection, goat anti-human IgG (1:5,000), goat anti-human IgG, F(ab′)₂ fragment specific (1:3,000), or goat anti-rabbit IgG (1:2,000) antibodies (Jackson ImmunoResearch Laboratories) were diluted in PBST and used as appropriate, with 50 µL added per well. Plates were incubated for 1 h at room temperature and washed six times with PBST. Wells were developed with 50 µL of 3,3’,5,5’-tetramethylbenzidine (Life Sciences) for 3–5 min, and reactions were stopped with 50 µL of 2 N sulfuric acid. Absorbance was measured at 450 nm using a BioTek Synergy plate reader. EC_50_ values were calculated from full binding curves using GraphPad Prism (version 10.3.1). For the core α(1,3)-fucose detection assay, samples with A450 values below 0.5 were assigned an EC_50_ value of 1 µg/mL to facilitate plotting and comparison.

For mouse serum IgG and IgE analysis, plates were coated with 50 µL of PBS containing 0.1 μg of the ExpiCHO-produced BG505 UFO trimer. Mouse sera were diluted 1:40 in blocking buffer and subjected to a 10-fold serial dilution. For each dilution, 50 µL was added per well and incubated for 1 h at room temperature, followed by five washes with PBST. Secondary antibodies included goat anti-mouse IgG (1:3,000; Jackson ImmunoResearch Laboratories) or goat anti-mouse IgE (1:2,000; Thermo Fisher), diluted in PBST. Wells were developed and read as described above. EC_50_ titers were calculated for serum IgG using GraphPad Prism (version 10.3.1). Due to low IgE responses, EC_50_ values could not be calculated, and A450 values were reported for IgE.

### Negative-stain electron microscopy (nsEM)

The experimental and computational protocols used for nsEM analysis were adapted from our previous studies (*65, 66, 152, 153*). BG505 UFO trimers, trimer–Fab complexes, and trimer-presenting SApNPs were analyzed at the Core Microscopy Facility at The Scripps Research Institute. Samples were prepared at concentrations of 0.008–0.015 mg/mL for trimers and trimer– Fab complexes and 0.01 mg/mL for SApNPs. Carbon-coated copper grids (400 mesh) were glow-discharged, and 8 µL of sample was adsorbed for 2 min. Excess sample was removed, and grids were stained with 2% uranyl formate for 2 min. Excess stain was wicked away, and grids were air-dried. BG505 UFO trimers and trimer-Fab complexes were analyzed at 120 kV using a Talos L120C transmission electron microscope (Thermo Fisher Scientific). Images were acquired with a CETA 16M CMOS camera at 73,000× magnification, corresponding to a pixel size of 1.93 Å and a defocus range of 0.5–2 µm. SApNP images were collected at 52,000× magnification, corresponding to a pixel size of 2.05 Å at the specimen plane with a defocus of −1.50 µm.

Computational analysis of trimer and trimer–antibody datasets was performed using the high-performance computing core facility at The Scripps Research Institute (TSRI). TIFF-format nsEM images were converted to MRC format using EMAN2 (*214*) and processed in CryoSPARC (version 5.0) (*215*). Contrast transfer function (CTF) parameters were estimated using Patch CTF Estimation. Particles were selected using the Blob Picker or Template Picker, with key parameters optimized for each dataset based on particle size and complex stoichiometry. Selected particles were subjected to iterative 2D classification to remove contaminants and poorly defined classes. For trimer–Fab datasets derived from Alloy antibodies, multiple rounds of particle picking and 2D classification were performed as needed to account for variable Fab occupancy, conformational heterogeneity, and particle orientation. Well-defined classes were used for *ab initio* reconstruction, followed by heterogeneous and/or homogeneous refinement. Final nsEM density maps and fitted structures were visualized and rendered using UCSF Chimera (*216*).

To facilitate comparison of experimental 2D class averages and 3D nsEM maps with known Env epitopes, simulated 30-Å-resolution density maps were generated for 16 representative bNAbs and one non-NAb in complex with the BG505 UFO trimer. Available Env–antibody complex structures were used as templates to position the corresponding Fabs on the BG505 UFO trimer structure (*154*) and to build models of BG505 UFO–Fab complexes. Simulated density maps were calculated using pdb2mrc in EMAN2 (*214*).

### Crystallization and data collection

Complexes of Hi5-produced BG505 UFO trimers with two antibody fragments were prepared for crystallization at a molar ratio of 1:3.5:3.5 (trimer:Fab/scFv:Fab). For the unmodified trimer– antibody complex, purified trimer was mixed with 3H109L Fab and 35O22 scFv. For the glycan-trimmed trimer–antibody complex, purified trimer was first treated with endoglycosidase H (Endo H) for 30 min at room temperature and then mixed with PGT124 Fab and 35O22 Fab. Complexes were purified by size-exclusion chromatography (SEC) on a Superdex 200 column to isolate the monodisperse trimer–antibody complex peak. Purified complexes were concentrated to ∼10 mg/mL and subjected to crystallization screening at 20°C using the sitting-drop vapor diffusion method on a Rigaku CrystalMation robotic system. Diffraction-quality crystals were obtained in 0.1 M Tris (pH 8.0), 15% (w/v) polyethylene glycol monomethyl ether 2000, and 0.1 M potassium chloride for the unmodified complex; and in 0.1 M HEPES (pH 7.5) with 10% (w/v) polyethylene glycol 8000 for the Endo H–treated complex. Crystals were cryoprotected with 15% glycerol or 15% ethylene glycol, respectively, and flash-cooled in liquid nitrogen. Diffraction data were collected at beamline 12-1 of the Stanford Synchrotron Radiation Lightsource.

### Structure determination and refinement

Crystals of the unmodified trimer in complex with 3H109L and 35O22 diffracted to a maximum resolution of 4.46 Å, and crystals of the Endo H–treated trimer in complex with PGT124 and 35O22 diffracted to 4.09 Å. Diffraction data were indexed, integrated, and scaled using HKL-2000 (*217*) in space group R32. Phases were obtained by molecular replacement in Phaser (*218*) using BG505 UFO (PDB: 6UTK), PGT124 (PDB: 6CE0), 35O22 (PDB: 6CE0), and 3H109L (PDB: 6MTJ) as search models. Iterative model rebuilding was carried out in Coot (*219*), and refinement was performed in Phenix (*220*). N-linked glycans were modeled conservatively only where supported by interpretable electron density, with glycan length at each site limited to the ordered portion visible in the maps. Data collection and refinement statistics are provided in **Table S1**.

### Mouse immunization and sample collection

Mouse immunization protocols were adapted from our previously described methods (*65, 66, 152, 153*). All animal experiments were conducted in accordance with the guidelines of the Association for the Assessment and Accreditation of Laboratory Animal Care (AAALAC) and approved by the Institutional Animal Care and Use Committee (IACUC) of The Scripps Research Institute. Female and male BALB/c mice (6–8 weeks old) were purchased from The Jackson Laboratory and housed in ventilated cages under environmentally controlled conditions (20°C, 50% humidity, 12 h light–dark cycle) at an animal biosafety level 2 (ABSL2) facility. Mice were immunized via the intraperitoneal (i.p.) route at weeks 0, 3, 6, and 9 at 3-week intervals. Each dose consisted of 200 μl of antigen/adjuvant mixture containing 10 μg of antigen and 100 μl of adjuvant. The adjuvant was 100 μl aluminum hydroxide (AH; InvivoGen; cat. no. vac-alu-50) or 40 μg (50 μl) CpG (ODN 1826; InvivoGen; cat. no. tlrl-1826) adsorbed onto 50 μl AH. Blood samples were collected two weeks after each injection via the retro-orbital sinus using heparinized capillary tubes. Samples were centrifuged at 13,200 × g for 10 min to separate serum. Following heat inactivation at 56°C for 30 min, sera were centrifuged again at 13,200 × g for 10 min to remove precipitates. At the final time point (week 11), spleens were harvested and dissociated through a 70-μm cell strainer (BD Falcon) to obtain single-cell suspensions. Splenocytes were centrifuged, washed in PBS, treated with 5 ml ACK lysing buffer (Lonza), and frozen in 3 ml freezing medium (10% DMSO/90% FCS). Serum samples from the Hi5 study and selected groups from the CHO^FucT3^ study were purified using CaptureSelect IgG-Fc (Multispecies) Affinity Matrix (Thermo Scientific) according to the manufacturer’s instructions. Purified IgG from individual mice was analyzed in TZM-bl assays to assess vaccine-induced NAb responses.

Humanized mouse immunization protocols were adapted from wild-type mouse protocols, with modifications to the antigen dose and adjuvant. Six female humanized transgenic mice—two ATX-GK and four ATX-GKH (Hyperimmune) mice—were obtained from Alloy Therapeutics under an academic license. Humanized mice were immunized intraperitoneally (i.p.) at weeks 0, 3, 6, and 9. Each dose consisted of 200 μl of antigen/adjuvant mixture containing 50 μg of antigen and 100 μl of adjuvant. The adjuvant consisted of 20 μg (1 μl) STING agonist (2′3′-cGAMP; Millipore Sigma; cat. no. 1441190-66-4) adsorbed onto 100 μl AH (InvivoGen). Blood samples were collected two weeks after each immunization, and sera or purified IgG were analyzed in TZM-bl neutralization assays.

### Pseudovirus production and neutralization assays

Pseudoviruses were generated by co-transfection of HEK293T cells (ATCC; cat. no. CRL-3216) with an HIV-1 Env–expressing plasmid and an Env-deficient genomic backbone plasmid (pSG3ΔEnv), as previously described (*65*). HIV-1 Env–expressing vectors for BG505 (cat. no. 11518), SF162 (cat. no. 10463), the 12-virus global panel (*170*) (cat. no. 12670), as well as TZM-bl cells, were obtained through the NIH AIDS Reagent Program managed by BEI Resources (https://www.beiresources.org/HIV.aspx). A T332N mutation was introduced into BG505 Env to generate the BG505.T332N clone. Pseudoviruses were harvested 72 h post-transfection and stored at −80°C until use. Neutralizing activity of each mouse sample was assessed using a single-round replication pseudovirus assay with TZM-bl target cells (*221*). Briefly, pseudovirus was incubated with serial dilutions of purified IgG or heat-inactivated sera in 96-well flat-bottom plates for 1 h at 37°C prior to addition of TZM-bl cells. For purified mouse IgG, a starting concentration of 300 μg/ml was used; for mouse sera, an initial 1:100 dilution was used, both followed by 3-fold serial dilutions. Serum samples were tested in duplicate across the full dilution series, whereas purified IgG samples were tested in singlet due to limited sample volume. For Alloy-derived human antibodies (IgG and Fab), starting concentrations were selected according to antibody potency and assay requirements: 1,000 μg/ml for weak NAbs, 300 μg/ml for C10, 100 μg/ml for SF162 in variant analysis, and 10 μg/ml for VRC01. For IC_50_ summary and color-coding in **Fig. 7f**, **Fig. 7g**, **Fig. 8e**, and **Fig. S8i**, 300 μg/ml was used as the reporting cutoff, irrespective of the higher starting concentration (1,000 μg/ml) used for some weakly neutralizing mAbs. Luciferase reporter activity was measured 60–72 h post-infection following cell lysis and addition of Bright-Glo Luciferase substrate (Promega). Luminescence was quantified using a BioTek microplate reader with Gen5 software. Percent neutralization was calculated relative to virus-only control wells, with background luminescence from uninfected cell controls subtracted. Dose–response curves were fitted by nonlinear regression in GraphPad Prism (version 9.3.1) using a 0–100% constraint. IC_50_ values were derived for purified IgG, single-cell-sorted Alloy antibodies (IgG and Fab), and reference bNAbs, while ID_50_ values were calculated for serum. Pseudoparticles displaying the envelope glycoproteins of MLV (MLV-pps) were included as a negative control and tested under identical conditions in the neutralization assays.

### Env-specific sorting of Alloy humanized mouse B cells, antibody cloning, and production

Alloy mouse splenocytes collected 14 days after the final immunization were prepared for antigen-specific B-cell sorting using a protocol adapted from the ATX-Gx™ User Guide (2024 Edition), with modifications where appropriate. Briefly, cells were first stained to exclude dead cells using Fixable Aqua Dead Cell Stain (Thermo Fisher). FcγIII (CD16) and FcγII (CD32) receptors were blocked with 20 μl of 2.4G2 mAb (BD Pharmingen). Cells were then incubated with 10 μg of biotinylated Avi-tagged BG505 UFO trimer probe for 5 min at 4 °C, followed by 2.5 μl of FITC-labeled anti-mouse IgG (Jackson ImmunoResearch) for 15 min at 4 °C. Finally, 5 μl of premium-grade APC-labeled streptavidin (Thermo Fisher) was added for 15 min at 4 °C. Between steps, cells were washed with 500 μl of FACS buffer (DPBS containing 2% FBS). FITC^+^APC^+^ Env-specific B cells were sorted using a MoFlo Astrios EQ (Beckman Coulter).

For antibody cloning, 3 μL of Random Hexamers (GeneLink), 2 μL of dNTPs, and 1 μL of SuperScript IV enzyme (Thermo Fisher) were preloaded into each well of a 96-well plate before single-cell sorting. Reverse transcription (RT) was performed according to the SuperScript IV protocol, yielding 25 μl of cDNA per cell. cDNA (5 μl) was then added to a PCR mix containing 12.5 μl of GoTaq Master Mix (Promega M7133), 5.5 μl of H2O, 1 μl of forward primer mix, and 1 μl of reverse primer mix for amplification of HCs and KCs (**Fig. S7a**; ATX-Gx^TM^ User Guide, pp. 22–23). A second PCR was performed using 5 μl of the first PCR product as template, with the corresponding nested primers and the same reaction mixture. PCR products were analyzed by 1% agarose gel electrophoresis, and products of the expected HC and KC sizes were validated by Sanger sequencing (GeneWiz). Sequences were analyzed using IMGT (http://www.imgt.org/) for germline gene and lineage assignment. Selected paired HC and KC sequences were amplified to introduce 5′ and 3′ restriction sites (AgeI and SalI for HC; AgeI and BsiWI for KC), and ligated into the corresponding heavy- and light-chain mammalian expression vectors.

For antibody production, paired HC and KC sequences were cloned into IgG1 expression vectors and transiently co-transfected at a 1:1 ratio into ExpiCHO cells (Thermo Fisher) using the ExpiFectamine CHO Transfection Kit according to the manufacturer’s instructions. Supernatants were harvested 12–14 days after transfection, clarified by centrifugation, and filtered through a 0.45-μm membrane. IgGs were purified by Protein A affinity chromatography (Thermo Fisher), eluted with 0.3 M citric acid (pH 3.0), and neutralized with 2 M Tris base (pH 9.0), whereas Fabs were purified by Protein G affinity chromatography (Thermo Fisher),, eluted with 0.2 M glycine (pH 2.2), and neutralized with 2 M Tris base (pH 9.0). Purified antibodies were concentrated and buffer-exchanged into PBS using 10-kDa centrifugal filters, with concentrations determined by UV_280_ absorbance using theoretical extinction coefficients.

### NGS and bioinformatics analysis of Alloy mouse B cells

Total RNA (including mRNA) was extracted from 10–20 million splenocytes from G2M1 and G2M2 using the RNeasy Mini Kit (Qiagen) and eluted in 30 μL of water. Reverse transcription (RT) was performed with 2 μg of total RNA according to the SuperScript IV protocol, yielding 25 μL of cDNA. Heavy- and light-chain variable regions were amplified from 5 μL of cDNA by two rounds of PCR using the same primer sets and reaction conditions described above for single-cell antibody cloning (**Fig. S7a**; ATX-Gx™ User Guide, pp. 22–23). PCR products of the expected size (∼400 bp) were resolved by 1% agarose gel electrophoresis and gel-purified (Qiagen). For NGS library preparation, 40 ng of each heavy- or light-chain amplicon was processed using the NEBNext Ultra II DNA Library Prep Kit for Illumina (New England Biolabs; E7645L), including end repair and adaptor ligation. After bead cleanup, NEB dual indexes were added to individual libraries. Final libraries were loaded at 9 pM onto an Element Biosciences AVITI platform for 600-cycle single-end sequencing, yielding ∼120 million passing-filter reads in total.

Sequencing data were processed using a modified version of the human Antibodyomics pipeline, with additional filtering steps to remove the poly-C tail from raw reads in Step 1 and exclude any sequences containing translation errors in Step 5 (**Fig. S8d**). Quantitative repertoire profiles were generated for germline gene usage, somatic hypermutation (SHM), and HCDR3 and KCDR3 loop length. Two-dimensional (2D) divergence/identity plots were generated to visualize selected antibody lineages within the G2M1 and G2M2 B-cell repertoires. Putative somatic variants of C10, B10, C5, and B8 were identified based on germline gene assignment and ≥95% CDR3 nucleotide sequence identity. NGS-derived C10 and B10 HC variants and putative germline precursors were selected for phylogenetic analysis and experimental validation. Selected HCs were paired with their respective parental light chains and transiently expressed in ExpiCHO cells (Thermo Fisher), followed by purification from culture supernatants after 12–14 days by Protein A affinity chromatography (Thermo Fisher).

### Statistical analysis

Data were collected from *n* = 8 mice per group in the wild-type mouse immunization studies and from *n = 2* mice per group in the ATX-Gx humanized mouse immunization study. All ELISA and serum neutralization assays were performed in duplicate, whereas IgG neutralization assays were performed in singlet to preserve samples. For CHO^FucT3^-expressed immunogens, differences among immunogen formats (trimer *vs.* SApNP) and glycan states (unmodified *vs.* glycan-trimmed) were compared using one-way analysis of variance (ANOVA), followed by Tukey’s multiple-comparisons post hoc test. Two-tailed unpaired *t-*tests were used to compare two independent vaccine groups. Statistical significance is indicated in the figures as follows: ns (not significant), \**p* < 0.05, \*\**p* < 0.01, \*\*\**p* < 0.001, and \*\*\*\**p* < 0.0001. Graphs were generated and statistical analyses were performed using GraphPad Prism (version 10.3.1).

## ACKNOWLEDGEMENTS

We acknowledge K. Vanderpool and S. Henderson of the Core Microscopy Facility at TSRI for their expert assistance with negative-stain EM work. We also thank A. Ward and G. Ozorowski for helpful discussions regarding negative-stain EM analysis and data interpretation. We also acknowledge S. Wang and J Shimashita of the Genomics Core Facility at TSRI for their expert assistance with Alloy mouse B-cell repertoire sequencing.

## Author contributions

Project design by L.H., X.H.L, I.A.W, and J.Z.; Hi5 and CHO^FucT3^ expression and purification of BG505 Env immunogens by C.D. and L.H.; SDS-PAGE and BN-PAGE by C.D. and L.H.; DSC and DLS by L.H.; ELISA by Y.-N.Z. and L.H.; BLI by L.H.; glycan analysis by M.N., J.D.A., and M.C.; negative-stain EM by L.H. and J.Z.; EM data interpretation by J.Z. and A.B.W.; X-ray crystallography and structural analysis by X.H.L. and I.A.W.; TZM-bl neutralization assays and data analysis by C.D., Y.-N.Z., L.H., and J.Z.; Env-specific single-cell sorting, antibody cloning, and antibody production by L.H.; NGS library preparation by L.H.; bioinformatics analysis by J.Z.; manuscript writing by L.H., Y.-N.Z., M.N., and J.Z. All authors commented on the manuscript.

## Data and material availability

All data needed to evaluate the conclusions of this research are available in the main text and Supplementary Information. Structural data obtained from X-ray crystallography have been deposited in the Protein Data Bank (PDB, https://www.rcsb.org/) under accession codes 38NW and 38NX. Mass spectrometry data have been deposited in the MassIVE server (https://massive.ucsd.edu) under accession code MSV000102992. All data supporting the findings of this study are available within the article and its Supplementary Information. Source data are provided with this paper.

