## Supplementary Information for "Xenoglycosylation of the HIV-1 Glycan Shield Promotes Cross-Clade Neutralization and Diverse Human Antibody Lineages"

**Table S1. X-ray data collection and refinement statistics.**

| **Data collection** | Hi5-produced BG505 UFO trimer + 3H109L Fab + 35022 scFv | Hi5-produced BG505 UFO trimer (Endo H-treated) + PGT124 Fab + 35022 Fab |
|---|---|---|
| Beamline | SSRL 12-1 | SSRL 12-1 |
| Wavelength (Å) | 0.9795 | 0.9795 |
| Space group | R32 | $P6_3$ |
| Unit cell parameters | | |
| a, b, c (Å) | 202.7 202.7 509.4 | 125.6 125.6 315.5 |
| α, β, γ (°) | 90, 90, 120 | 90 90 120 |
| Resolution (Å) [a] | 50.00-4.46 (4.54-4.46) | 50.00-4.09 (4.16-4.09) |
| Unique reflections [a] | 24,927 (1,225) | 21,801 (1,008) |
| Redundancy [a] | 8.6 (8.3) | 6.0 (4.3) |
| Completeness (%) [a] | 100.0 (100.0) | 98.2 (90.7) |
| $<I/\sigma_I>$ [a] | 4.4 (1.0) | 4.9 (1.0) |
| $R_{sym}$ (%) [a,b] | 48.4 (>100) | 31.4 (>100) |
| $R_{pim}$ (%) [a,b] | 17.5 (>100) | 12.8 (>100) |
| $CC_{1/2}$ (%) [a] | 88.5 (37.9) | 98.8 (41.3) |
| **Refinement statistics** | | |
| Resolution (Å) | 49.42-4.46 | 49.13-4.09 |
| Reflections (work) | 22,798 | 20,286 |
| Reflections (test) | 1,240 | 990 |
| $R_{cryst}$ / $R_{free}$ (%) [c] | 29.4/31.9 | 30.6/35.3 |
| No. of atoms | 10,714 | 11,791 |
| Antigen | 4,541 | 4,516 |
| Fab/scFv | 5,154 | 6,749 |
| Glycan | 1,019 | 526 |
| Average *B*-values (Å$^2$) | 245 | 222 |
| Antigen | 225 | 203 |
| Fab/scFv | 252 | 232 |
| Glycan | 299 | 248 |
| Wilson *B*-value (Å$^2$) | 183 | 172 |
| No. of protomers per ASU [d] | 1 | 1 |
| **RMSD from ideal geometry** | | |
| Bond length (Å) | 0.006 | 0.003 |
| Bond angle (°) | 1.38 | 0.70 |
| **Ramachandran statistics (%)** | | |
| Favored | 91.1 | 93.6 |
| Outliers | 0.97 | 0.90 |
| PDB ID | 38NW | 38NX |

[a] Parentheses denote statistics for the outer resolution shell.

[b] $R_{sym} = \sum_{hkl}\sum_i |I_{hkl,i} - <I_{hkl}>| / \sum_{hkl}\sum_i I_{hkl,i}$, and $R_{pim} = \sum_{hkl}[1/(N-1)]^{1/2}\sum_i |I_{hkl,i} - <I_{hkl}>| / \sum_{hkl}\sum_i I_{hkl,i}$, where $I_{hkl,i}$ is the scaled intensity of the i$^{th}$ measurement of reflection *h*, *k*, *l*, $< I_{hkl}>$ is the average intensity for that reflection, and *N* is the redundancy.

[c] $R_{cryst} = \sum_{hkl} |F_o - F_c| / \sum_{hkl} |F_o|$, where $F_o$ and $F_c$ are the observed and calculated structure-factor amplitudes, respectively. $R_{free}$ was calculated in the same manner using the test set excluded from refinement.

[d] No. of protomers refers to the number of HIV-1 Env protomers per asymmetric unit (ASU); each HIV-1 Env trimer contains three protomers.

### Fig. S1

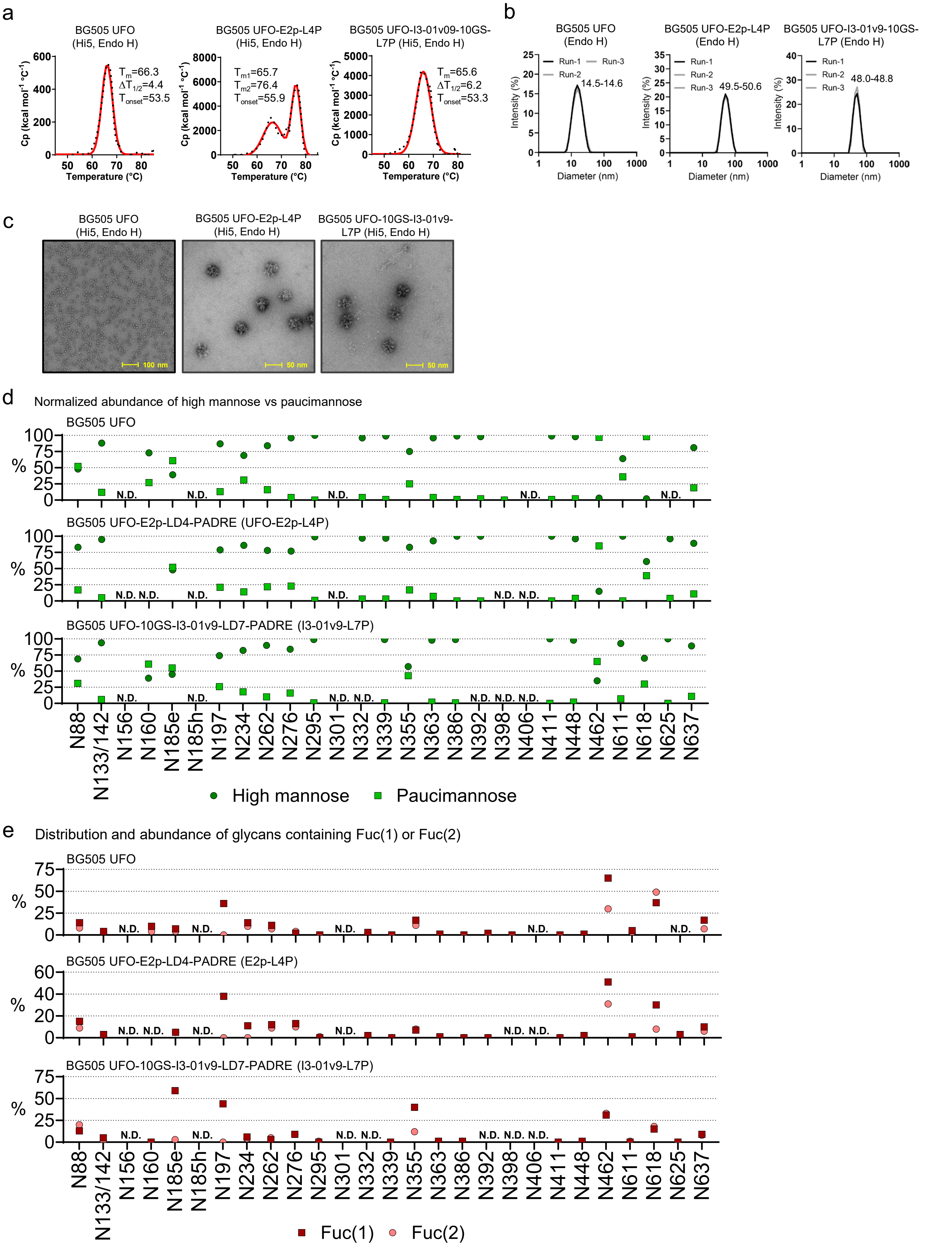
a
BG505 UFO (Hi5, Endo H)
Tm=66.3
ΔT1/2=4.4
Tonset=53.5
BG505 UFO-E2p-L4P (Hi5, Endo H)
Tm1=65.7
Tm2=76.4
Tonset=55.9
BG505 UFO-I3-01v09-10GS-L7P (Hi5, Endo H)
Tm=65.6
ΔT1/2=6.2
Tonset=53.3
Cp (kcal mol-1 °C-1)
Temperature (°C)
b
BG505 UFO (Endo H)
14.5-14.6
BG505 UFO-E2p-L4P (Endo H)
49.5-50.6
BG505 UFO-I3-01v9-10GS-L7P (Endo H)
48.0-48.8
Run-1
Run-2
Run-3
Intensity (%)
Diameter (nm)
c
BG505 UFO (Hi5, Endo H)
BG505 UFO-E2p-L4P (Hi5, Endo H)
BG505 UFO-10GS-I3-01v9-L7P (Hi5, Endo H)
100 nm
50 nm
50 nm
d
Normalized abundance of high mannose vs paucimannose
BG505 UFO
BG505 UFO-E2p-LD4-PADRE (UFO-E2p-L4P)
BG505 UFO-10GS-I3-01v9-LD7-PADRE (I3-01v9-L7P)
N.D.
N88
N133/142
N156
N160
N185e
N185h
N197
N234
N262
N276
N295
N301
N332
N339
N355
N363
N386
N392
N398
N406
N411
N448
N462
N611
N618
N625
N637
High mannose
Paucimannose
e
Distribution and abundance of glycans containing Fuc(1) or Fuc(2)
BG505 UFO
BG505 UFO-E2p-LD4-PADRE (E2p-L4P)
BG505 UFO-10GS-I3-01v9-LD7-PADRE (I3-01v9-L7P)
N.D.
Fuc(1)
Fuc(2)

Fig. S1

f

BG505 UFO (Hi5, unmodified, GNL/SEC)

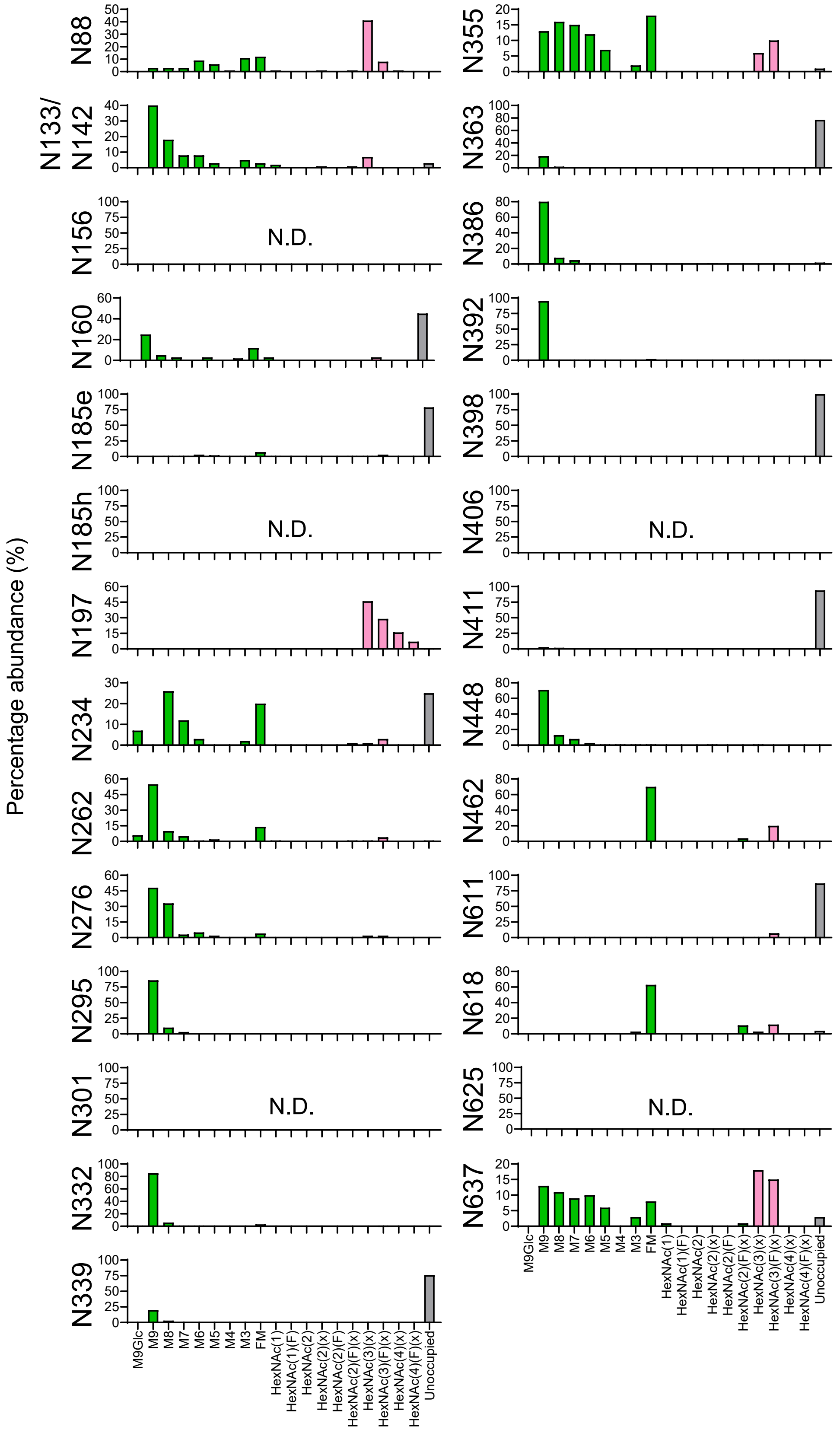

g

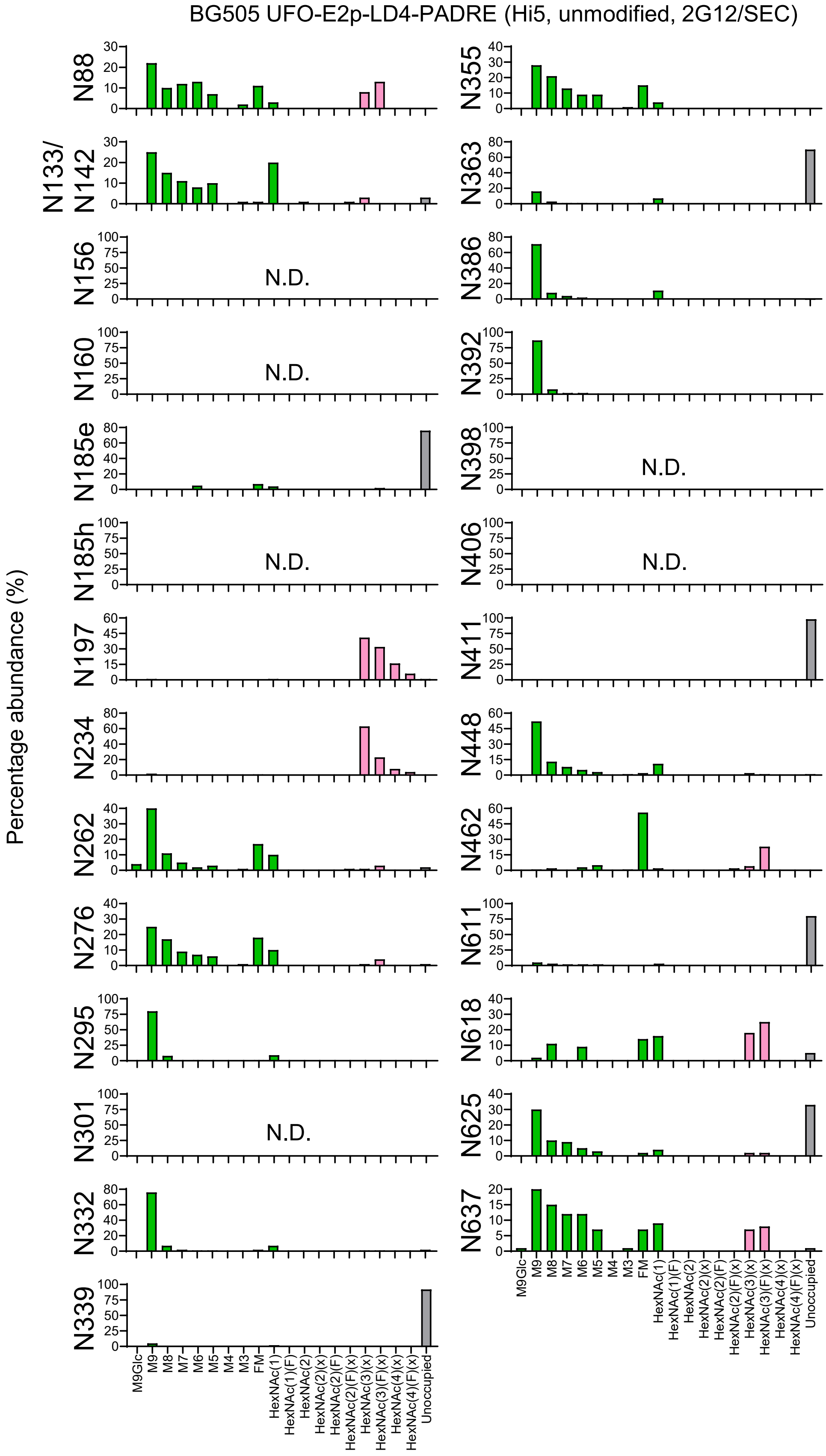
BG505 UFO-E2p-LD4-PADRE (Hi5, unmodified, 2G12/SEC)
Percentage abundance (%)
N88
N133/
N142
N156
N160
N185e
N185h
N197
N234
N262
N276
N295
N301
N332
N339
N355
N363
N386
N392
N398
N406
N411
N448
N462
N611
N618
N625
N637
N.D.
M9Glc
M9
M8
M7
M6
M5
M4
M3
FM
HexNAc(1)
HexNAc(1)(F)
HexNAc(2)
HexNAc(2)(x)
HexNAc(2)(F)
HexNAc(2)(F)(x)
HexNAc(3)(x)
HexNAc(3)(F)(x)
HexNAc(4)(x)
HexNAc(4)(F)(x)
Unoccupied

Fig. S1

h

BG505 UFO-I3-01v9-LD7-PADRE (Hi5, unmodified, 2G12/SEC)

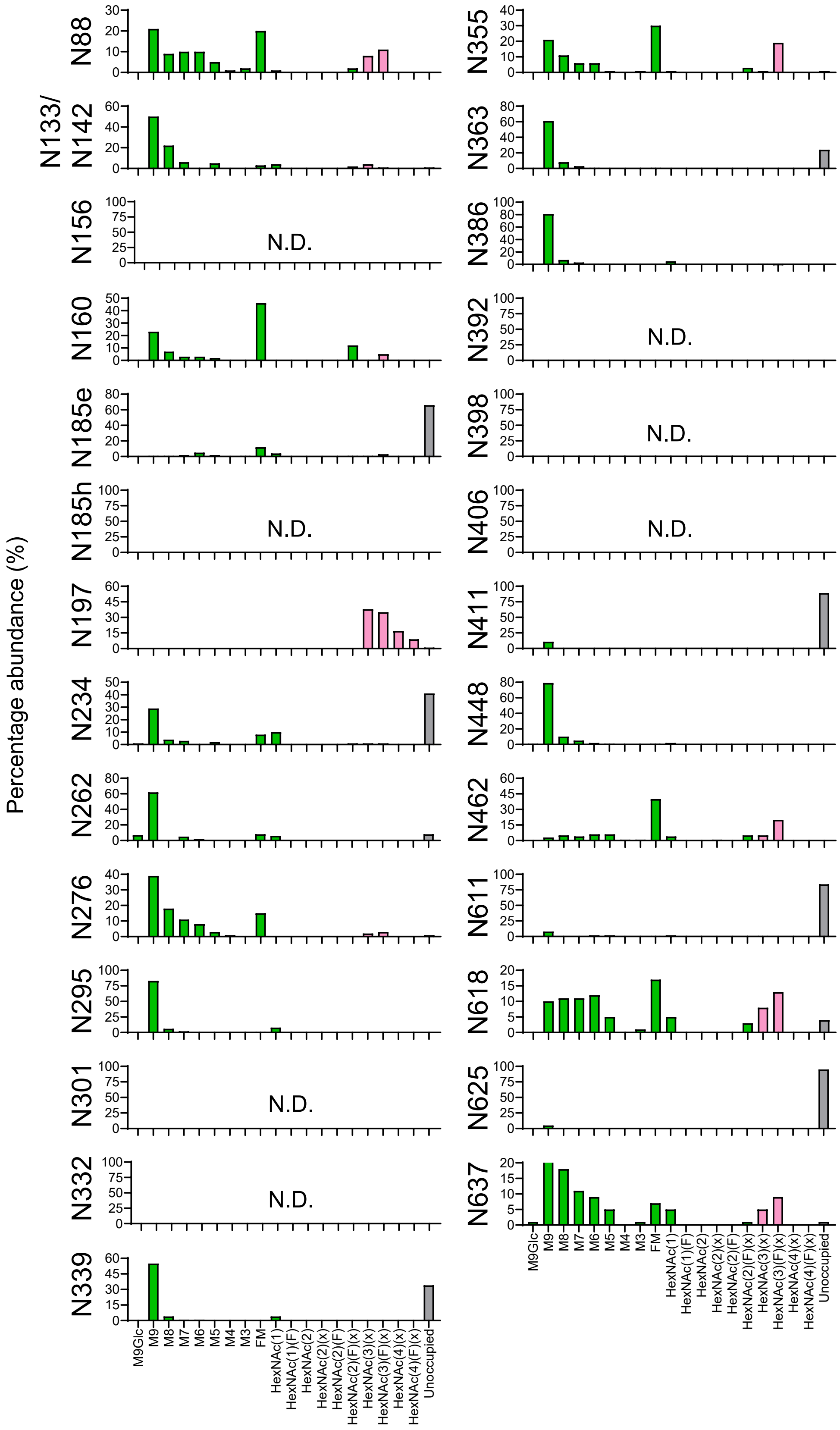

### Fig. S1

i Detection of core α(1,3)-fucose on BG505 UFO trimers produced in ExpiCHO and Hi5 cells

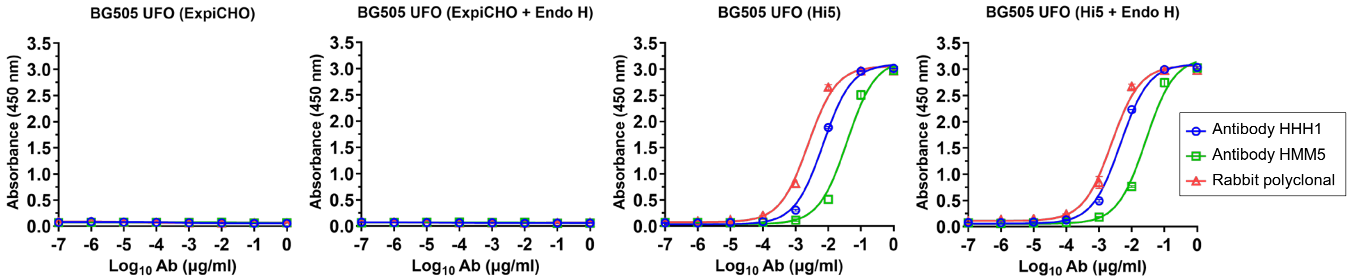


**BG505 UFO (ExpiCHO, GNL/SEC-purified)**

| Antibody HHH1 | | Antibody HMM5 | | Rabbit polyclonal | |
|---|---|---|---|---|---|
| $EC_{50}$ | SD | $EC_{50}$ | SD | $EC_{50}$ | SD |
| 100 | 0 | 100 | 0 | 100 | 0 |

**BG505 UFO (ExpiCHO, GNL/SEC-purified, Endo H-treated)**

| Antibody HHH1 | | Antibody HMM5 | | Rabbit polyclonal | |
|---|---|---|---|---|---|
| $EC_{50}$ | SD | $EC_{50}$ | SD | $EC_{50}$ | SD |
| 100 | 0 | 100 | 0 | 100 | 0 |

**BG505 UFO (Hi5, GNL/SEC-purified)**

| Antibody HHH1 | | Antibody HMM5 | | Rabbit polyclonal | |
|---|---|---|---|---|---|
| $EC_{50}$ | SD | $EC_{50}$ | SD | $EC_{50}$ | SD |
| 0.00694 | 0.000143 | 0.03734 | 0.002355 | 0.00243 | 8.556E-05 |

**BG505 UFO (Hi5, GNL/SEC-purified, Endo H-treated)**

| Antibody HHH1 | | Antibody HMM5 | | Rabbit polyclonal | |
|---|---|---|---|---|---|
| $EC_{50}$ | SD | $EC_{50}$ | SD | $EC_{50}$ | SD |
| 0.00453 | 8.132E-05 | 0.02677 | 0.001039 | 0.00242 | 0.000320 |

**Fig. S1. In vitro characterization of BG505 Env immunogens transiently produced in Hi5 insect cells.** (**a**) DSC thermograms of Hi5-produced, glycan-trimmed BG505 UFO trimers and SApNPs. Experimental data (black dots) and Gaussian fits (red lines) are shown, with key thermal parameters indicated: melting temperature(s) ($T_m$ or $T_{m1}/T_{m2}$), onset temperature ($T_{onset}$), and peak half-width ($\Delta T_{1/2}$). (**b**) Particle size distributions of Hi5-produced, glycan-trimmed BG505 UFO trimers and SApNPs. DLS measurements from three independent production batches are shown for each construct, with hydrodynamic diameters (Dh) indicated. (**c**) Representative nsEM images of produced, glycan-trimmed BG505 UFO trimers (left), E2p-L4P SApNPs (middle), and I3-01v9-L7P SApNPs (right). Scale bar, 100/50 nm. For panels (a–c), prior to Endo H treatment, BG505 UFO trimers were purified from Hi5 supernatant by GNL followed by SEC, whereas SApNPs were purified by IAC using a 2G12 antibody column followed by SEC. (**d**) Normalized abundance of high-mannose versus paucimannose glycans on Hi5-produced BG505 UFO trimers and SApNPs. Glycan types spanning from M9Glc(x)–M4 are classified as high mannose, and glycan types spanning GlcNAc(2)Hex(2) ± Fuc–M3 ± Fuc are classified as paucimannose. (**e**) Abundance of fucosylated glycans containing one [Fuc(1)] or two [Fuc(2)] fucose residues on Hi5-produced BG505 UFO trimers and SApNPs. (**f**–**h**) Site-specific glycan composition analysis of (**f**) BG505 UFO trimers, (**g**) E2p SApNPs, and (**h**) I3-01v9 SApNPs. Quantitative mass spectrometric analysis of glycans at individual N-linked sites is summarized into simplified glycan categories. Oligomannose-type glycans (M9–M5; Man9GlcNAc2 to Man5GlcNAc2) are shown in green; afucosylated and fucosylated hybrid-type glycans (hybrid and F hybrid) are shown as dashed pink; complex glycans are grouped by antenna number and core fucosylation and are shown in pink. Unoccupied glycosylation sites are shown in gray. (**i**) Detection of core α(1,3)-fucosylation on ExpiCHO- and Hi5-produced BG505 UFO trimers by ELISA using core α(1,3)-fucose-specific HHH1 (IgG-H1), HMM5 (IgG-M5), and rabbit polyclonal antibodies. Binding curves (top) and corresponding $EC_{50}$ values with standard deviations (bottom) are shown. Plates were coated with 0.1 µg of antigen, and antibodies were tested in a 10-fold dilution series starting at 1 µg ml$^{-1}$. Error bars represent the difference between duplicate values at each concentration tested for each sample.

Fig. S2

a

Hi5-produced BG505 UFO trimer (AHQ)

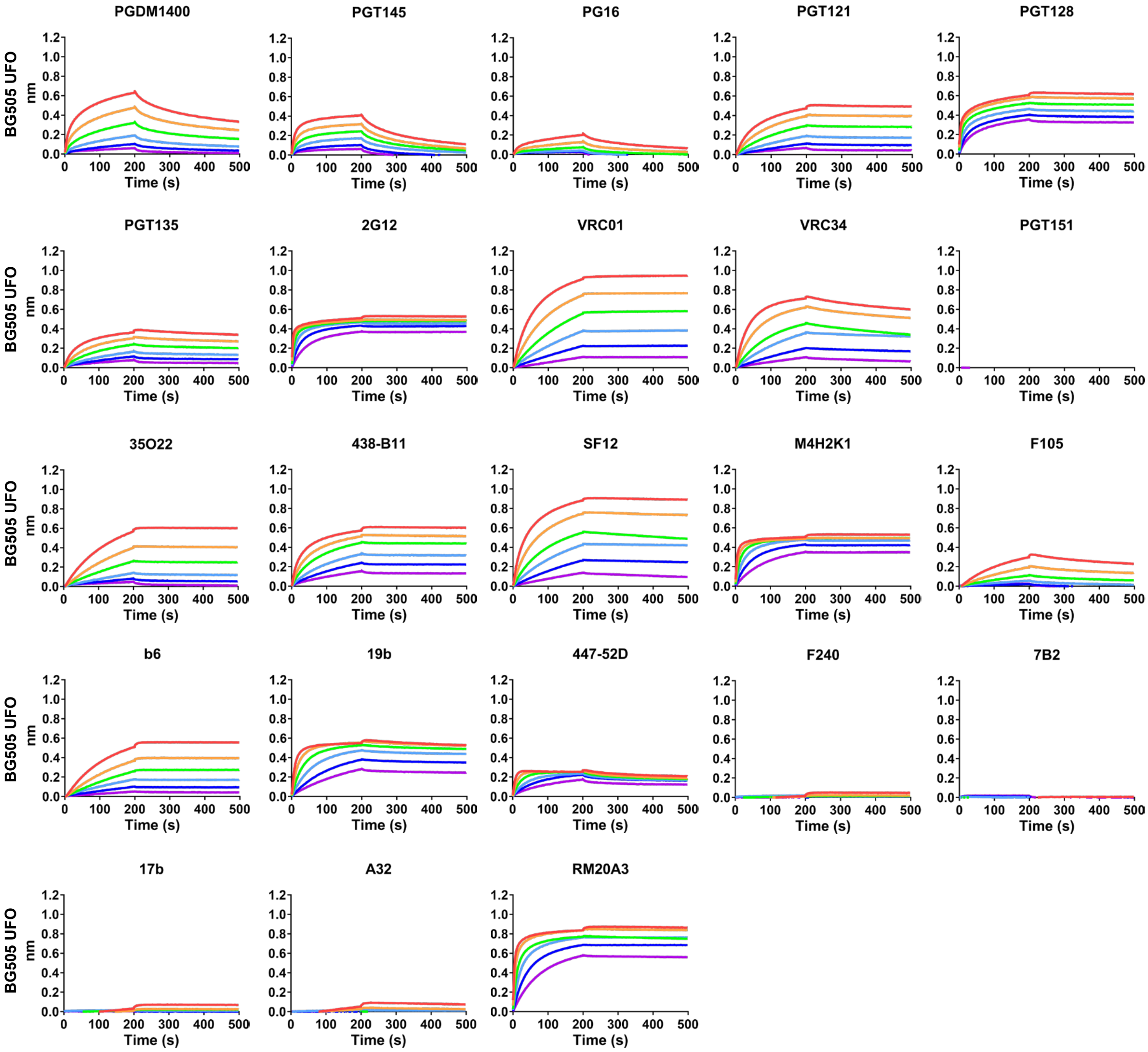

Fig. S2

b

Hi5-produced BG505 UFO-E2p-L4P SApNP (AHQ)

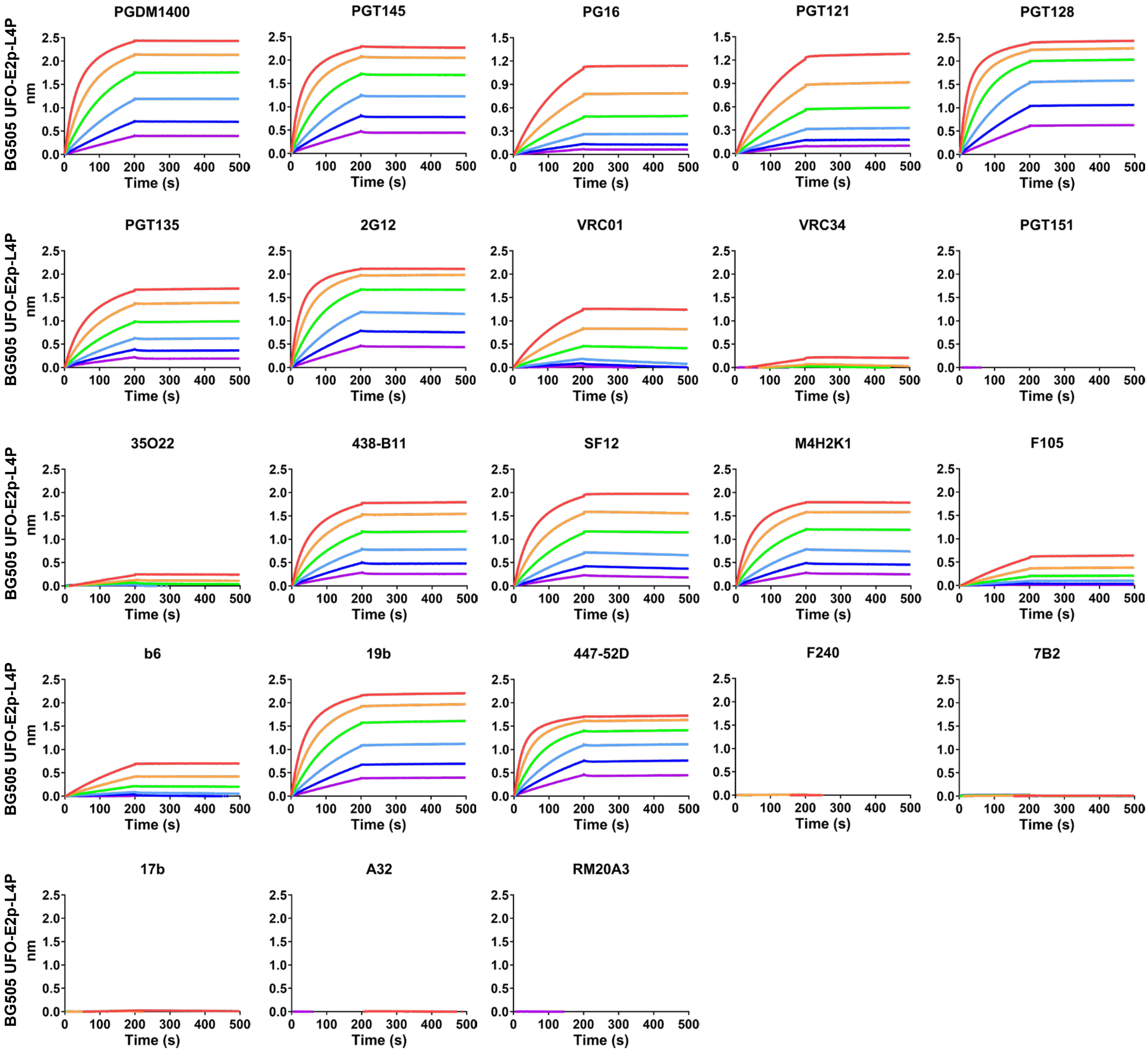

C

Hi5-produced BG505 UFO-10GS-I3-01v9-L7P SApNP (AHQ)

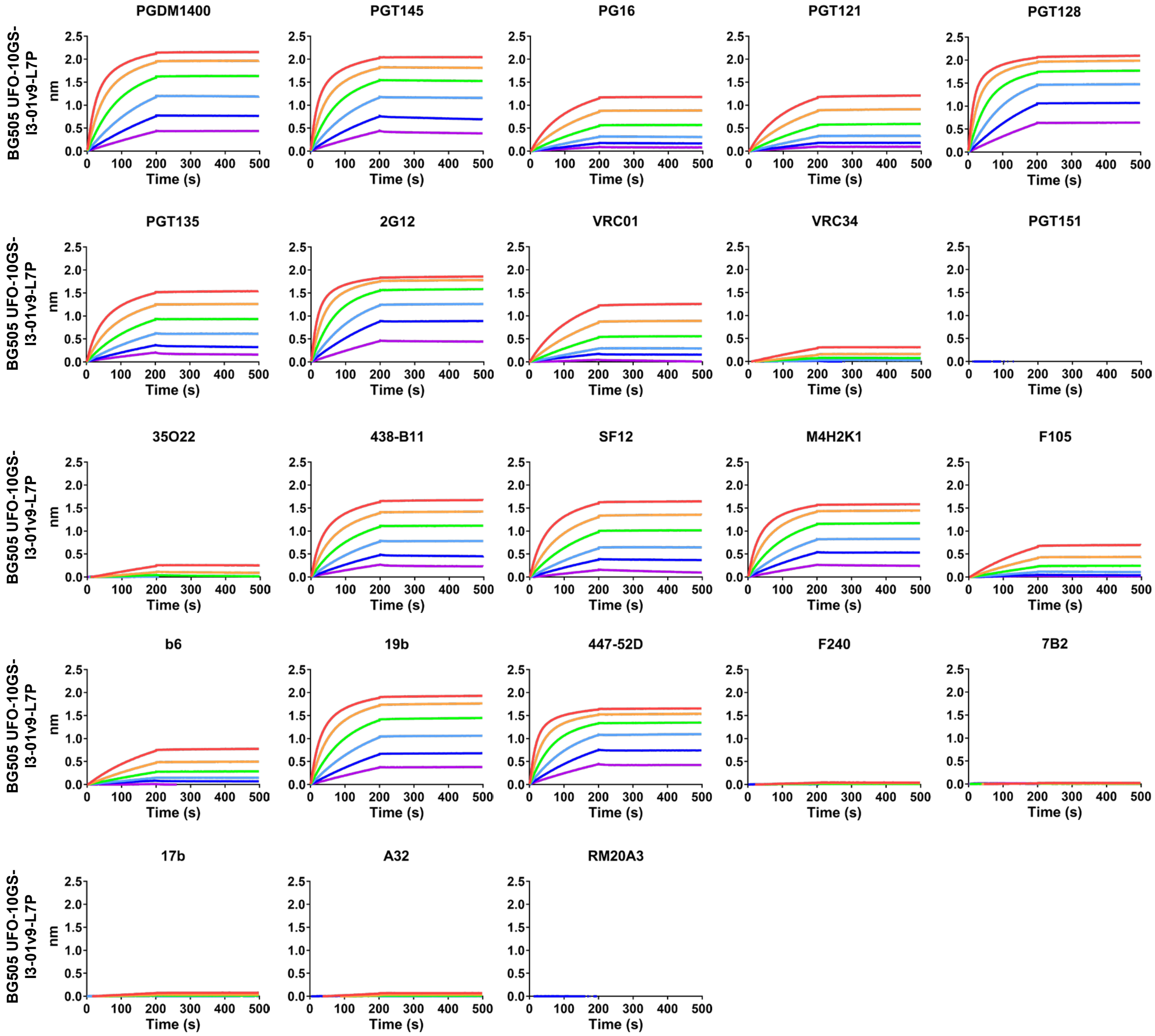

**Fig. S2**

d

Hi5-produced, glycan-trimmed BG505 UFO trimer (AHQ)

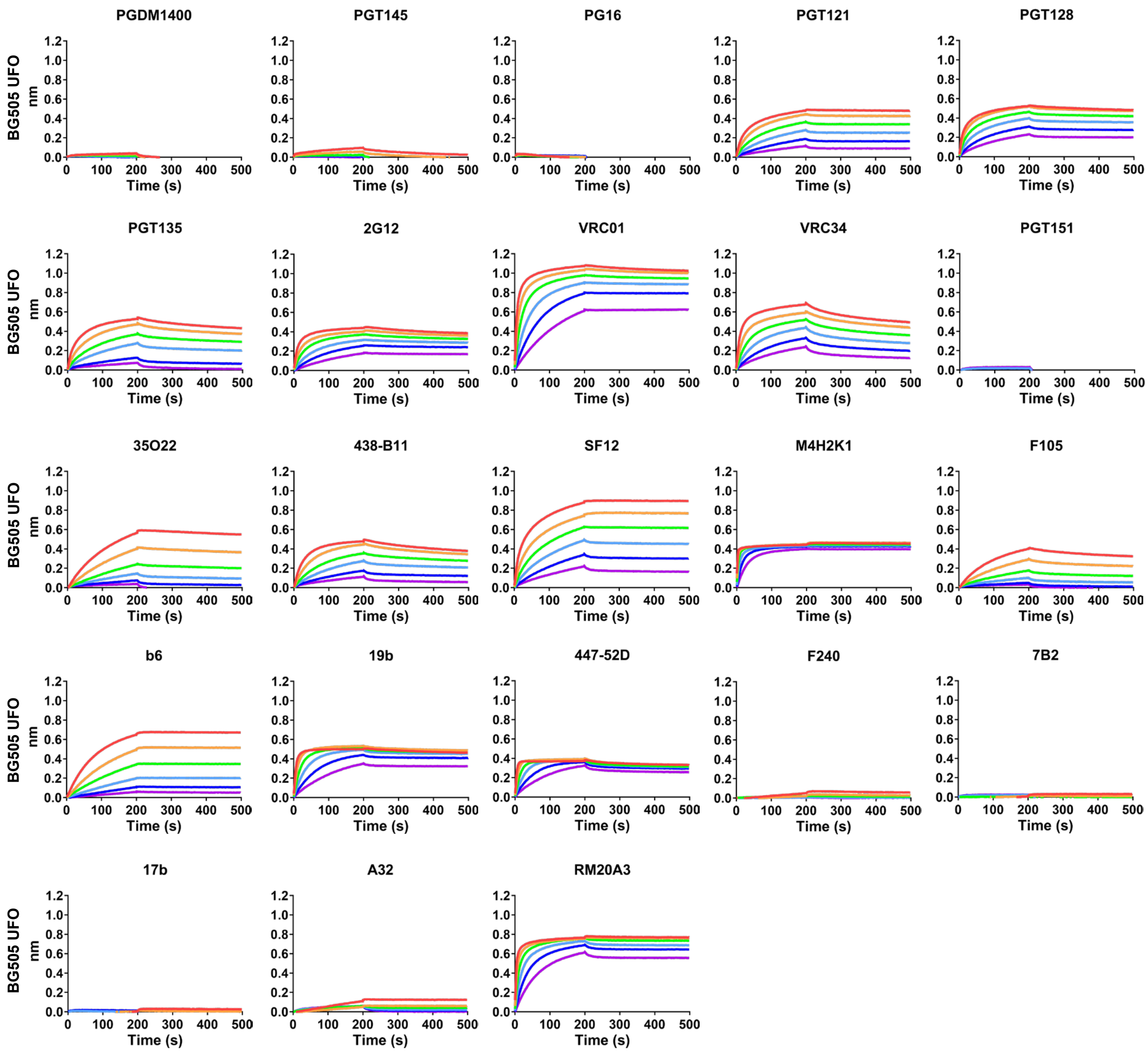

### Fig. S2

e

Hi-produced, glycan-trimmed BG505 UFO-E2p-L4P SApNP (AHQ)

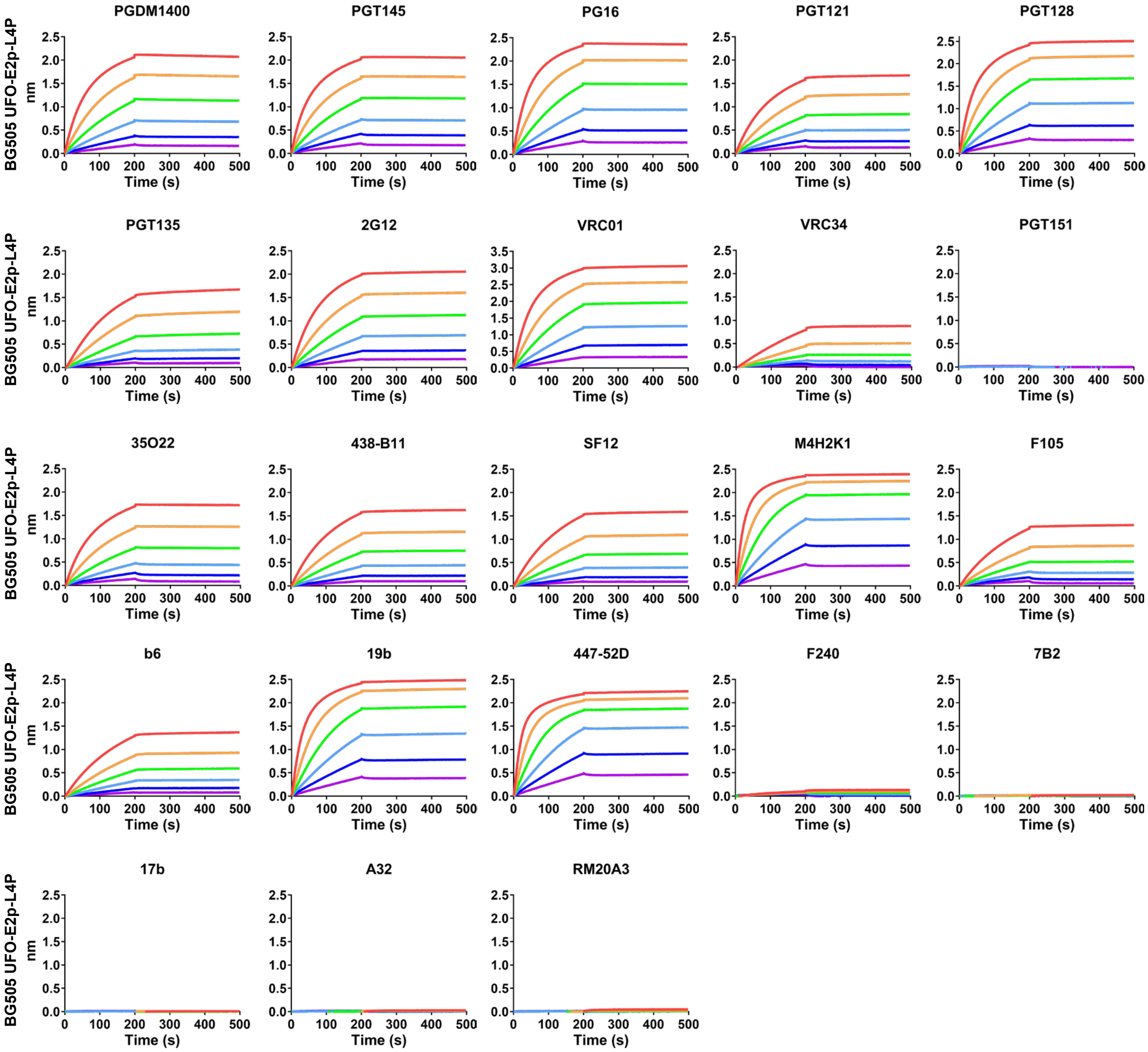

### Fig. S2

f

Hi5-produced, glycan-trimmed BG505 UFO-10GS-I3-01v9-L7P SApNP (AHQ)

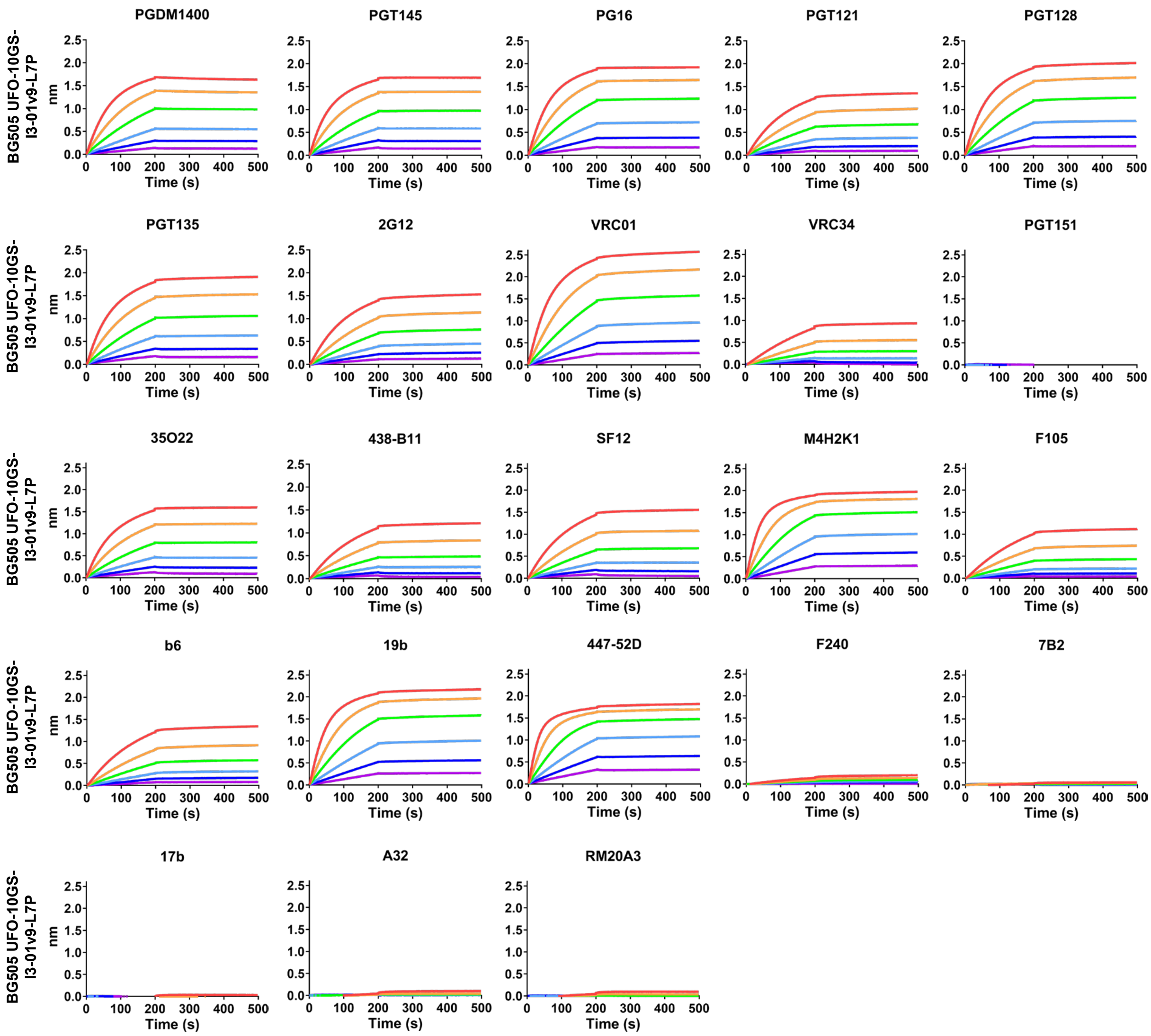

**Fig. S2**

g

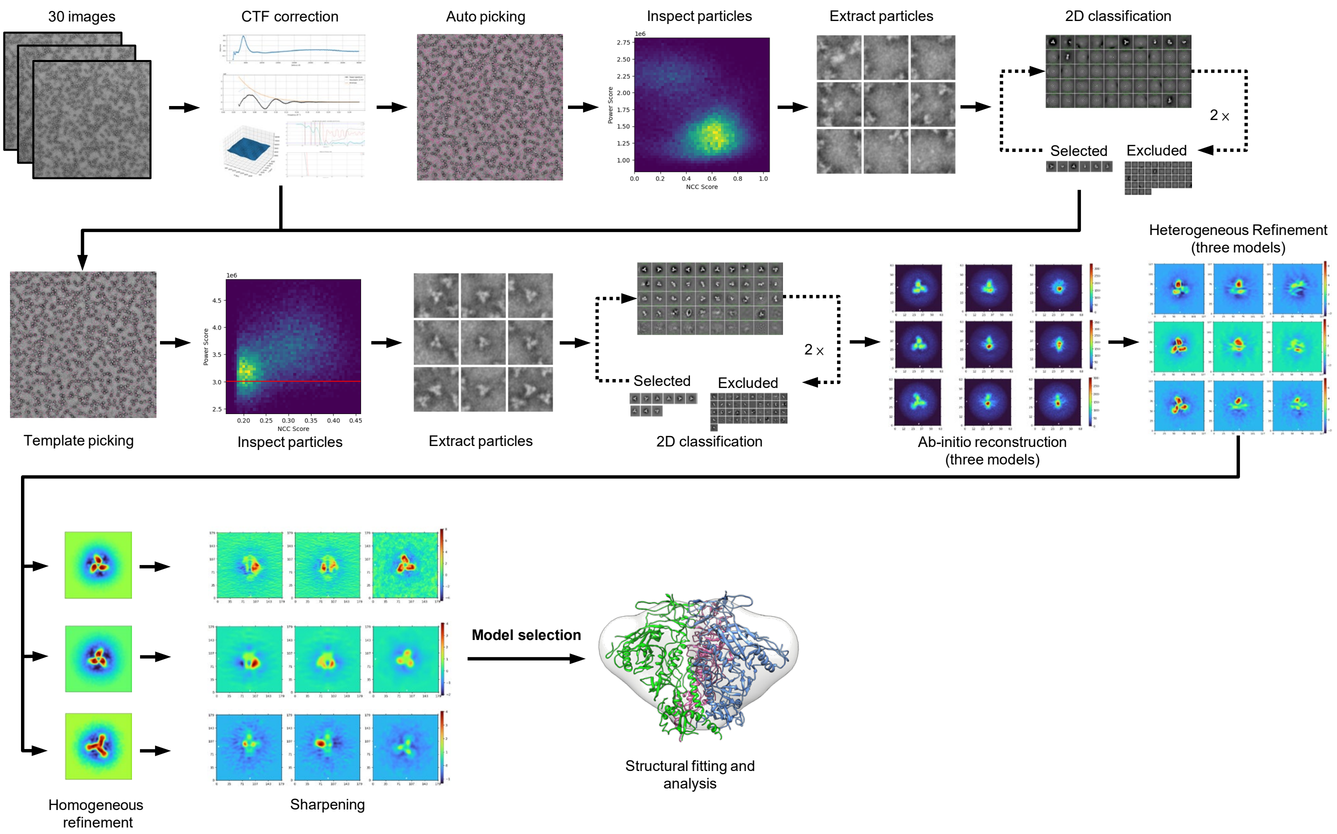

30 images
CTF correction
Auto picking
Inspect particles
Extract particles
2D classification
2 ×
Selected
Excluded
NCC Score
Power Score
Template picking
Inspect particles
Extract particles
2D classification
Ab-initio reconstruction
(three models)
Heterogeneous Refinement
(three models)
Homogeneous
refinement
Sharpening
Model selection
Structural fitting and
analysis


h

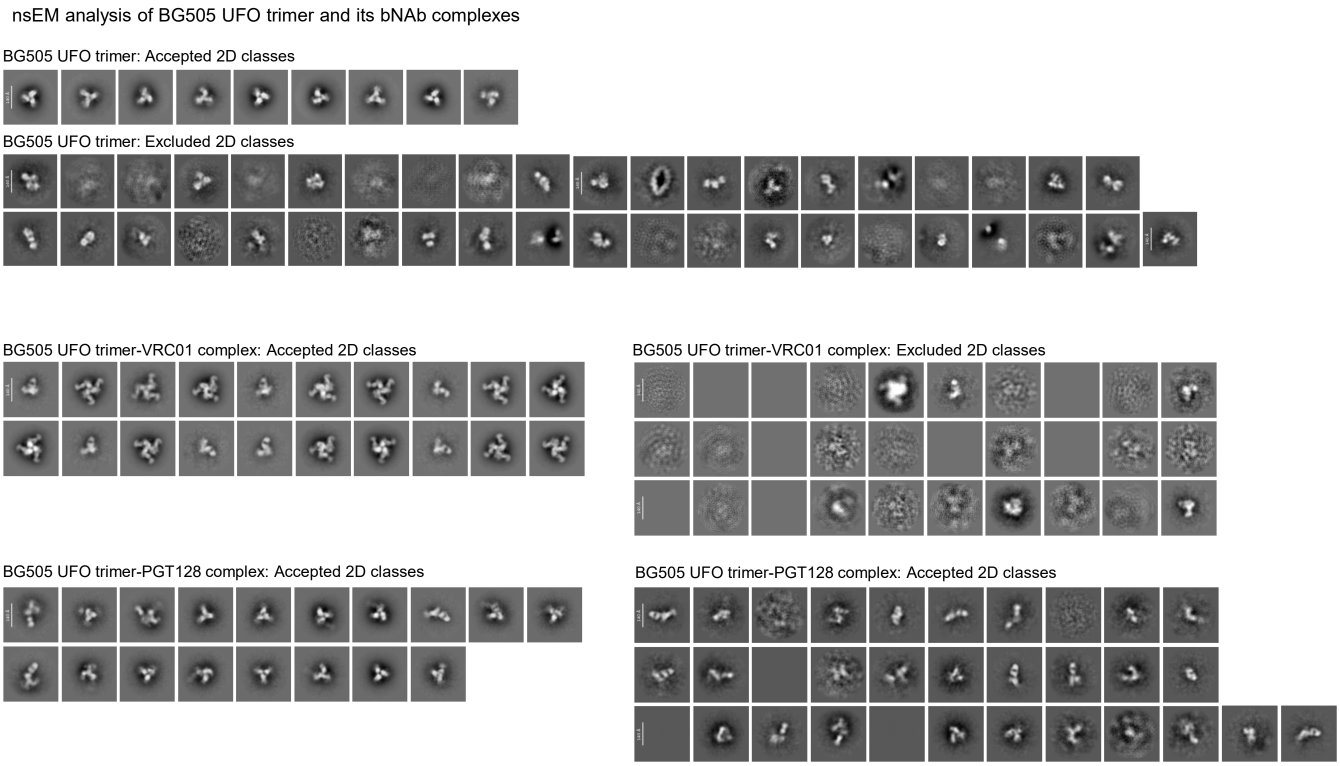

nsEM analysis of BG505 UFO trimer and its bNAb complexes
BG505 UFO trimer: Accepted 2D classes
BG505 UFO trimer: Excluded 2D classes
BG505 UFO trimer-VRC01 complex: Accepted 2D classes
BG505 UFO trimer-VRC01 complex: Excluded 2D classes
BG505 UFO trimer-PGT128 complex: Accepted 2D classes
BG505 UFO trimer-PGT128 complex: Accepted 2D classes

### Fig. S2

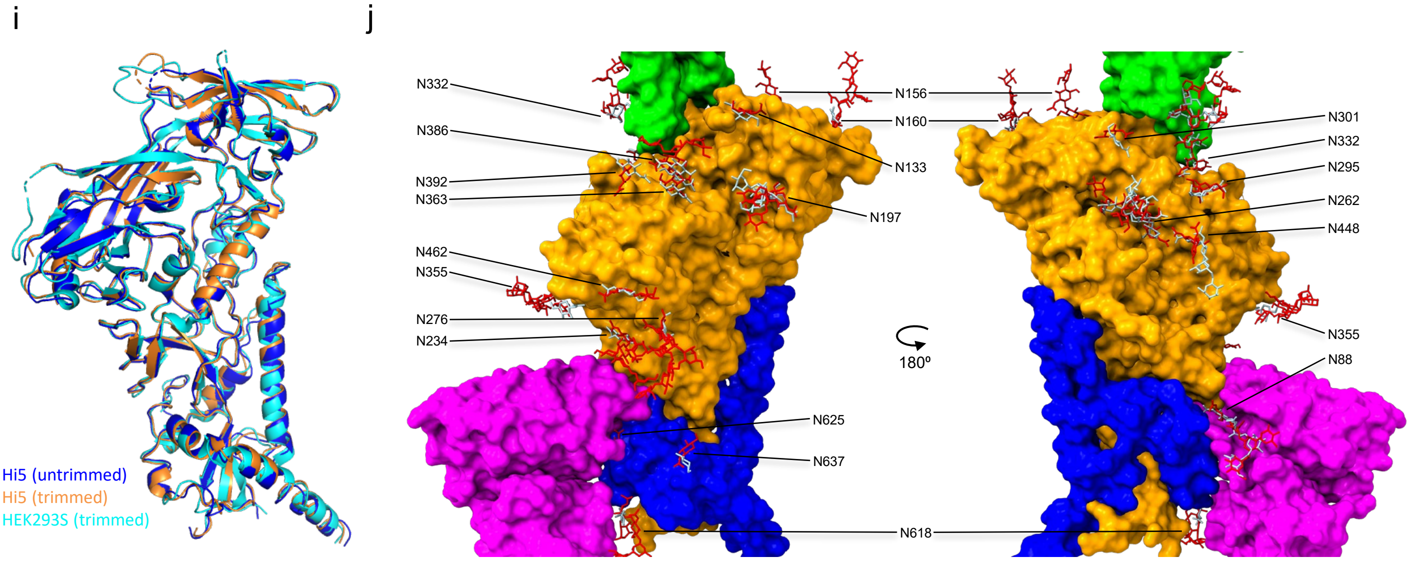


**Fig. S2. Antigenic and structural characterization of Hi5-produced BG505 Env immunogens.** (**a**–**f**) BLI analysis of Hi5-produced BG505 Env immunogens carrying (**a–c**) unmodified or (**d–f**) Endo H–trimmed glycans for binding to a panel of 23 HIV-1 antibodies including NAbs, bNAbs, and non-NAbs. A two-fold concentration gradient of antigen, starting at 266 nM for the trimer and 5.6 nM for the two SApNPs, was used in a six-point dilution series. (**g**) Flowchart illustrating image processing, 2D classification, and 3D reconstruction of negative-stain EM (nsEM) data for Hi5-produced BG505 UFO trimers and bNAb complexes using CryoSPARC. (**h**) nsEM 2D classes selected for or excluded from 3D reconstruction for the BG505 UFO trimer, trimer–VRC01 complex, and trimer–PGT128 complex. (**i**) Cα superposition of BG505 UFO trimer structures derived from Hi5 cells without Endo H treatment (blue), Hi5 cells with Endo H treatment (orange), and HEK293S cells with Endo H treatment (cyan). (**j**) Surface representation of a BG505 UFO protomer with N-linked glycans modeled from electron density and mapped onto the protomer (red, glycans from the untrimmed Hi5 structure; gray, glycans from the Endo H–treated Hi5 structure). gp120 is shown in gold, gp41 in blue, 35O22 in purple, and 3H109L/PGT124 in green.

### Fig. S3

a Purified week 11 IgG from mice immunized with Hi5-produced BG505 Env immunogens exhibits tier 2 neutralization

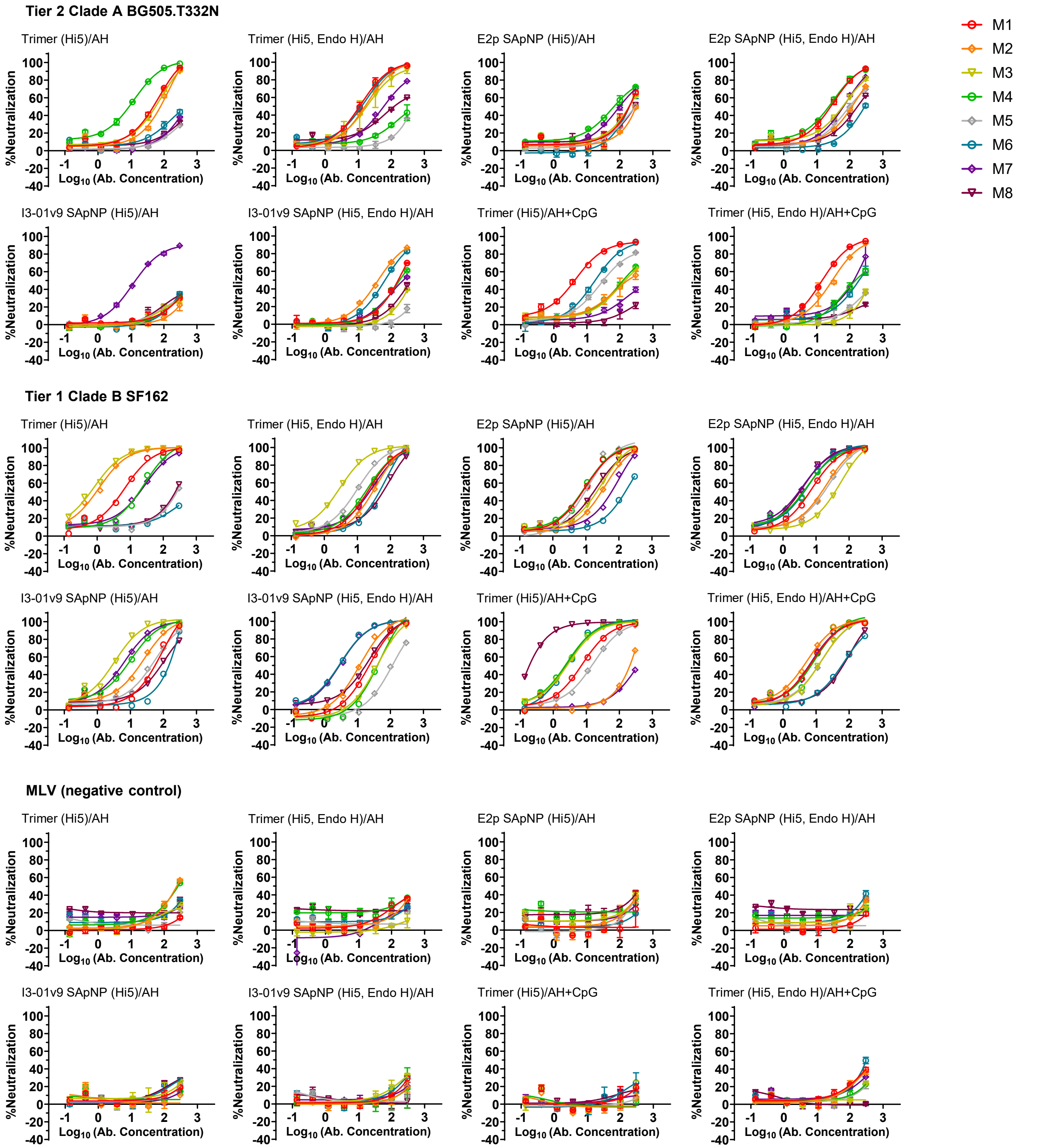

Fig. S3

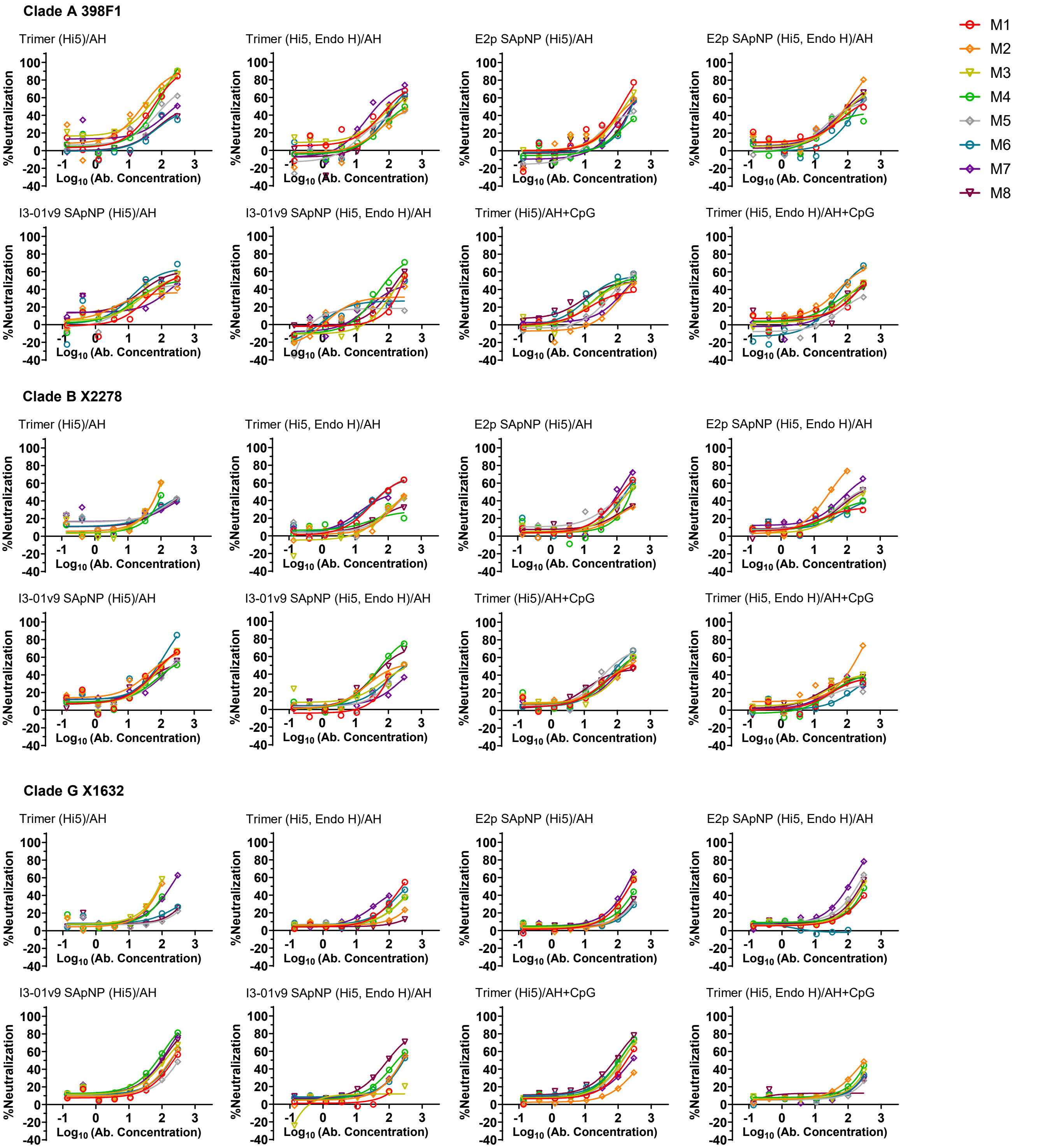
M1
M2
M3
M4
M5
M6
M7
M8
Clade A 398F1
Trimer (Hi5)/AH
Trimer (Hi5, Endo H)/AH
E2p SApNP (Hi5)/AH
E2p SApNP (Hi5, Endo H)/AH
I3-01v9 SApNP (Hi5)/AH
I3-01v9 SApNP (Hi5, Endo H)/AH
Trimer (Hi5)/AH+CpG
Trimer (Hi5, Endo H)/AH+CpG
Clade B X2278
Trimer (Hi5)/AH
Trimer (Hi5, Endo H)/AH
E2p SApNP (Hi5)/AH
E2p SApNP (Hi5, Endo H)/AH
I3-01v9 SApNP (Hi5)/AH
I3-01v9 SApNP (Hi5, Endo H)/AH
Trimer (Hi5)/AH+CpG
Trimer (Hi5, Endo H)/AH+CpG
Clade G X1632
Trimer (Hi5)/AH
Trimer (Hi5, Endo H)/AH
E2p SApNP (Hi5)/AH
E2p SApNP (Hi5, Endo H)/AH
I3-01v9 SApNP (Hi5)/AH
I3-01v9 SApNP (Hi5, Endo H)/AH
Trimer (Hi5)/AH+CpG
Trimer (Hi5, Endo H)/AH+CpG
%Neutralization
$Log_{10}$ (Ab. Concentration)
100
80
60
40
20
0
-20
-40
-1
0
1
2
3

Fig. S3

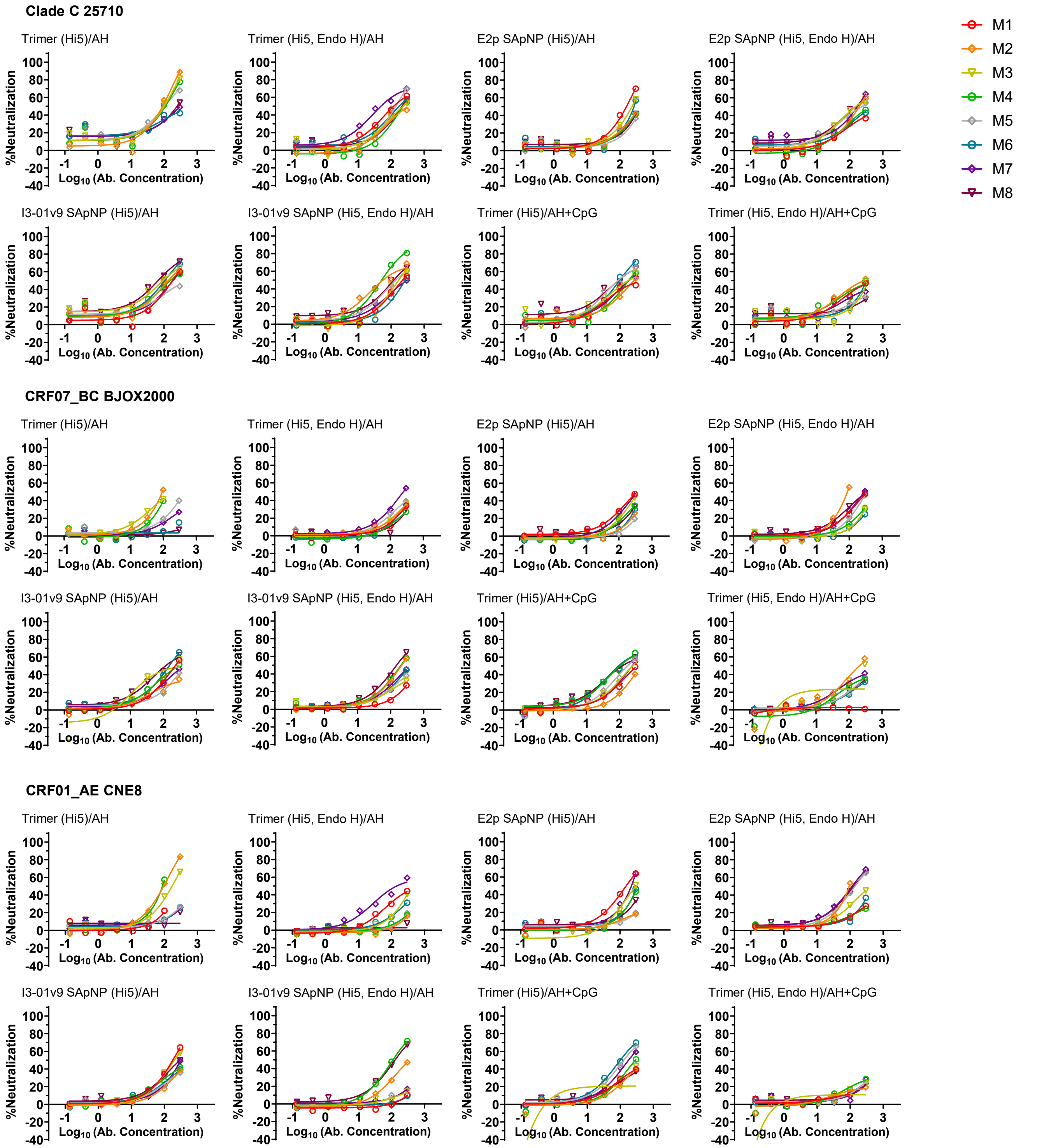

Clade C 25710
Trimer (Hi5)/AH
Trimer (Hi5, Endo H)/AH
E2p SApNP (Hi5)/AH
E2p SApNP (Hi5, Endo H)/AH
I3-01v9 SApNP (Hi5)/AH
I3-01v9 SApNP (Hi5, Endo H)/AH
Trimer (Hi5)/AH+CpG
Trimer (Hi5, Endo H)/AH+CpG
CRF07_BC BJOX2000
Trimer (Hi5)/AH
Trimer (Hi5, Endo H)/AH
E2p SApNP (Hi5)/AH
E2p SApNP (Hi5, Endo H)/AH
I3-01v9 SApNP (Hi5)/AH
I3-01v9 SApNP (Hi5, Endo H)/AH
Trimer (Hi5)/AH+CpG
Trimer (Hi5, Endo H)/AH+CpG
CRF01_AE CNE8
Trimer (Hi5)/AH
Trimer (Hi5, Endo H)/AH
E2p SApNP (Hi5)/AH
E2p SApNP (Hi5, Endo H)/AH
I3-01v9 SApNP (Hi5)/AH
I3-01v9 SApNP (Hi5, Endo H)/AH
Trimer (Hi5)/AH+CpG
Trimer (Hi5, Endo H)/AH+CpG
%Neutralization
Log10 (Ab. Concentration)
M1
M2
M3
M4
M5
M6
M7
M8

### Fig. S3

b

Neutralization $IC_{50}$ values of purified IgG

| W11 BG505 T332N | M1 | M2 | M3 | M4 | M5 | M6 | M7 | M8 | RR | Geomean |
|---|---|---|---|---|---|---|---|---|---|---|
| Trimer (Hi5)/AH | 41.54 | 67.24 | 48.54 | 7.492 | 600 | 600 | 600 | 600 | 50% | 138.0 |
| Trimer (Hi5, Endo H)/AH | 10.59 | 15.29 | 17.47 | 600 | 600 | 11.75 | 57.74 | 108.1 | 75% | 54.2 |
| E2p SApNP (Hi5)/AH | 176.2 | 600 | 174.6 | 63.16 | 265 | 205.5 | 95.76 | 247.4 | 88% | 187.1 |
| E2p SApNP (Hi5, Endo H)/AH | 24.13 | 105.1 | 53.26 | 20.16 | 96.19 | 276.8 | 58.86 | 153.4 | 100% | 71.1 |
| I3-01v9 SApNP (Hi5)/AH | 600 | 600 | 600 | 600 | 600 | 600 | 15.12 | 600 | 13% | 378.7 |
| I3-01v9 SApNP (Hi5, Endo H)/AH | 158.5 | 40.8 | 600 | 181.8 | 600 | 65.39 | 211.5 | 600 | 63% | 208.1 |
| Trimer (Hi5)/AH+CpG | 4.102 | 140.4 | 130.4 | 113.7 | 33.19 | 17.85 | 600 | 600 | 75% | 80.8 |
| Trimer (Hi5, Endo H)/AH+CpG | 14.85 | 29.45 | 600 | 150.4 | 600 | 176.7 | 108.7 | 600 | 63% | 151.2 |

| W11 SF162 | M1 | M2 | M3 | M4 | M5 | M6 | M7 | M8 | RR | Geomean |
|---|---|---|---|---|---|---|---|---|---|---|
| Trimer (Hi5)/AH | 5.297 | 0.8824 | 0.6007 | 16.36 | 230.1 | 600 | 16.95 | 198.7 | 88% | 19.6 |
| Trimer (Hi5, Endo H)/AH | 19.61 | 27.39 | 2.344 | 16.87 | 8.452 | 41.09 | 21.19 | 52.29 | 100% | 17.3 |
| E2p SApNP (Hi5)/AH | 7.29 | 26.44 | 18.67 | 6.845 | 8.898 | 143.2 | 50.91 | 15.92 | 100% | 20.0 |
| E2p SApNP (Hi5, Endo H)/AH | 5.988 | 12.98 | 32.06 | 3.598 | 15.07 | 3.831 | 2.612 | 2.919 | 100% | 6.7 |
| I3-01v9 SApNP (Hi5)/AH | 48.4 | 16.32 | 2.536 | 6.628 | 38.39 | 118.6 | 4.939 | 74.05 | 100% | 19.6 |
| I3-01v9 SApNP (Hi5, Endo H)/AH | 23.29 | 11.8 | 40.77 | 40.49 | 122.6 | 2.677 | 2.799 | 15.94 | 100% | 16.9 |
| Trimer (Hi5)/AH+CpG | 6.739 | 201 | 2.443 | 2.135 | 13.7 | 2.41 | 600 | 0.1831 | 88% | 8.4 |
| Trimer (Hi5, Endo H)/AH+CpG | 5.875 | 3.854 | 10.76 | 6.375 | 10.66 | 54.97 | 5.696 | 55.6 | 100% | 11.4 |

| W11 MLV | M1 | M2 | M3 | M4 | M5 | M6 | M7 | M8 | RR | Geomean |
|---|---|---|---|---|---|---|---|---|---|---|
| Trimer (Hi5)/AH | 600 | 236.3 | 600 | 239.3 | 600 | 600 | 600 | 600 | 25% | 476.1 |
| Trimer (Hi5, Endo H)/AH | 600 | 600 | 600 | 600 | 600 | 600 | 600 | 600 | 0% | 600.0 |
| E2p SApNP (Hi5)/AH | 600 | 600 | 600 | 600 | 600 | 600 | 600 | 600 | 0% | 600.0 |
| E2p SApNP (Hi5, Endo H)/AH | 600 | 600 | 600 | 600 | 600 | 600 | 600 | 600 | 0% | 600.0 |
| I3-01v9 SApNP (Hi5)/AH | 600 | 600 | 600 | 600 | 600 | 600 | 600 | 600 | 0% | 600.0 |
| I3-01v9 SApNP (Hi5, Endo H)/AH | 600 | 600 | 600 | 600 | 600 | 600 | 600 | 600 | 0% | 600.0 |
| Trimer (Hi5)/AH+CpG | 600 | 600 | 600 | 600 | 600 | 600 | 600 | 600 | 0% | 600.0 |
| Trimer (Hi5, Endo H)/AH+CpG | 600 | 600 | 600 | 600 | 600 | 600 | 600 | 600 | 0% | 600.0 |

| W11 398F1 | M1 | M2 | M3 | M4 | M5 | M6 | M7 | M8 | RR | Geomean |
|---|---|---|---|---|---|---|---|---|---|---|
| Trimer (Hi5)/AH | 53.48 | 25.13 | 31.71 | 59.15 | 112.1 | 600 | 185.9 | 600 | 75% | 108.3 |
| Trimer (Hi5, Endo H)/AH | 99.19 | 600 | 117.7 | 600 | 600 | 145.6 | 55.94 | 169.8 | 63% | 207.9 |
| E2p SApNP (Hi5)/AH | 121.9 | 178.2 | 160.5 | 600 | 600 | 290 | 250.1 | 279.6 | 75% | 266.5 |
| E2p SApNP (Hi5, Endo H)/AH | 600 | 76.13 | 116 | 600 | 120.3 | 207.3 | 108.4 | 102 | 75% | 174.9 |
| I3-01v9 SApNP (Hi5)/AH | 150.8 | 600 | 128.7 | 136.7 | 600 | 58.53 | 600 | 70.37 | 63% | 198.0 |
| I3-01v9 SApNP (Hi5, Endo H)/AH | 600 | 600 | 261.5 | 104.7 | 600 | 600 | 248.2 | 187.5 | 50% | 336.7 |
| Trimer (Hi5)/AH+CpG | 600 | 600 | 600 | 125.8 | 166.5 | 89.89 | 600 | 111.9 | 50% | 268.9 |
| Trimer (Hi5, Endo H)/AH+CpG | 600 | 101.4 | 600 | 600 | 600 | 125.1 | 600 | 600 | 25% | 394.9 |

| W11 X2278 | M1 | M2 | M3 | M4 | M5 | M6 | M7 | M8 | RR | Geomean |
|---|---|---|---|---|---|---|---|---|---|---|
| Trimer (Hi5)/AH | N/A | 75.52 | 80.95 | 126 | 600 | 600 | 600 | 600 | 43% | 268.2 |
| Trimer (Hi5, Endo H)/AH | 86.63 | 600 | 600 | 600 | 600 | 82.01 | 71.91 | 600 | 38% | 281.8 |
| E2p SApNP (Hi5)/AH | 138.3 | 600 | 205.8 | 292.7 | 164.2 | 177.5 | 110.2 | 600 | 75% | 236.0 |
| E2p SApNP (Hi5, Endo H)/AH | 600 | 29.75 | 600 | 600 | 158.8 | 600 | 97.76 | 174.7 | 50% | 238.5 |
| I3-01v9 SApNP (Hi5)/AH | 93.11 | 73.52 | 102.8 | 168.2 | 166.4 | 61.59 | 165.3 | 143.3 | 100% | 114.1 |
| I3-01v9 SApNP (Hi5, Endo H)/AH | 199.5 | 146.9 | 219.7 | 65.14 | 208.1 | 236 | 600 | 83.05 | 88% | 178.4 |
| Trimer (Hi5)/AH+CpG | 600 | 145.8 | 153.4 | 127.2 | 75.44 | 101.8 | 146.5 | 600 | 75% | 181.0 |
| Trimer (Hi5, Endo H)/AH+CpG | 600 | 117.5 | 600 | 600 | 600 | 600 | 600 | 600 | 13% | 489.4 |

| W11 X1632 | M1 | M2 | M3 | M4 | M5 | M6 | M7 | M8 | RR | Geomean |
|---|---|---|---|---|---|---|---|---|---|---|
| Trimer (Hi5)/AH | N/A | 89.24 | 74.33 | 146.1 | 600 | 600 | 162.9 | 600 | 57% | 230.0 |
| Trimer (Hi5, Endo H)/AH | 227.7 | 600 | 600 | 600 | 600 | 600 | 117.1 | 600 | 25% | 433.4 |
| E2p SApNP (Hi5)/AH | 228.6 | 600 | 221 | 600 | 600 | 600 | 167.6 | 600 | 38% | 400.2 |
| E2p SApNP (Hi5, Endo H)/AH | 600 | N/A | 253.1 | 600 | 163.7 | 600 | 88.42 | 215.1 | 50% | 289.4 |
| I3-01v9 SApNP (Hi5)/AH | 198.6 | 157.3 | 105.9 | 60.71 | 600 | 158 | 79.63 | 87.63 | 88% | 138.2 |
| I3-01v9 SApNP (Hi5, Endo H)/AH | 600 | 245.2 | 600 | 166 | N/A | 254.3 | 236.7 | 89.12 | 63% | 259.2 |
| Trimer (Hi5)/AH+CpG | 166.1 | 600 | 120.3 | 98.15 | 124.8 | 93.2 | 220.5 | 69 | 88% | 146.2 |
| Trimer (Hi5, Endo H)/AH+CpG | N/A | 600 | 600 | 600 | 600 | 600 | 600 | 600 | 0% | 600.0 |

| W11 25710 | M1 | M2 | M3 | M4 | M5 | M6 | M7 | M8 | RR | Geomean |
|---|---|---|---|---|---|---|---|---|---|---|
| Trimer (Hi5)/AH | N/A | 78.81 | 84.53 | 92.83 | 92.62 | 600 | 600 | 187.2 | 71% | 168.5 |
| Trimer (Hi5, Endo H)/AH | 120.3 | 600 | 201.9 | 243.2 | 136.8 | 159.4 | 54.71 | 201 | 88% | 174.2 |
| E2p SApNP (Hi5)/AH | 151.6 | 600 | 226 | 600 | 600 | 292 | 600 | 600 | 38% | 408.7 |
| E2p SApNP (Hi5, Endo H)/AH | 600 | 188.1 | 137.7 | 600 | 167.5 | 600 | 141.6 | 129.6 | 63% | 253.8 |
| I3-01v9 SApNP (Hi5)/AH | 170.2 | 123.2 | 83.14 | 133.1 | 600 | 95.55 | 161.2 | 60.11 | 88% | 137.7 |
| I3-01v9 SApNP (Hi5, Endo H)/AH | 210.4 | 68.19 | 173.1 | 53 | 133.4 | 265.3 | 600 | 109.8 | 88% | 153.4 |
| Trimer (Hi5)/AH+CpG | 600 | 222.1 | 149.6 | 171.8 | 94.9 | 93.23 | 162 | 84.63 | 88% | 159.3 |
| Trimer (Hi5, Endo H)/AH+CpG | 600 | 169 | 600 | 206.7 | 600 | 600 | 600 | 600 | 13% | 448.2 |

| W11 BJOX2000 | M1 | M2 | M3 | M4 | M5 | M6 | M7 | M8 | RR | Geomean |
|---|---|---|---|---|---|---|---|---|---|---|
| Trimer (Hi5)/AH | N/A | 104.6 | 111.9 | 167.5 | 600 | 600 | 600 | 600 | 43% | 306.5 |
| Trimer (Hi5, Endo H)/AH | 600 | 600 | 600 | 600 | 600 | 600 | 233.3 | 600 | 13% | 533.2 |
| E2p SApNP (Hi5)/AH | 600 | 600 | 600 | 600 | 600 | 600 | 600 | 600 | 0% | 600.0 |
| E2p SApNP (Hi5, Endo H)/AH | 600 | 104.6 | 600 | 600 | 600 | 600 | 273.8 | 600 | 25% | 437.3 |
| I3-01v9 SApNP (Hi5)/AH | 217.7 | 600 | 148.4 | 205.8 | 600 | 140.5 | 600 | 112.8 | 63% | 262.8 |
| I3-01v9 SApNP (Hi5, Endo H)/AH | 600 | 209.1 | 600 | 198.2 | 600 | 600 | 600 | 147 | 38% | 384.1 |
| Trimer (Hi5)/AH+CpG | 600 | 600 | 223.5 | 107 | 170.9 | 104.2 | 226.8 | 116.8 | 75% | 211.9 |
| Trimer (Hi5, Endo H)/AH+CpG | 600 | 198.4 | 233.7 | 600 | 600 | 600 | 600 | 600 | 25% | 464.4 |

| W11 CNE8 | M1 | M2 | M3 | M4 | M5 | M6 | M7 | M8 | RR | Geomean |
|---|---|---|---|---|---|---|---|---|---|---|
| Trimer (Hi5)/AH | 600 | 90.51 | 167.9 | 98.86 | 600 | 600 | 600 | 600 | 38% | 322.4 |
| Trimer (Hi5, Endo H)/AH | 600 | 600 | 600 | 600 | 600 | 600 | 130.4 | 600 | 13% | 495.8 |
| E2p SApNP (Hi5)/AH | 151.5 | 600 | 600 | 600 | 600 | 600 | 219.9 | 600 | 25% | 445.6 |
| E2p SApNP (Hi5, Endo H)/AH | 600 | 109.8 | 600 | 600 | 136.8 | 600 | 110.7 | 123.3 | 50% | 267.9 |
| I3-01v9 SApNP (Hi5)/AH | 178.8 | 600 | 204.4 | 600 | 600 | 600 | 600 | 600 | 25% | 450.8 |
| I3-01v9 SApNP (Hi5, Endo H)/AH | 600 | 600 | 600 | 115 | 600 | 600 | 600 | 126.6 | 25% | 401.8 |
| Trimer (Hi5)/AH+CpG | 600 | 600 | 600 | 269.9 | 127 | 101.9 | 172.6 | 600 | 50% | 306.6 |
| Trimer (Hi5, Endo H)/AH+CpG | 600 | 600 | 600 | 600 | 600 | 600 | 600 | 600 | 0% | 600.0 |

### Fig. S3

**Fig. S3. Immunogenicity of Hi5-produced BG505 Env immunogens in wild-type mice.** (**a**) Neutralization curves of purified week 11 IgG from individual mice immunized with Hi5-produced BG505 UFO trimer, E2p SApNP, and I3-01v9 SApNP vaccines (*n* = 8 mice per group) against a pseudovirus panel comprising the autologous tier 2 clade A BG505.T332N, the tier 1 clade B SF162, and six heterologous tier 2 strains: 398F1 (clade A), X2278 (clade B), X1632 (clade B), 25710 (clade C), BJOX2000 (CRF07_BC), and CNE8 (CRF01_AE). Pseudotyped murine leukemia virus (MLV) served as a negative control. (**b**) Summary of neutralization IC50 values of purified IgG. Color coding: green to red (low to high neutralization). Error bars represent the difference between duplicate measurements at each concentration for each sample. $IC_{50}$ values were calculated using GraphPad Prism.

### Fig. S4

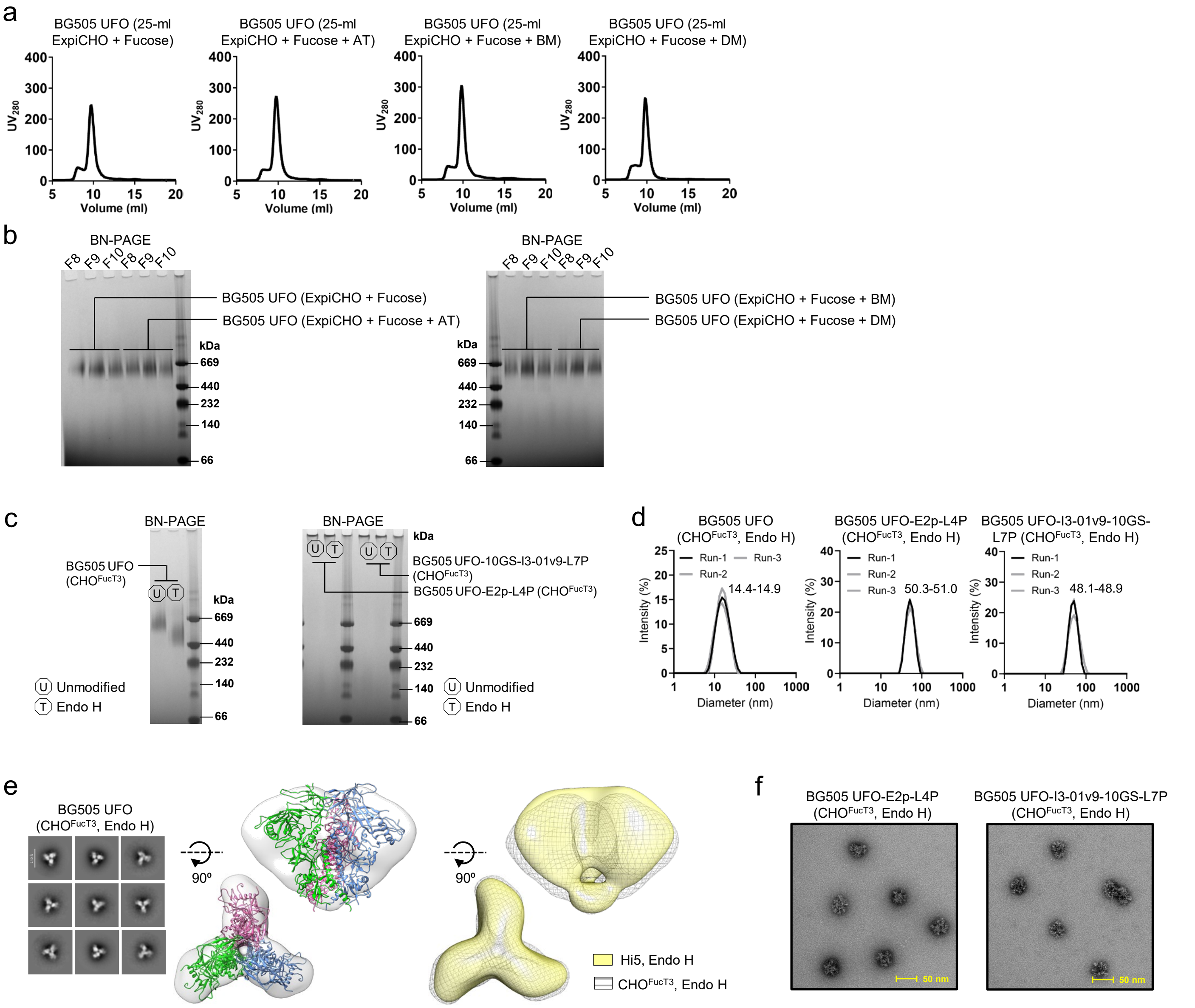

a
BG505 UFO (25-ml ExpiCHO + Fucose)
BG505 UFO (25-ml ExpiCHO + Fucose + AT)
BG505 UFO (25-ml ExpiCHO + Fucose + BM)
BG505 UFO (25-ml ExpiCHO + Fucose + DM)
UV280
Volume (ml)
b
BN-PAGE
F8 F9 F10 F8 F9 F10
BG505 UFO (ExpiCHO + Fucose)
BG505 UFO (ExpiCHO + Fucose + AT)
BG505 UFO (ExpiCHO + Fucose + BM)
BG505 UFO (ExpiCHO + Fucose + DM)
kDa
669
440
232
140
66
c
BN-PAGE
BG505 UFO (CHOFucT3)
BG505 UFO-10GS-I3-01v9-L7P (CHOFucT3)
BG505 UFO-E2p-L4P (CHOFucT3)
U Unmodified
T Endo H
d
BG505 UFO (CHOFucT3, Endo H)
BG505 UFO-E2p-L4P (CHOFucT3, Endo H)
BG505 UFO-I3-01v9-10GS-L7P (CHOFucT3, Endo H)
Run-1
Run-2
Run-3
14.4-14.9
50.3-51.0
48.1-48.9
Intensity (%)
Diameter (nm)
e
BG505 UFO (CHOFucT3, Endo H)
90º
Hi5, Endo H
CHOFucT3, Endo H
f
BG505 UFO-E2p-L4P (CHOFucT3, Endo H)
BG505 UFO-I3-01v9-10GS-L7P (CHOFucT3, Endo H)
50 nm

### Fig. S4

g

CHO$^{FucT3}$-produced BG505 UFO trimer (AHQ)

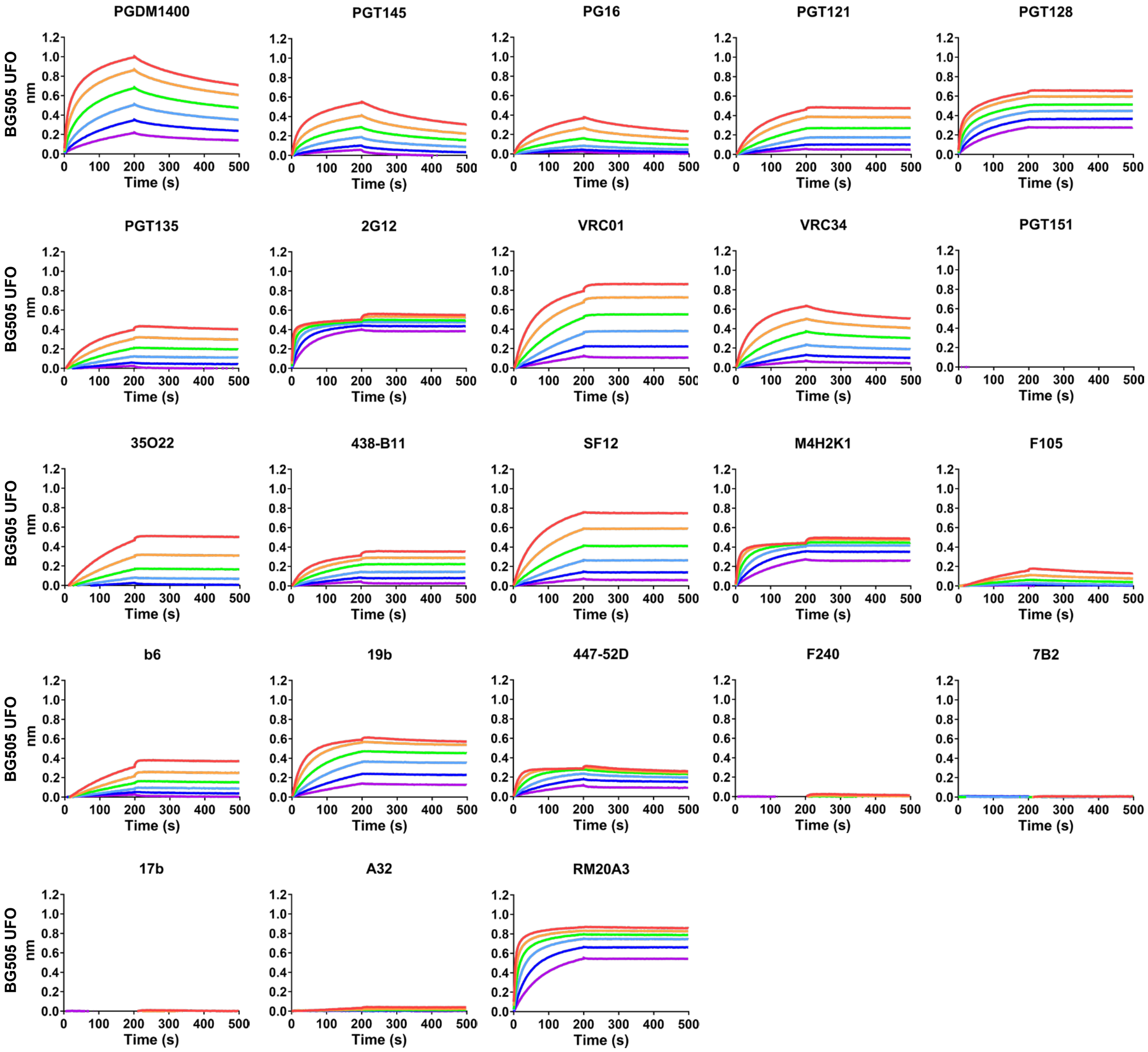

### Fig. S4

h

CHO$^{FucT3}$-produced BG505 UFO-E2p-L4P SApNP (AHQ)

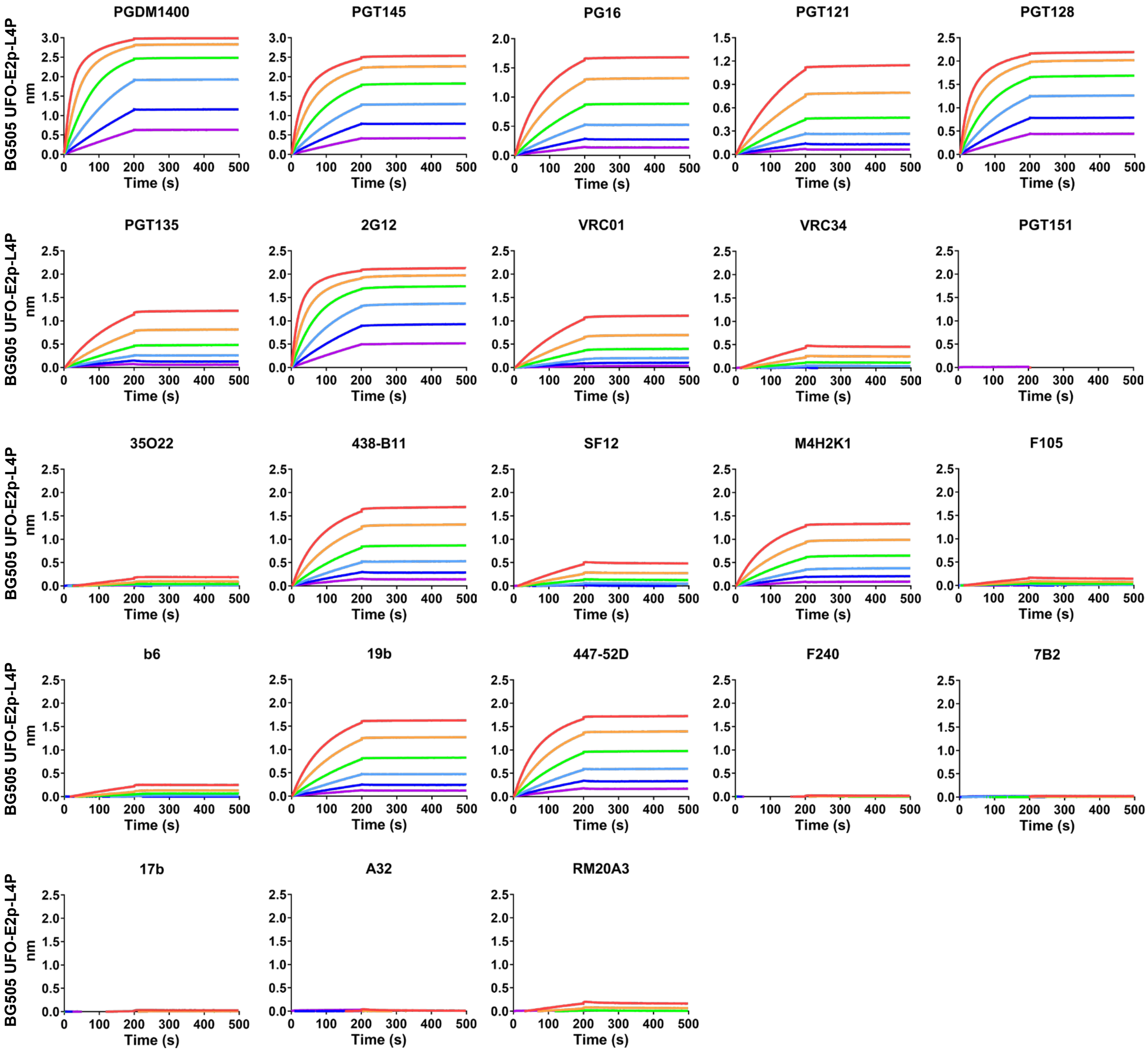

### Fig. S4

i

CHO$^{FucT3}$-produced BG505 UFO-10GS-I3-01v9-L7P SApNP (AHQ)

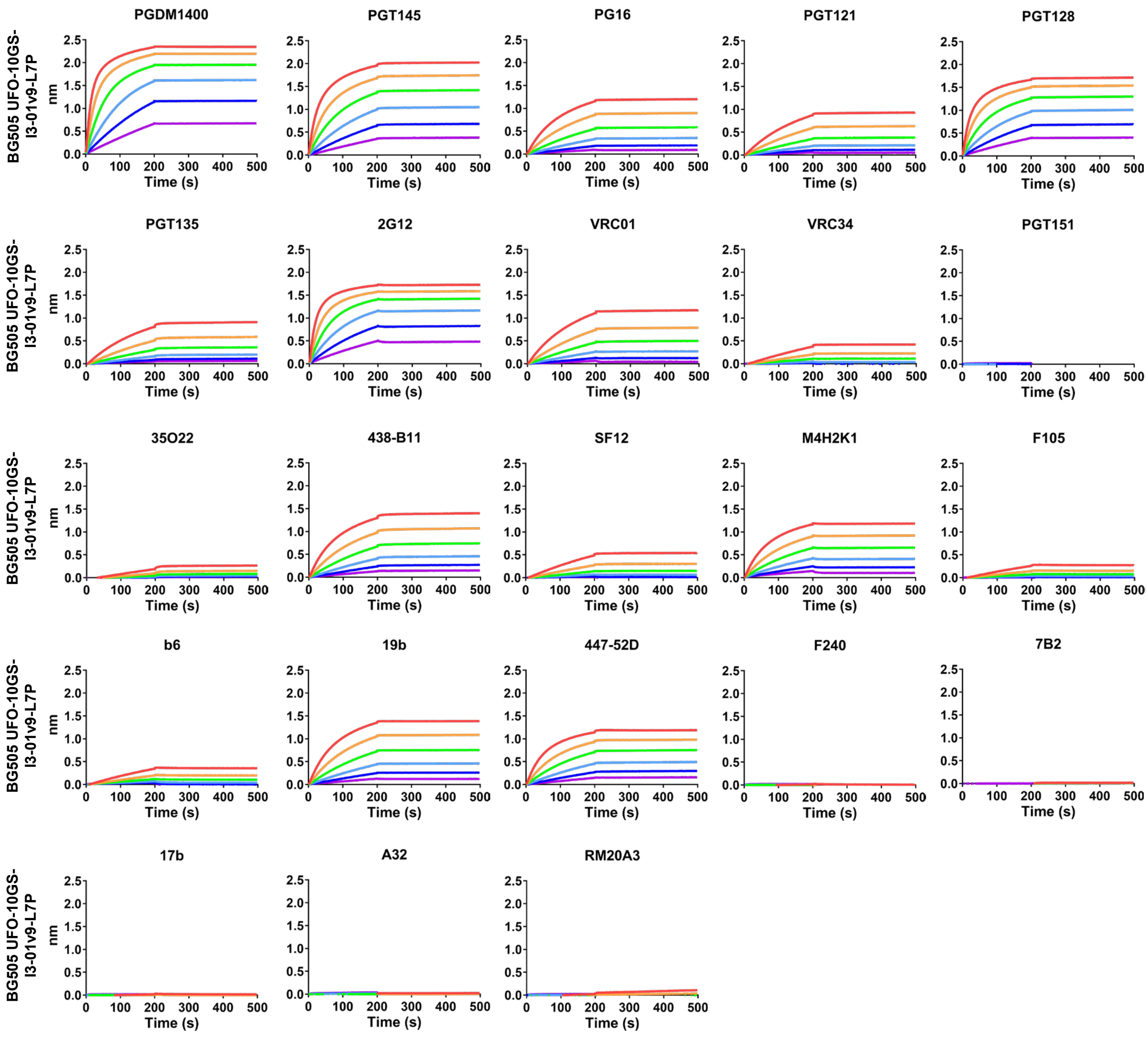

**Fig. S4**

j

CHO[FucT3]-produced, glycan-trimmed BG505 UFO trimer (AHQ)

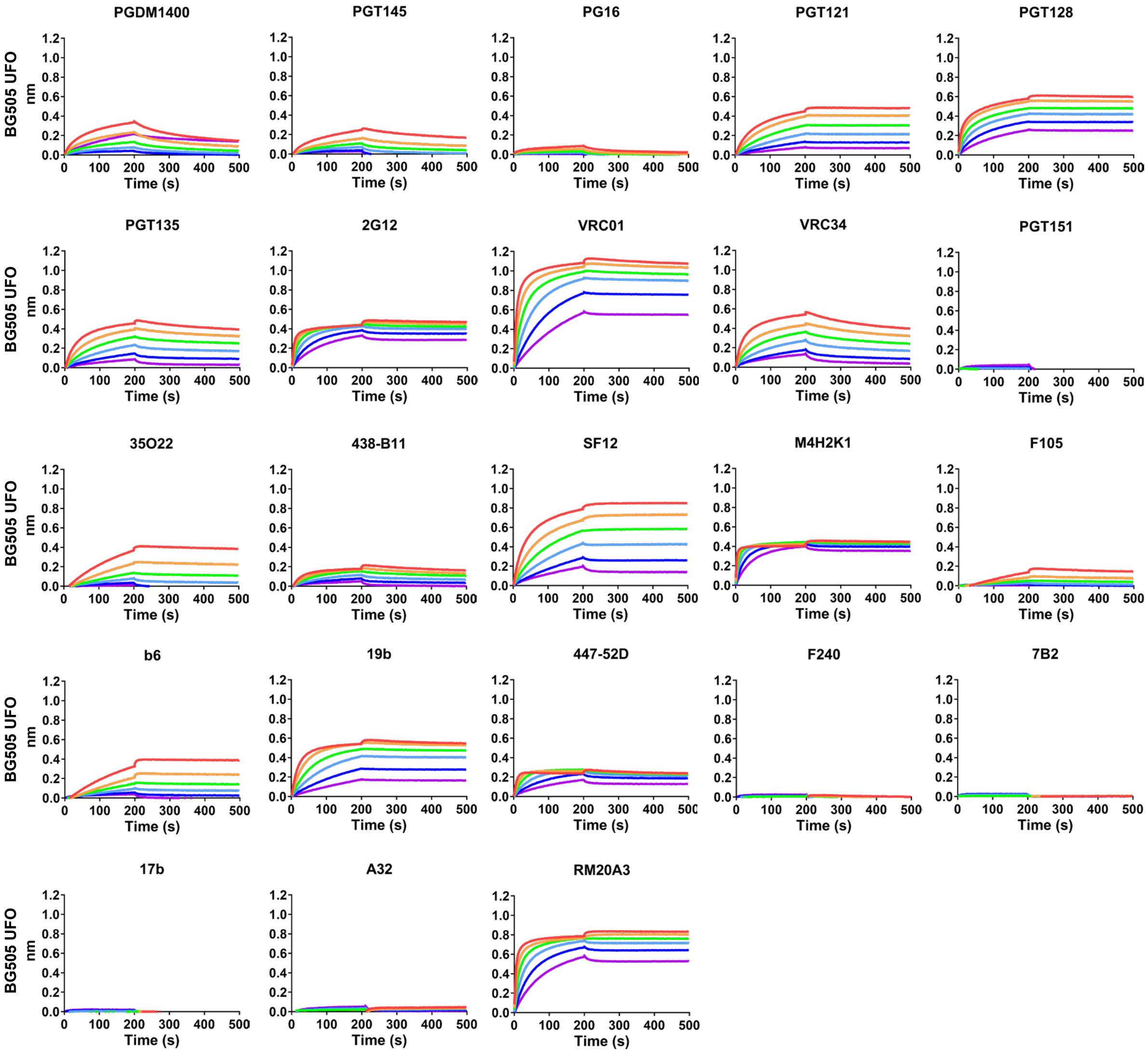

k

CHO$^{FucT3}$-produced, glycan-trimmed BG505 UFO-E2p-L4P SApNP (AHQ)

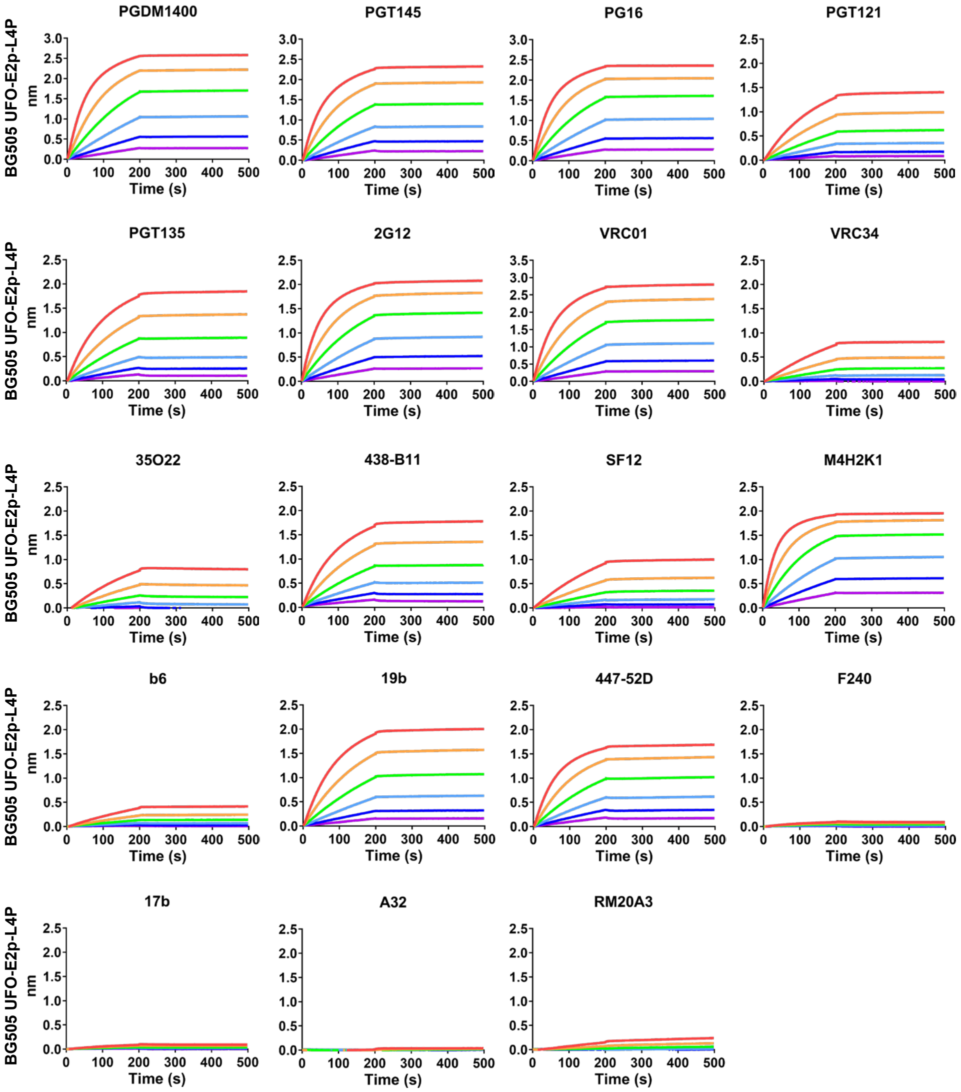

**Fig. S4**

I

CHO$^{FucT3}$-produced, glycan-trimmed BG505 UFO-10GS-I3-01v9-L7P SApNP (AHQ)

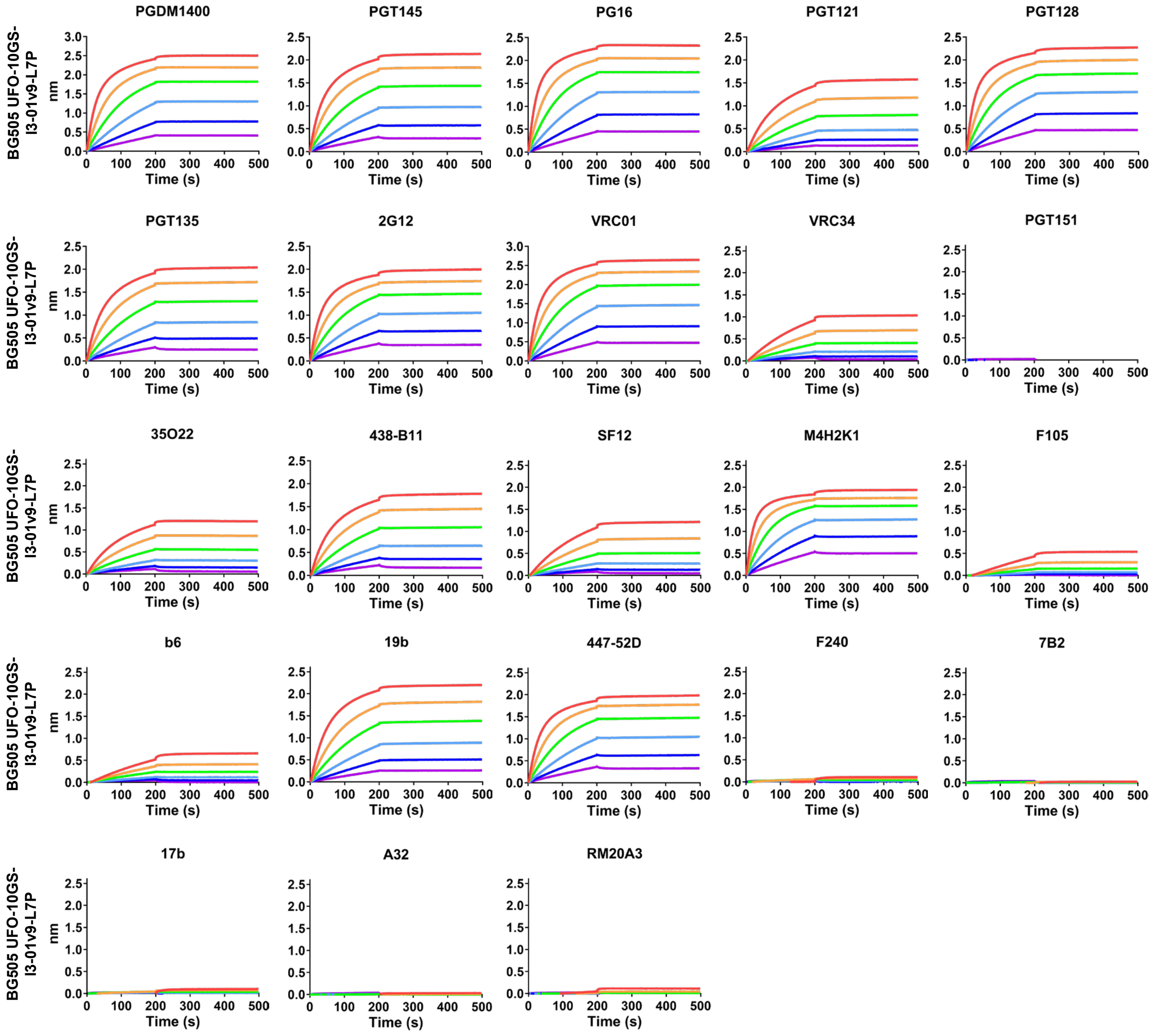

**Fig. S4. In vitro characterization of glycoengineered CHO$^{FucT3}$-produced BG505 Env immunogens.** (**a**) SEC profiles of BG505 UFO trimers transiently expressed in the ExpiCHO system in the presence of L-fucose alone or L-fucose in combination with one of the core α(1,3)-fucosyltransferases (FucT) from *Arabidopsis thaliana* (AT, plant), *Bombyx mori* (BM, insect), and *Drosophila melanogaster* (DM, insect). (**b**) BN-PAGE analysis of three SEC fractions (8-10) for BG505 UFO trimers transiently expressed in the ExpiCHO system in the presence of L-fucose alone or L-fucose in combination with one of the core α(1,3)-fucosyltransferases (AT, BM, and DM). (**c**) BN-PAGE analysis of CHO$^{FucT3}$-produced BG505 UFO trimers and SApNPs (E2p and I3-01v9) carrying unmodified or Endo H-trimmed glycans. CHO$^{FucT3}$ indicates a glycoengineered mammalian expression system in which a glycoprotein is co-expressed with three FucTs (AT, BM, and DM) in ExpiCHO cells supplemented with L-fucose. (**d**) Particle size distributions of CHO$^{FucT3}$-produced, glycan-trimmed BG505 UFO trimers and SApNPs. DLS measurements from three independent production batches are shown for each construct, with hydrodynamic diameters (Dh) indicated. (**e**) nsEM analysis of CHO$^{FucT3}$-produced, glycan-trimmed BG505 UFO trimers. Left: representative 2D class averages; middle: side (top) and top (bottom) views of the 3D reconstruction with the crystal structure of the BG505 UFO trimer (PDB: 6UTK) fitted into the density map; right: side (top) and top (bottom) views of superimposed 3D reconstructions derived from Hi5- and CHO$^{FucT3}$-produced, glycan-trimmed BG505 UFO trimers. EM density color-coding scheme: Hi5 (yellow) and CHO$^{FucT3}$ (black mesh) (**f**) nsEM images of CHO$^{FucT3}$-produced, glycan-trimmed BG505 UFO trimer-presenting E2p and I3-01v9 SApNPs. Scale bar: 50 nm. (**g–l**) BLI analysis of CHO$^{FucT3}$-produced BG505 Env immunogens carrying (**g–i**) unmodified or (**j–l**) Endo H–trimmed glycans binding to a panel of 23 HIV-1 antibodies, including NAbs, bNAbs, and non-NAbs. A twofold concentration gradient of antigen, starting at 266 nM for the trimer and 5.6 nM for the two SApNPs, was used in a six-point dilution series.

### Fig. S5

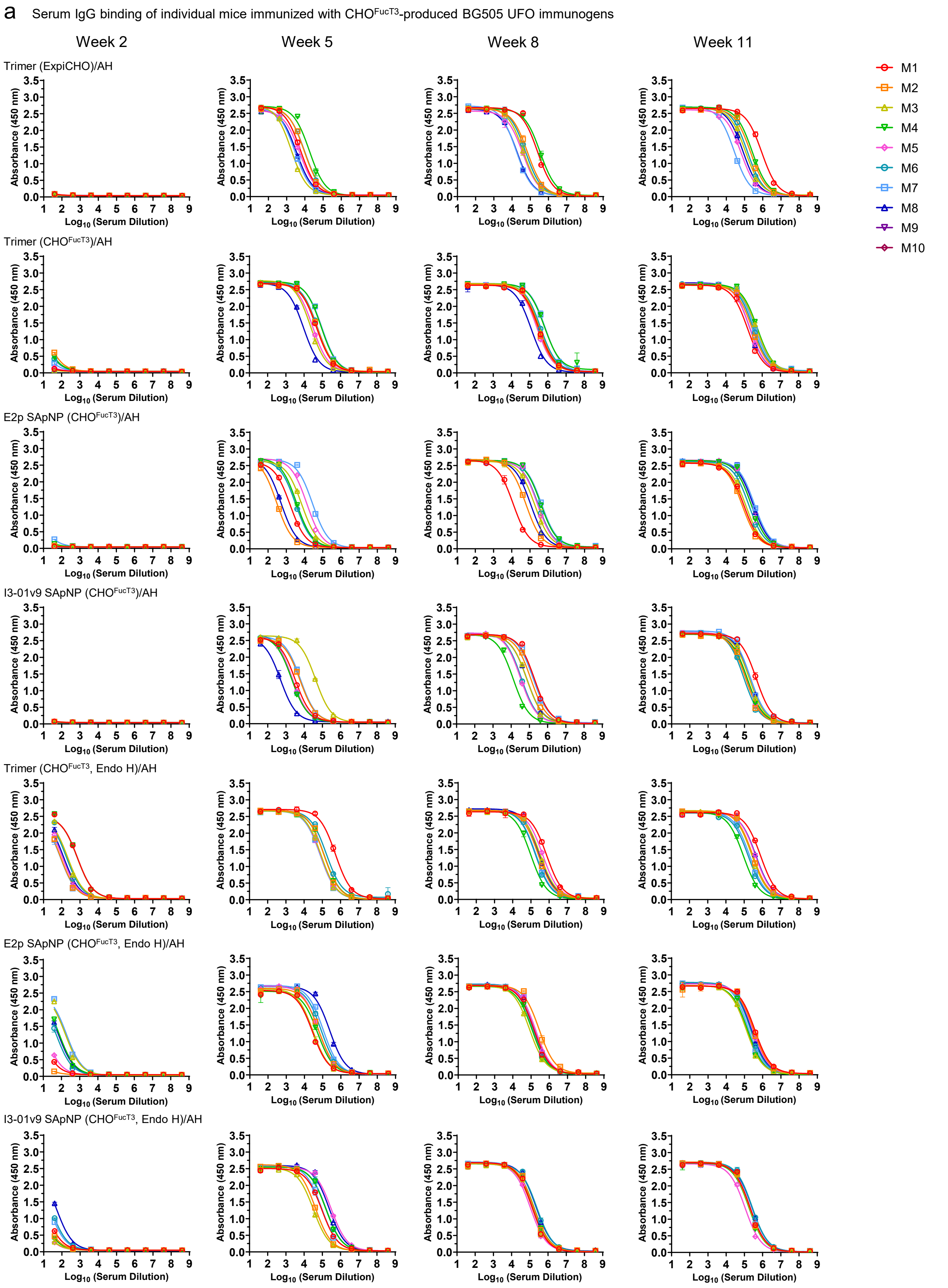

a Serum IgG binding of individual mice immunized with CHO$^{FucT3}$-produced BG505 UFO immunogens
Week 2
Week 5
Week 8
Week 11
Trimer (ExpiCHO)/AH
Trimer (CHO$^{FucT3}$)/AH
E2p SApNP (CHO$^{FucT3}$)/AH
I3-01v9 SApNP (CHO$^{FucT3}$)/AH
Trimer (CHO$^{FucT3}$, Endo H)/AH
E2p SApNP (CHO$^{FucT3}$, Endo H)/AH
I3-01v9 SApNP (CHO$^{FucT3}$, Endo H)/AH
Absorbance (450 nm)
$Log_{10}$ (Serum Dilution)
M1
M2
M3
M4
M5
M6
M7
M8
M9
M10

### Fig. S5

**b** Mouse serum IgG ELISA EC$_{50}$ titers

| Antigen | EC$_{50}$ titers (week 2) | | | | | | | | Geometric Mean |
|---|---|---|---|---|---|---|---|---|---|
| | M1 | M2 | M3 | M4 | M5 | M6 | M7 | M8 | |
| I3-01v9 SApNP (CHO$^{FucT3}$, Endo H) | 12.2 | 8.3 | 4.7 | 6.8 | 4.3 | 25.8 | 21.1 | 51.9 | 11.9 |
| E2p SApNP (CHO$^{FucT3}$, Endo H) | 7.4 | 1.7 | 164.0 | 75.5 | 12.7 | 50.8 | 189.1 | 67.0 | 32.5 |
| Trimer (CHO$^{FucT3}$, Endo H) | 720.7 | 88.3 | 213.0 | 715.4 | 108.9 | 183.5 | 84.7 | 129.6 | 195.4 |
| I3-01v9 SApNP (CHO$^{FucT3}$) | 0.6 | 0.5 | 0.7 | 0.2 | 0.4 | 0.3 | 0.2 | 0.2 | 0.4 |
| E2p SApNP (CHO$^{FucT3}$) | 0.2 | 0.6 | 0.3 | 1.4 | 0.2 | 0.2 | 4.1 | 0.2 | 0.4 |
| Trimer (CHO$^{FucT3}$) | 1.3 | 12.0 | 0.5 | 8.5 | 2.0 | 7.3 | 4.3 | 0.9 | 2.8 |
| Trimer (ExpiCHO) | 0.8 | 0.8 | 0.3 | 0.2 | 0.9 | 0.2 | 0.9 | 0.6 | 0.5 |

| Antigen | EC$_{50}$ titers (week 5) | | | | | | | | Geometric Mean |
|---|---|---|---|---|---|---|---|---|---|
| | M1 | M2 | M3 | M4 | M5 | M6 | M7 | M8 | |
| I3-01v9 SApNP (CHO$^{FucT3}$, Endo H) | 93363 | 41513 | 32204 | 156562 | 307700 | 162315 | 92302 | 236057 | 109890 |
| E2p SApNP (CHO$^{FucT3}$, Endo H) | 27877 | 62533 | 29698 | 52089 | 59079 | 82802 | 107470 | 243143 | 65642 |
| Trimer (CHO$^{FucT3}$, Endo H) | 522930 | 122900 | 85452 | 126776 | 79768 | 166412 | 76646 | 79505 | 124117 |
| I3-01v9 SApNP (CHO$^{FucT3}$) | 3221 | 5930 | 41818 | 2049 | 2261 | 2160 | 6236 | 499.5 | 3544 |
| E2p SApNP (CHO$^{FucT3}$) | 1617 | 295.6 | 7082 | 3682 | 13325 | 3215 | 27631 | 511.5 | 3053 |
| Trimer (CHO$^{FucT3}$) | 48392 | 52272 | 22643 | 88140 | 29480 | 54133 | 94701 | 9309 | 40404 |
| Trimer (ExpiCHO) | 5884 | 9271 | 1805 | 17819 | 5223 | 9677 | 2818 | 3502 | 5530 |

| Antigen | EC$_{50}$ titers (week 8) | | | | | | | | Geometric Mean |
|---|---|---|---|---|---|---|---|---|---|
| | M1 | M2 | M3 | M4 | M5 | M6 | M7 | M8 | |
| I3-01v9 SApNP (CHO$^{FucT3}$, Endo H) | 134178 | 161099 | 134531 | 170045 | 107297 | 244855 | 142443 | 221485 | 159067 |
| E2p SApNP (CHO$^{FucT3}$, Endo H) | 146583 | 278498 | 83469 | 117082 | 177246 | 132292 | 161107 | 168170 | 149791 |
| Trimer (CHO$^{FucT3}$, Endo H) | 732954 | 252006 | 322050 | 97673 | 444054 | 176554 | 185606 | 284690 | 264657 |
| I3-01v9 SApNP (CHO$^{FucT3}$) | 190345 | 121953 | 73240 | 12940 | 32523 | 36100 | 161623 | 73045 | 64644 |
| E2p SApNP (CHO$^{FucT3}$) | 11955 | 58904 | 181442 | 388660 | 260319 | 284595 | 427199 | 109162 | 142661 |
| Trimer (CHO$^{FucT3}$) | 327068 | 293774 | 346319 | 645953 | 249176 | 401196 | 711982 | 120320 | 341291 |
| Trimer (ExpiCHO) | 254748 | 70721 | 42334 | 348836 | 38776 | 55513 | 16933 | 19140 | 60754 |

| Antigen | EC$_{50}$ titers (week 11) | | | | | | | | Geometric Mean |
|---|---|---|---|---|---|---|---|---|---|
| | M1 | M2 | M3 | M4 | M5 | M6 | M7 | M8 | |
| I3-01v9 SApNP (CHO$^{FucT3}$, Endo H) | 203598 | 192413 | 185013 | 178164 | 110133 | 266479 | 206445 | 262767 | 194591 |
| E2p SApNP (CHO$^{FucT3}$, Endo H) | 370717 | 324036 | 126256 | 149263 | 278001 | 205436 | 164167 | 229380 | 216733 |
| Trimer (CHO$^{FucT3}$, Endo H) | 802066 | 294683 | 260152 | 90637 | 414110 | 167301 | 198766 | 441177 | 276190 |
| I3-01v9 SApNP (CHO$^{FucT3}$) | 445556 | 119010 | 183117 | 144677 | 129671 | 91093 | 204096 | 216367 | 171049 |
| E2p SApNP (CHO$^{FucT3}$) | 94601 | 75174 | 118277 | 233771 | 230443 | 173281 | 351376 | 295629 | 173355 |
| Trimer (CHO$^{FucT3}$) | 158082 | 284133 | 496333 | 543398 | 210742 | 319971 | 388271 | 203868 | 299454 |
| Trimer (ExpiCHO) | 882109 | 213457 | 112748 | 262068 | 63993 | 142971 | 27077 | 83654 | 135746 |

Statistical analysis

#### Week 2

| One-way ANOVA with Tukey's multiple comparisons test (w2) | Summary | Adjusted P Value |
|---|---|---|
| Trimer (ExpiCHO) vs. Trimer (CHO$^{FucT3}$) | ns | >0.9999 |
| Trimer (ExpiCHO) vs. E2p SApNP (CHO$^{FucT3}$) | ns | >0.9999 |
| Trimer (ExpiCHO) vs. I3-01v9 SApNP (CHO$^{FucT3}$) | ns | >0.9999 |
| Trimer (ExpiCHO) vs. Trimer (CHO$^{FucT3}$, Endo H) | **** | <0.0001 |
| Trimer (ExpiCHO) vs. E2p SApNP (CHO$^{FucT3}$, Endo H) | ns | 0.8413 |
| Trimer (ExpiCHO) vs. I3-01v9 SApNP (CHO$^{FucT3}$, Endo H) | ns | >0.9999 |
| Trimer (CHO$^{FucT3}$) vs. E2p SApNP (CHO$^{FucT3}$) | ns | >0.9999 |
| Trimer (CHO$^{FucT3}$) vs. I3-01v9 SApNP (CHO$^{FucT3}$) | ns | >0.9999 |
| Trimer (CHO$^{FucT3}$) vs. Trimer (CHO$^{FucT3}$, Endo H) | **** | <0.0001 |
| Trimer (CHO$^{FucT3}$) vs. E2p SApNP (CHO$^{FucT3}$, Endo H) | ns | 0.8745 |
| Trimer (CHO$^{FucT3}$) vs. I3-01v9 SApNP (CHO$^{FucT3}$, Endo H) | ns | >0.9999 |
| E2p SApNP (CHO$^{FucT3}$) vs. I3-01v9 SApNP (CHO$^{FucT3}$) | ns | >0.9999 |
| E2p SApNP (CHO$^{FucT3}$) vs. Trimer (CHO$^{FucT3}$, Endo H) | **** | <0.0001 |
| E2p SApNP (CHO$^{FucT3}$) vs. E2p SApNP (CHO$^{FucT3}$, Endo H) | ns | 0.844 |
| E2p SApNP (CHO$^{FucT3}$) vs. I3-01v9 SApNP (CHO$^{FucT3}$, Endo H) | ns | >0.9999 |
| I3-01v9 SApNP (CHO$^{FucT3}$) vs. Trimer (CHO$^{FucT3}$, Endo H) | **** | <0.0001 |
| I3-01v9 SApNP (CHO$^{FucT3}$) vs. E2p SApNP (CHO$^{FucT3}$, Endo H) | ns | 0.8395 |
| I3-01v9 SApNP (CHO$^{FucT3}$) vs. I3-01v9 SApNP (CHO$^{FucT3}$, Endo H) | ns | >0.9999 |
| Trimer (CHO$^{FucT3}$, Endo H) vs. E2p SApNP (CHO$^{FucT3}$, Endo H) | ** | 0.0049 |
| Trimer (CHO$^{FucT3}$, Endo H) vs. I3-01v9 SApNP (CHO$^{FucT3}$, Endo H) | *** | 0.0002 |
| E2p SApNP (CHO$^{FucT3}$, Endo H) vs. I3-01v9 SApNP (CHO$^{FucT3}$, Endo H) | ns | 0.949 |

#### Week 5

| One-way ANOVA with Tukey's multiple comparisons test (w5) | Summary | Adjusted P Value |
|---|---|---|
| Trimer (ExpiCHO) vs. Trimer (CHO$^{FucT3}$) | ns | 0.904 |
| Trimer (ExpiCHO) vs. E2p SApNP (CHO$^{FucT3}$) | ns | >0.9999 |
| Trimer (ExpiCHO) vs. I3-01v9 SApNP (CHO$^{FucT3}$) | ns | >0.9999 |
| Trimer (ExpiCHO) vs. Trimer (CHO$^{FucT3}$, Endo H) | ** | 0.0029 |
| Trimer (ExpiCHO) vs. E2p SApNP (CHO$^{FucT3}$, Endo H) | ns | 0.3888 |
| Trimer (ExpiCHO) vs. I3-01v9 SApNP (CHO$^{FucT3}$, Endo H) | * | 0.0117 |
| Trimer (CHO$^{FucT3}$) vs. E2p SApNP (CHO$^{FucT3}$) | ns | 0.9057 |
| Trimer (CHO$^{FucT3}$) vs. I3-01v9 SApNP (CHO$^{FucT3}$) | ns | 0.9136 |
| Trimer (CHO$^{FucT3}$) vs. Trimer (CHO$^{FucT3}$, Endo H) | ns | 0.0723 |
| Trimer (CHO$^{FucT3}$) vs. E2p SApNP (CHO$^{FucT3}$, Endo H) | ns | 0.9707 |
| Trimer (CHO$^{FucT3}$) vs. I3-01v9 SApNP (CHO$^{FucT3}$, Endo H) | ns | 0.1994 |
| E2p SApNP (CHO$^{FucT3}$) vs. I3-01v9 SApNP (CHO$^{FucT3}$) | ns | >0.9999 |
| E2p SApNP (CHO$^{FucT3}$) vs. Trimer (CHO$^{FucT3}$, Endo H) | ** | 0.0029 |
| E2p SApNP (CHO$^{FucT3}$) vs. E2p SApNP (CHO$^{FucT3}$, Endo H) | ns | 0.3915 |
| E2p SApNP (CHO$^{FucT3}$) vs. I3-01v9 SApNP (CHO$^{FucT3}$, Endo H) | * | 0.0118 |
| I3-01v9 SApNP (CHO$^{FucT3}$) vs. Trimer (CHO$^{FucT3}$, Endo H) | ** | 0.0032 |
| I3-01v9 SApNP (CHO$^{FucT3}$) vs. E2p SApNP (CHO$^{FucT3}$, Endo H) | ns | 0.4052 |
| I3-01v9 SApNP (CHO$^{FucT3}$) vs. I3-01v9 SApNP (CHO$^{FucT3}$, Endo H) | * | 0.0126 |
| Trimer (CHO$^{FucT3}$, Endo H) vs. E2p SApNP (CHO$^{FucT3}$, Endo H) | ns | 0.4149 |
| Trimer (CHO$^{FucT3}$, Endo H) vs. I3-01v9 SApNP (CHO$^{FucT3}$, Endo H) | ns | 0.9991 |
| E2p SApNP (CHO$^{FucT3}$, Endo H) vs. I3-01v9 SApNP (CHO$^{FucT3}$, Endo H) | ns | 0.7126 |

#### Week 8

| One-way ANOVA with Tukey's multiple comparisons test (w8) | Summary | Adjusted P Value |
|---|---|---|
| Trimer (ExpiCHO) vs. Trimer (CHO$^{FucT3}$) | ** | 0.0023 |
| Trimer (ExpiCHO) vs. E2p SApNP (CHO$^{FucT3}$) | ns | 0.6695 |
| Trimer (ExpiCHO) vs. I3-01v9 SApNP (CHO$^{FucT3}$) | ns | >0.9999 |
| Trimer (ExpiCHO) vs. Trimer (CHO$^{FucT3}$, Endo H) | ns | 0.0526 |
| Trimer (ExpiCHO) vs. E2p SApNP (CHO$^{FucT3}$, Endo H) | ns | 0.9865 |
| Trimer (ExpiCHO) vs. I3-01v9 SApNP (CHO$^{FucT3}$, Endo H) | ns | 0.9756 |
| Trimer (CHO$^{FucT3}$) vs. E2p SApNP (CHO$^{FucT3}$) | ns | 0.1663 |
| Trimer (CHO$^{FucT3}$) vs. I3-01v9 SApNP (CHO$^{FucT3}$) | *** | 0.001 |
| Trimer (CHO$^{FucT3}$) vs. Trimer (CHO$^{FucT3}$, Endo H) | ns | 0.9215 |
| Trimer (CHO$^{FucT3}$) vs. E2p SApNP (CHO$^{FucT3}$, Endo H) | * | 0.0218 |
| Trimer (CHO$^{FucT3}$) vs. I3-01v9 SApNP (CHO$^{FucT3}$, Endo H) | * | 0.0282 |
| E2p SApNP (CHO$^{FucT3}$) vs. I3-01v9 SApNP (CHO$^{FucT3}$) | ns | 0.4962 |
| E2p SApNP (CHO$^{FucT3}$) vs. Trimer (CHO$^{FucT3}$, Endo H) | ns | 0.7816 |
| E2p SApNP (CHO$^{FucT3}$) vs. E2p SApNP (CHO$^{FucT3}$, Endo H) | ns | 0.9783 |
| E2p SApNP (CHO$^{FucT3}$) vs. I3-01v9 SApNP (CHO$^{FucT3}$, Endo H) | ns | 0.9883 |
| I3-01v9 SApNP (CHO$^{FucT3}$) vs. Trimer (CHO$^{FucT3}$, Endo H) | * | 0.0263 |
| I3-01v9 SApNP (CHO$^{FucT3}$) vs. E2p SApNP (CHO$^{FucT3}$, Endo H) | ns | 0.9413 |
| I3-01v9 SApNP (CHO$^{FucT3}$) vs. I3-01v9 SApNP (CHO$^{FucT3}$, Endo H) | ns | 0.9131 |
| Trimer (CHO$^{FucT3}$, Endo H) vs. E2p SApNP (CHO$^{FucT3}$, Endo H) | ns | 0.2739 |
| Trimer (CHO$^{FucT3}$, Endo H) vs. I3-01v9 SApNP (CHO$^{FucT3}$, Endo H) | ns | 0.322 |
| E2p SApNP (CHO$^{FucT3}$, Endo H) vs. I3-01v9 SApNP (CHO$^{FucT3}$, Endo H) | ns | >0.9999 |

#### Week 11

| One-way ANOVA with Tukey's multiple comparisons test (w11) | Summary | Adjusted P Value |
|---|---|---|
| Trimer (ExpiCHO) vs. Trimer (CHO$^{FucT3}$) | ns | 0.8585 |
| Trimer (ExpiCHO) vs. E2p SApNP (CHO$^{FucT3}$) | ns | 0.9999 |
| Trimer (ExpiCHO) vs. I3-01v9 SApNP (CHO$^{FucT3}$) | ns | 0.9997 |
| Trimer (ExpiCHO) vs. Trimer (CHO$^{FucT3}$, Endo H) | ns | 0.8104 |
| Trimer (ExpiCHO) vs. E2p SApNP (CHO$^{FucT3}$, Endo H) | ns | >0.9999 |
| Trimer (ExpiCHO) vs. I3-01v9 SApNP (CHO$^{FucT3}$, Endo H) | ns | >0.9999 |
| Trimer (CHO$^{FucT3}$) vs. E2p SApNP (CHO$^{FucT3}$) | ns | 0.6733 |
| Trimer (CHO$^{FucT3}$) vs. I3-01v9 SApNP (CHO$^{FucT3}$) | ns | 0.6345 |
| Trimer (CHO$^{FucT3}$) vs. Trimer (CHO$^{FucT3}$, Endo H) | ns | >0.9999 |
| Trimer (CHO$^{FucT3}$) vs. E2p SApNP (CHO$^{FucT3}$, Endo H) | ns | 0.8963 |
| Trimer (CHO$^{FucT3}$) vs. I3-01v9 SApNP (CHO$^{FucT3}$, Endo H) | ns | 0.7048 |
| E2p SApNP (CHO$^{FucT3}$) vs. I3-01v9 SApNP (CHO$^{FucT3}$) | ns | >0.9999 |
| E2p SApNP (CHO$^{FucT3}$) vs. Trimer (CHO$^{FucT3}$, Endo H) | ns | 0.6092 |
| E2p SApNP (CHO$^{FucT3}$) vs. E2p SApNP (CHO$^{FucT3}$, Endo H) | ns | 0.9995 |
| E2p SApNP (CHO$^{FucT3}$) vs. I3-01v9 SApNP (CHO$^{FucT3}$, Endo H) | ns | >0.9999 |
| I3-01v9 SApNP (CHO$^{FucT3}$) vs. Trimer (CHO$^{FucT3}$, Endo H) | ns | 0.5696 |
| I3-01v9 SApNP (CHO$^{FucT3}$) vs. E2p SApNP (CHO$^{FucT3}$, Endo H) | ns | 0.9989 |
| I3-01v9 SApNP (CHO$^{FucT3}$) vs. I3-01v9 SApNP (CHO$^{FucT3}$, Endo H) | ns | >0.9999 |
| Trimer (CHO$^{FucT3}$, Endo H) vs. E2p SApNP (CHO$^{FucT3}$, Endo H) | ns | 0.8551 |
| Trimer (CHO$^{FucT3}$, Endo H) vs. I3-01v9 SApNP (CHO$^{FucT3}$, Endo H) | ns | 0.6419 |
| E2p SApNP (CHO$^{FucT3}$, Endo H) vs. I3-01v9 SApNP (CHO$^{FucT3}$, Endo H) | ns | 0.9997 |

**C** Mouse serum IgE ELISA OD$_{450}$ values

| Antigen | OD$_{450}$ values (week 2) | | | | | | | | Geometric Mean |
|---|---|---|---|---|---|---|---|---|---|
| | M1 | M2 | M3 | M4 | M5 | M6 | M7 | M8 | |
| I3-01v9 SApNP (CHO$^{FucT3}$, Endo H) | 0.065 | 0.064 | 0.072 | 0.065 | 0.057 | 0.072 | 0.066 | 0.073 | 0.067 |
| E2p SApNP (CHO$^{FucT3}$, Endo H) | 0.059 | 0.059 | 0.061 | 0.057 | 0.060 | 0.054 | 0.151 | 0.074 | 0.067 |
| Trimer (CHO$^{FucT3}$, Endo H) | 0.198 | 0.063 | 0.068 | 0.142 | 0.088 | 0.073 | 0.085 | 0.074 | 0.091 |
| I3-01v9 SApNP (CHO$^{FucT3}$) | 0.051 | 0.052 | 0.054 | 0.052 | 0.050 | 0.050 | 0.051 | 0.049 | 0.051 |
| E2p SApNP (CHO$^{FucT3}$) | 0.049 | 0.051 | 0.048 | 0.049 | 0.055 | 0.051 | 0.050 | 0.051 | 0.051 |
| Trimer (CHO$^{FucT3}$) | 0.050 | 0.056 | 0.051 | 0.061 | 0.054 | 0.054 | 0.053 | 0.055 | 0.054 |
| Trimer (ExpiCHO) | 0.050 | 0.055 | 0.052 | 0.048 | 0.051 | 0.050 | 0.054 | 0.050 | 0.051 |

| Antigen | OD$_{450}$ values (week 5) | | | | | | | | Geometric Mean |
|---|---|---|---|---|---|---|---|---|---|
| | M1 | M2 | M3 | M4 | M5 | M6 | M7 | M8 | |
| I3-01v9 SApNP (CHO$^{FucT3}$, Endo H) | 0.119 | 0.262 | 0.207 | 0.145 | 0.162 | 0.102 | 0.174 | 0.140 | 0.157 |
| E2p SApNP (CHO$^{FucT3}$, Endo H) | 0.128 | 0.109 | 0.082 | 0.094 | 0.112 | 0.103 | 0.121 | 0.095 | 0.105 |
| Trimer (CHO$^{FucT3}$, Endo H) | 0.160 | 0.109 | 0.127 | 0.206 | 0.154 | 0.179 | 0.144 | 0.097 | 0.143 |
| I3-01v9 SApNP (CHO$^{FucT3}$) | 0.230 | 0.177 | 0.052 | 0.157 | 0.143 | 0.127 | 0.268 | 0.058 | 0.132 |
| E2p SApNP (CHO$^{FucT3}$) | 0.065 | 0.103 | 0.080 | 0.088 | 0.104 | 0.250 | 0.190 | 0.081 | 0.108 |
| Trimer (CHO$^{FucT3}$) | 0.132 | 0.115 | 0.184 | 0.140 | 0.338 | 0.142 | 0.109 | 0.228 | 0.162 |
| Trimer (ExpiCHO) | 0.083 | 0.071 | 0.062 | 0.074 | 0.065 | 0.109 | 0.059 | 0.059 | 0.071 |

| Antigen | OD$_{450}$ values (week 8) | | | | | | | | Geometric Mean |
|---|---|---|---|---|---|---|---|---|---|
| | M1 | M2 | M3 | M4 | M5 | M6 | M7 | M8 | |
| I3-01v9 SApNP (CHO$^{FucT3}$, Endo H) | 0.116 | 0.250 | 0.124 | 0.125 | 0.177 | 0.183 | 0.121 | 0.118 | 0.146 |
| E2p SApNP (CHO$^{FucT3}$, Endo H) | 0.109 | 0.074 | 0.174 | 0.103 | 0.120 | 0.087 | 0.109 | 0.091 | 0.105 |
| Trimer (CHO$^{FucT3}$, Endo H) | 0.176 | 0.164 | 0.167 | 0.218 | 0.135 | 0.203 | 0.201 | 0.103 | 0.167 |
| I3-01v9 SApNP (CHO$^{FucT3}$) | 0.124 | 0.214 | 0.093 | 0.469 | 0.640 | 0.319 | 0.138 | 0.245 | 0.231 |
| E2p SApNP (CHO$^{FucT3}$) | 0.451 | 0.138 | 0.149 | 0.199 | 0.107 | 0.103 | 0.148 | 0.125 | 0.158 |
| Trimer (CHO$^{FucT3}$) | 0.220 | 0.143 | 0.146 | 0.150 | 0.528 | 0.153 | 0.128 | 0.379 | 0.202 |
| Trimer (ExpiCHO) | 0.119 | 0.199 | 0.084 | 0.105 | 0.081 | 0.256 | 0.076 | 0.119 | 0.119 |

| Antigen | OD$_{450}$ values (week 11) | | | | | | | | Geometric Mean |
|---|---|---|---|---|---|---|---|---|---|
| | M1 | M2 | M3 | M4 | M5 | M6 | M7 | M8 | |
| I3-01v9 SApNP (CHO$^{FucT3}$, Endo H) | 0.233 | 0.330 | 0.256 | 0.247 | 0.232 | 0.284 | 0.202 | 0.258 | 0.253 |
| E2p SApNP (CHO$^{FucT3}$, Endo H) | 0.175 | 0.130 | 0.259 | 0.168 | 0.162 | 0.134 | 0.165 | 0.144 | 0.164 |
| Trimer (CHO$^{FucT3}$, Endo H) | 0.328 | 0.207 | 0.293 | 0.248 | 0.167 | 0.192 | 0.256 | 0.170 | 0.226 |
| I3-01v9 SApNP (CHO$^{FucT3}$) | 0.224 | 0.219 | 0.244 | 0.185 | 0.804 | 0.623 | 0.209 | 0.253 | 0.296 |
| E2p SApNP (CHO$^{FucT3}$) | 0.317 | 0.245 | 0.177 | 0.157 | 0.143 | 0.183 | 0.196 | 0.146 | 0.189 |
| Trimer (CHO$^{FucT3}$) | 0.350 | 0.183 | 0.263 | 0.234 | 1.021 | 0.314 | 0.176 | 0.472 | 0.318 |
| Trimer (ExpiCHO) | 0.142 | 0.216 | 0.078 | 0.216 | 0.115 | 0.240 | 0.085 | 0.170 | 0.146 |

#### Statistical analysis

##### Week 2

| One-way ANOVA with Tukey's multiple comparisons test (w2) | Summary | Adjusted P Value |
|---|---|---|
| Trimer (ExpiCHO) vs. Trimer (CHO$^{FucT3}$) | ns | >0.9999 |
| Trimer (ExpiCHO) vs. E2p SApNP (CHO$^{FucT3}$) | ns | >0.9999 |
| Trimer (ExpiCHO) vs. I3-01v9 SApNP (CHO$^{FucT3}$) | ns | >0.9999 |
| Trimer (ExpiCHO) vs. Trimer (CHO$^{FucT3}$, Endo H) | ** | 0.0012 |
| Trimer (ExpiCHO) vs. E2p SApNP (CHO$^{FucT3}$, Endo H) | ns | 0.4951 |
| Trimer (ExpiCHO) vs. I3-01v9 SApNP (CHO$^{FucT3}$, Endo H) | ns | 0.7875 |
| Trimer (CHO$^{FucT3}$) vs. E2p SApNP (CHO$^{FucT3}$) | ns | 0.9999 |
| Trimer (CHO$^{FucT3}$) vs. I3-01v9 SApNP (CHO$^{FucT3}$) | ns | >0.9999 |
| Trimer (CHO$^{FucT3}$) vs. Trimer (CHO$^{FucT3}$, Endo H) | ** | 0.0028 |
| Trimer (CHO$^{FucT3}$) vs. E2p SApNP (CHO$^{FucT3}$, Endo H) | ns | 0.6724 |
| Trimer (CHO$^{FucT3}$) vs. I3-01v9 SApNP (CHO$^{FucT3}$, Endo H) | ns | 0.9101 |
| E2p SApNP (CHO$^{FucT3}$) vs. I3-01v9 SApNP (CHO$^{FucT3}$) | ns | >0.9999 |
| E2p SApNP (CHO$^{FucT3}$) vs. Trimer (CHO$^{FucT3}$, Endo H) | *** | 0.001 |
| E2p SApNP (CHO$^{FucT3}$) vs. E2p SApNP (CHO$^{FucT3}$, Endo H) | ns | 0.4518 |
| E2p SApNP (CHO$^{FucT3}$) vs. I3-01v9 SApNP (CHO$^{FucT3}$, Endo H) | ns | 0.7488 |
| I3-01v9 SApNP (CHO$^{FucT3}$) vs. Trimer (CHO$^{FucT3}$, Endo H) | ** | 0.0012 |
| I3-01v9 SApNP (CHO$^{FucT3}$) vs. E2p SApNP (CHO$^{FucT3}$, Endo H) | ns | 0.4878 |
| I3-01v9 SApNP (CHO$^{FucT3}$) vs. I3-01v9 SApNP (CHO$^{FucT3}$, Endo H) | ns | 0.7812 |
| Trimer (CHO$^{FucT3}$, Endo H) vs. E2p SApNP (CHO$^{FucT3}$, Endo H) | ns | 0.1903 |
| Trimer (CHO$^{FucT3}$, Endo H) vs. I3-01v9 SApNP (CHO$^{FucT3}$, Endo H) | ns | 0.0682 |
| E2p SApNP (CHO$^{FucT3}$, Endo H) vs. I3-01v9 SApNP (CHO$^{FucT3}$, Endo H) | ns | 0.9991 |

##### Week 5

| One-way ANOVA with Tukey's multiple comparisons test (w5) | Summary | Adjusted P Value |
|---|---|---|
| Trimer (ExpiCHO) vs. Trimer (CHO$^{FucT3}$) | ** | 0.0081 |
| Trimer (ExpiCHO) vs. E2p SApNP (CHO$^{FucT3}$) | ns | 0.5796 |
| Trimer (ExpiCHO) vs. I3-01v9 SApNP (CHO$^{FucT3}$) | ns | 0.0714 |
| Trimer (ExpiCHO) vs. Trimer (CHO$^{FucT3}$, Endo H) | ns | 0.105 |
| Trimer (ExpiCHO) vs. E2p SApNP (CHO$^{FucT3}$, Endo H) | ns | 0.884 |
| Trimer (ExpiCHO) vs. I3-01v9 SApNP (CHO$^{FucT3}$, Endo H) | * | 0.0221 |
| Trimer (CHO$^{FucT3}$) vs. E2p SApNP (CHO$^{FucT3}$) | ns | 0.4374 |
| Trimer (CHO$^{FucT3}$) vs. I3-01v9 SApNP (CHO$^{FucT3}$) | ns | 0.982 |
| Trimer (CHO$^{FucT3}$) vs. Trimer (CHO$^{FucT3}$, Endo H) | ns | 0.9549 |
| Trimer (CHO$^{FucT3}$) vs. E2p SApNP (CHO$^{FucT3}$, Endo H) | ns | 0.172 |
| Trimer (CHO$^{FucT3}$) vs. I3-01v9 SApNP (CHO$^{FucT3}$, Endo H) | ns | 0.9998 |
| E2p SApNP (CHO$^{FucT3}$) vs. I3-01v9 SApNP (CHO$^{FucT3}$) | ns | 0.9032 |
| E2p SApNP (CHO$^{FucT3}$) vs. Trimer (CHO$^{FucT3}$, Endo H) | ns | 0.9518 |
| E2p SApNP (CHO$^{FucT3}$) vs. E2p SApNP (CHO$^{FucT3}$, Endo H) | ns | 0.998 |
| E2p SApNP (CHO$^{FucT3}$) vs. I3-01v9 SApNP (CHO$^{FucT3}$, Endo H) | ns | 0.6663 |
| I3-01v9 SApNP (CHO$^{FucT3}$) vs. Trimer (CHO$^{FucT3}$, Endo H) | ns | >0.9999 |
| I3-01v9 SApNP (CHO$^{FucT3}$) vs. E2p SApNP (CHO$^{FucT3}$, Endo H) | ns | 0.6128 |
| I3-01v9 SApNP (CHO$^{FucT3}$) vs. I3-01v9 SApNP (CHO$^{FucT3}$, Endo H) | ns | 0.9992 |
| Trimer (CHO$^{FucT3}$, Endo H) vs. E2p SApNP (CHO$^{FucT3}$, Endo H) | ns | 0.718 |
| Trimer (CHO$^{FucT3}$, Endo H) vs. I3-01v9 SApNP (CHO$^{FucT3}$, Endo H) | ns | 0.9956 |
| E2p SApNP (CHO$^{FucT3}$, Endo H) vs. I3-01v9 SApNP (CHO$^{FucT3}$, Endo H) | ns | 0.3301 |

##### Week 8

| One-way ANOVA with Tukey's multiple comparisons test (w8) | Summary | Adjusted P Value |
|---|---|---|
| Trimer (ExpiCHO) vs. Trimer (CHO$^{FucT3}$) | ns | 0.4899 |
| Trimer (ExpiCHO) vs. E2p SApNP (CHO$^{FucT3}$) | ns | 0.9715 |
| Trimer (ExpiCHO) vs. I3-01v9 SApNP (CHO$^{FucT3}$) | ns | 0.0899 |
| Trimer (ExpiCHO) vs. Trimer (CHO$^{FucT3}$, Endo H) | ns | 0.9867 |
| Trimer (ExpiCHO) vs. E2p SApNP (CHO$^{FucT3}$, Endo H) | ns | 0.9996 |
| Trimer (ExpiCHO) vs. I3-01v9 SApNP (CHO$^{FucT3}$, Endo H) | ns | 0.9996 |
| Trimer (CHO$^{FucT3}$) vs. E2p SApNP (CHO$^{FucT3}$) | ns | 0.9506 |
| Trimer (CHO$^{FucT3}$) vs. I3-01v9 SApNP (CHO$^{FucT3}$) | ns | 0.966 |
| Trimer (CHO$^{FucT3}$) vs. Trimer (CHO$^{FucT3}$, Endo H) | ns | 0.916 |
| Trimer (CHO$^{FucT3}$) vs. E2p SApNP (CHO$^{FucT3}$, Endo H) | ns | 0.2626 |
| Trimer (CHO$^{FucT3}$) vs. I3-01v9 SApNP (CHO$^{FucT3}$, Endo H) | ns | 0.75 |
| E2p SApNP (CHO$^{FucT3}$) vs. I3-01v9 SApNP (CHO$^{FucT3}$) | ns | 0.4691 |
| E2p SApNP (CHO$^{FucT3}$) vs. Trimer (CHO$^{FucT3}$, Endo H) | ns | >0.9999 |
| E2p SApNP (CHO$^{FucT3}$) vs. E2p SApNP (CHO$^{FucT3}$, Endo H) | ns | 0.8484 |
| E2p SApNP (CHO$^{FucT3}$) vs. I3-01v9 SApNP (CHO$^{FucT3}$, Endo H) | ns | 0.9989 |
| I3-01v9 SApNP (CHO$^{FucT3}$) vs. Trimer (CHO$^{FucT3}$, Endo H) | ns | 0.3933 |
| I3-01v9 SApNP (CHO$^{FucT3}$) vs. E2p SApNP (CHO$^{FucT3}$, Endo H) | * | 0.0333 |
| I3-01v9 SApNP (CHO$^{FucT3}$) vs. I3-01v9 SApNP (CHO$^{FucT3}$, Endo H) | ns | 0.2134 |
| Trimer (CHO$^{FucT3}$, Endo H) vs. E2p SApNP (CHO$^{FucT3}$, Endo H) | ns | 0.8998 |
| Trimer (CHO$^{FucT3}$, Endo H) vs. I3-01v9 SApNP (CHO$^{FucT3}$, Endo H) | ns | 0.9998 |
| E2p SApNP (CHO$^{FucT3}$, Endo H) vs. I3-01v9 SApNP (CHO$^{FucT3}$, Endo H) | ns | 0.9822 |

##### Week 11

| One-way ANOVA with Tukey's multiple comparisons test (w11) | Summary | Adjusted P Value |
|---|---|---|
| Trimer (ExpiCHO) vs. Trimer (CHO$^{FucT3}$) | ns | 0.055 |
| Trimer (ExpiCHO) vs. E2p SApNP (CHO$^{FucT3}$) | ns | 0.9984 |
| Trimer (ExpiCHO) vs. I3-01v9 SApNP (CHO$^{FucT3}$) | ns | 0.1488 |
| Trimer (ExpiCHO) vs. Trimer (CHO$^{FucT3}$, Endo H) | ns | 0.9423 |
| Trimer (ExpiCHO) vs. E2p SApNP (CHO$^{FucT3}$, Endo H) | ns | >0.9999 |
| Trimer (ExpiCHO) vs. I3-01v9 SApNP (CHO$^{FucT3}$, Endo H) | ns | 0.8237 |
| Trimer (CHO$^{FucT3}$) vs. E2p SApNP (CHO$^{FucT3}$) | ns | 0.1775 |
| Trimer (CHO$^{FucT3}$) vs. I3-01v9 SApNP (CHO$^{FucT3}$) | ns | 0.9994 |
| Trimer (CHO$^{FucT3}$) vs. Trimer (CHO$^{FucT3}$, Endo H) | ns | 0.4295 |
| Trimer (CHO$^{FucT3}$) vs. E2p SApNP (CHO$^{FucT3}$, Endo H) | ns | 0.0752 |
| Trimer (CHO$^{FucT3}$) vs. I3-01v9 SApNP (CHO$^{FucT3}$, Endo H) | ns | 0.63 |
| E2p SApNP (CHO$^{FucT3}$) vs. I3-01v9 SApNP (CHO$^{FucT3}$) | ns | 0.3831 |
| E2p SApNP (CHO$^{FucT3}$) vs. Trimer (CHO$^{FucT3}$, Endo H) | ns | 0.9985 |
| E2p SApNP (CHO$^{FucT3}$) vs. E2p SApNP (CHO$^{FucT3}$, Endo H) | ns | 0.9997 |
| E2p SApNP (CHO$^{FucT3}$) vs. I3-01v9 SApNP (CHO$^{FucT3}$, Endo H) | ns | 0.9807 |
| I3-01v9 SApNP (CHO$^{FucT3}$) vs. Trimer (CHO$^{FucT3}$, Endo H) | ns | 0.7074 |
| I3-01v9 SApNP (CHO$^{FucT3}$) vs. E2p SApNP (CHO$^{FucT3}$, Endo H) | ns | 0.1934 |
| I3-01v9 SApNP (CHO$^{FucT3}$) vs. I3-01v9 SApNP (CHO$^{FucT3}$, Endo H) | ns | 0.8722 |
| Trimer (CHO$^{FucT3}$, Endo H) vs. E2p SApNP (CHO$^{FucT3}$, Endo H) | ns | 0.9695 |
| Trimer (CHO$^{FucT3}$, Endo H) vs. I3-01v9 SApNP (CHO$^{FucT3}$, Endo H) | ns | >0.9999 |
| E2p SApNP (CHO$^{FucT3}$, Endo H) vs. I3-01v9 SApNP (CHO$^{FucT3}$, Endo H) | ns | 0.8821 |

### Fig. S5

d Week 11 serum neutralization in mice immunized with CHO$^{FucT3}$-produced BG505 UFO immunogens

**Tier 2 Clade A BG505.T332N**

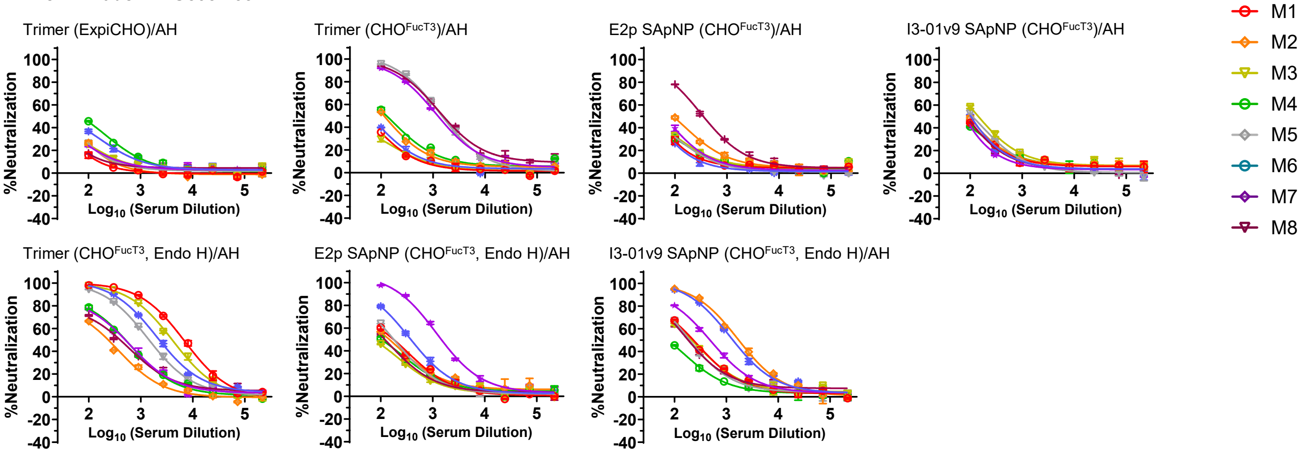


**MLV (negative control)**

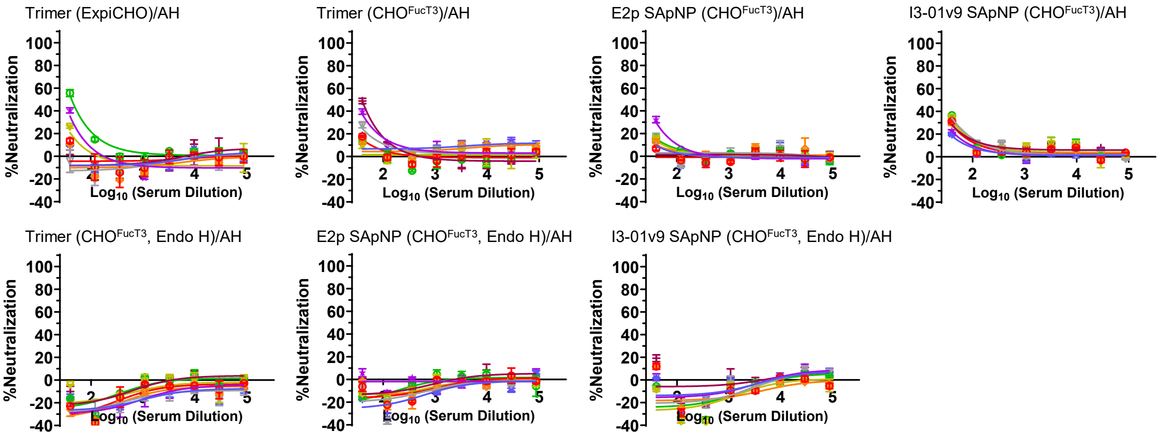


Mouse serum neutralizing $ID_{50}$ titers

| W11 BG505 T332N | M1 | M2 | M3 | M4 | M5 | M6 | M7 | M8 | RR | Geomean |
|---|---|---|---|---|---|---|---|---|---|---|
| I3-01v9 SApNP (CHO$^{FucT3}$, Endo H) | 236.3 | 1762 | 224.4 | 50 | 207.1 | 1414 | 492.1 | 204.6 | 88% | 329.1 |
| E2p SApNP (CHO$^{FucT3}$, Endo H) | 182.2 | 166.8 | 50 | 132.2 | 183.9 | 390.7 | 1583 | 131.1 | 88% | 204.0 |
| Trimer (CHO$^{FucT3}$, Endo H) | 6573 | 228.8 | 3975 | 485.3 | 1477 | 2403 | 486.8 | 388 | 100% | 1086.7 |
| I3-01v9 SApNP (CHO$^{FucT3}$) | 50 | 50 | 159.6 | 50 | 117.4 | 103.4 | 50 | 50 | 38% | 70.4 |
| E2p SApNP (CHO$^{FucT3}$) | 50 | 50 | 50 | 50 | 50 | 50 | 50 | 372.4 | 13% | 64.3 |
| Trimer (CHO$^{FucT3}$) | 50 | 124.8 | 50 | 145.5 | 1540 | 50 | 1301 | 1719 | 63% | 229.9 |
| Trimer (ExpiCHO) | 50 | 50 | 50 | 50 | 50 | 50 | 50 | 50 | 0% | 50 |

Statistical analysis

| One-way ANOVA with Tukey's multiple comparisons test (w11) | Summary | Adjusted P Value |
|---|---|---|
| Trimer (ExpiCHO) vs. Trimer (CHO$^{FucT3}$) | ns | 0.8893 |
| Trimer (ExpiCHO) vs. E2p SApNP (CHO$^{FucT3}$) | ns | >0.9999 |
| Trimer (ExpiCHO) vs. I3-01v9 SApNP (CHO$^{FucT3}$) | ns | >0.9999 |
| Trimer (ExpiCHO) vs. Trimer (CHO$^{FucT3}$, Endo H) | ** | 0.0027 |
| Trimer (ExpiCHO) vs. E2p SApNP (CHO$^{FucT3}$, Endo H) | ns | 0.9952 |
| Trimer (ExpiCHO) vs. I3-01v9 SApNP (CHO$^{FucT3}$, Endo H) | ns | 0.9247 |
| Trimer (CHO$^{FucT3}$) vs. E2p SApNP (CHO$^{FucT3}$) | ns | 0.9192 |
| Trimer (CHO$^{FucT3}$) vs. I3-01v9 SApNP (CHO$^{FucT3}$) | ns | 0.9113 |
| Trimer (CHO$^{FucT3}$) vs. Trimer (CHO$^{FucT3}$, Endo H) | ns | 0.0761 |
| Trimer (CHO$^{FucT3}$) vs. E2p SApNP (CHO$^{FucT3}$, Endo H) | ns | 0.9974 |
| Trimer (CHO$^{FucT3}$) vs. I3-01v9 SApNP (CHO$^{FucT3}$, Endo H) | ns | >0.9999 |
| E2p SApNP (CHO$^{FucT3}$) vs. I3-01v9 SApNP (CHO$^{FucT3}$) | ns | >0.9999 |
| E2p SApNP (CHO$^{FucT3}$) vs. Trimer (CHO$^{FucT3}$, Endo H) | ** | 0.0036 |
| E2p SApNP (CHO$^{FucT3}$) vs. E2p SApNP (CHO$^{FucT3}$, Endo H) | ns | 0.9978 |
| E2p SApNP (CHO$^{FucT3}$) vs. I3-01v9 SApNP (CHO$^{FucT3}$, Endo H) | ns | 0.9477 |
| I3-01v9 SApNP (CHO$^{FucT3}$) vs. Trimer (CHO$^{FucT3}$, Endo H) | ** | 0.0033 |
| I3-01v9 SApNP (CHO$^{FucT3}$) vs. E2p SApNP (CHO$^{FucT3}$, Endo H) | ns | 0.9972 |
| I3-01v9 SApNP (CHO$^{FucT3}$) vs. I3-01v9 SApNP (CHO$^{FucT3}$, Endo H) | ns | 0.9417 |
| Trimer (CHO$^{FucT3}$, Endo H) vs. E2p SApNP (CHO$^{FucT3}$, Endo H) | * | 0.0177 |
| Trimer (CHO$^{FucT3}$, Endo H) vs. I3-01v9 SApNP (CHO$^{FucT3}$, Endo H) | ns | 0.0596 |
| E2p SApNP (CHO$^{FucT3}$, Endo H) vs. I3-01v9 SApNP (CHO$^{FucT3}$, Endo H) | ns | 0.9992 |

**Fig. S5**

e Purified week 11 IgG neutralization of individual mice immunized with CHO$^{FucT3}$-produced BG505 UFO immunogens

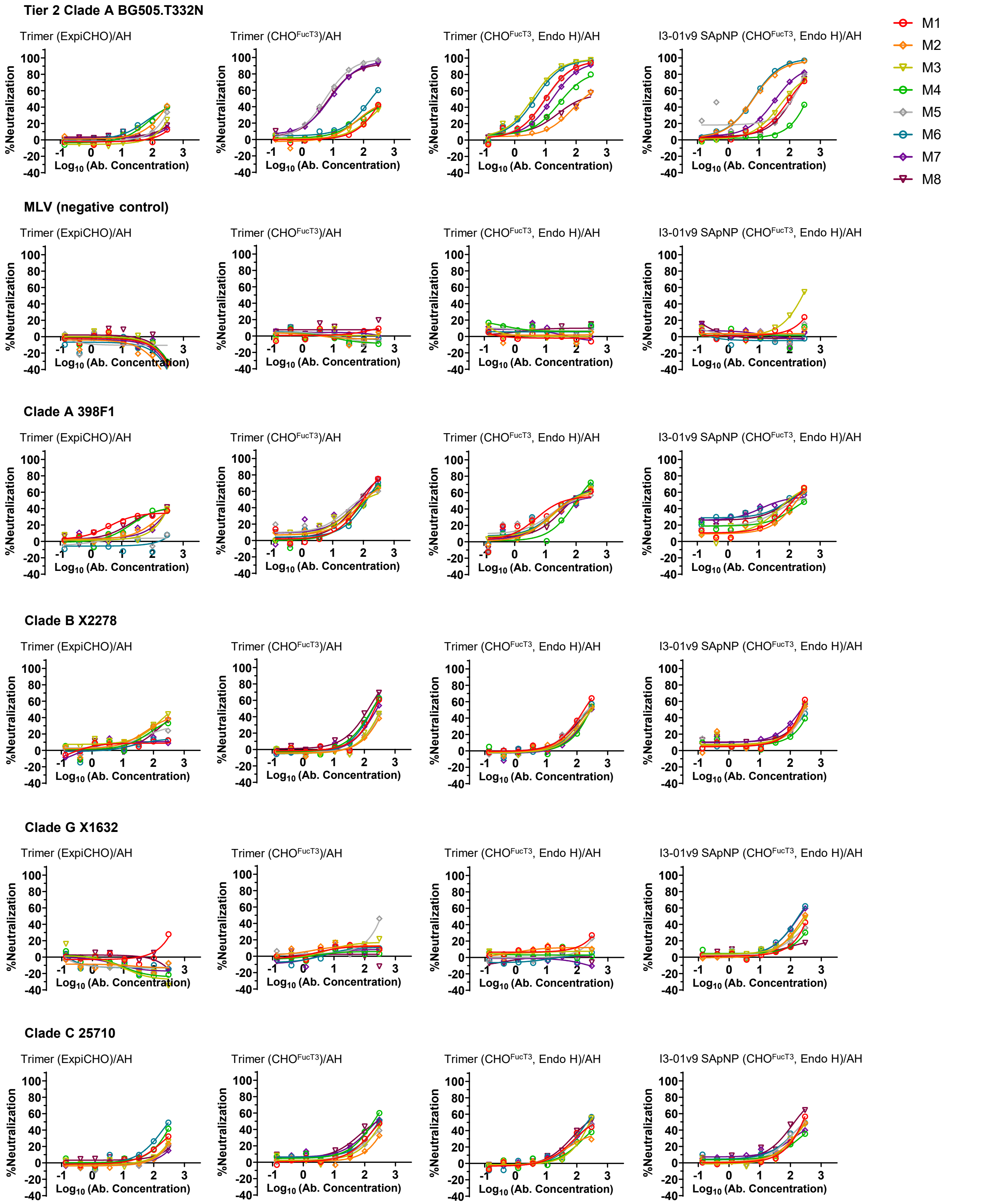

### Fig. S5

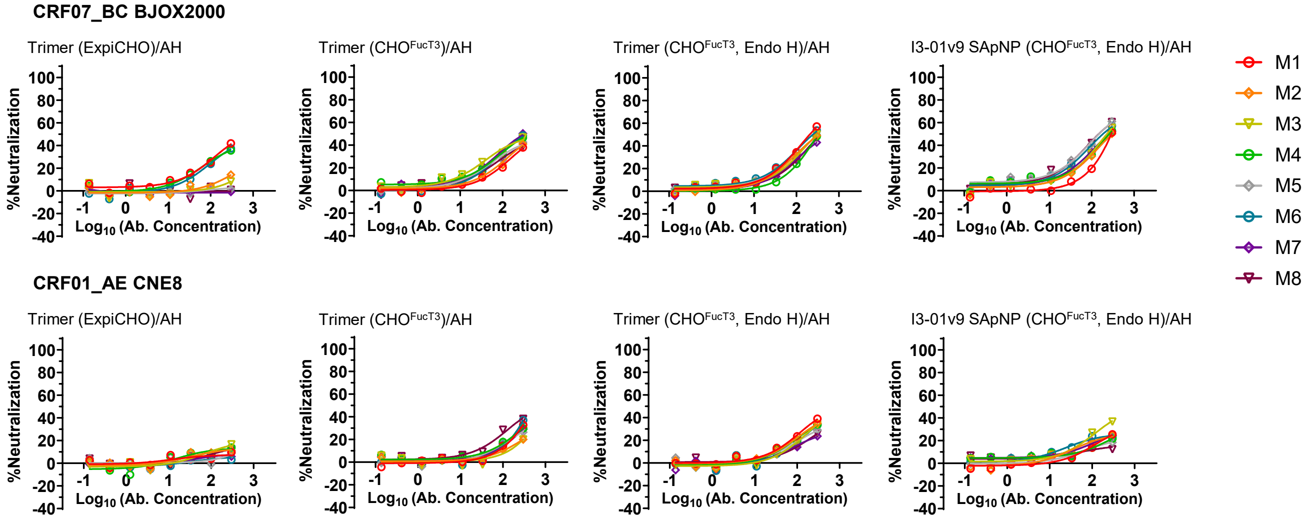


f

Neutralization $IC_{50}$ values of purified IgG

| W11 BG505 T332N | M1 | M2 | M3 | M4 | M5 | M6 | M7 | M8 | RR | Geomean |
|---|---|---|---|---|---|---|---|---|---|---|
| I3-01v9 SApNP ($CHO^{FucT3}$, Endo H) | 99.44 | 6.717 | 75.34 | 600 | 94.1 | 6.253 | 36.26 | 113.2 | 88% | 54.1 |
| Trimer ($CHO^{FucT3}$, Endo H) | 9.942 | 152 | 3.644 | 38.78 | 10.23 | 4.459 | 16.42 | 122.4 | 100% | 19.3 |
| Trimer ($CHO^{FucT3}$) | 600 | 600 | 600 | 600 | 5.822 | 168.6 | 8.049 | 7.964 | 50% | 97.5 |
| Trimer (ExpiCHO) | 600 | 600 | 600 | 600 | 600 | 600 | 600 | 600 | 0% | 600.0 |

| W11 MLV | M1 | M2 | M3 | M4 | M5 | M6 | M7 | M8 | RR | Geomean |
|---|---|---|---|---|---|---|---|---|---|---|
| I3-01v9 SApNP ($CHO^{FucT3}$, Endo H) | 600 | 600 | 269.7 | 600 | 600 | 600 | 600 | 600 | 13% | 542.9 |
| Trimer ($CHO^{FucT3}$, Endo H) | 600 | 600 | 600 | 600 | 600 | 600 | 600 | 600 | 0% | 600.0 |
| Trimer ($CHO^{FucT3}$) | 600 | 600 | 600 | 600 | 600 | 600 | 600 | 600 | 0% | 600.0 |
| Trimer (ExpiCHO) | 600 | 600 | 600 | 600 | 600 | 600 | 600 | 600 | 0% | 600.0 |

| W11 398F1 | M1 | M2 | M3 | M4 | M5 | M6 | M7 | M8 | RR | Geomean |
|---|---|---|---|---|---|---|---|---|---|---|
| I3-01v9 SApNP ($CHO^{FucT3}$, Endo H) | 118.2 | 154.9 | 88.99 | 182.4 | 111 | 63.57 | 51.16 | 73.63 | 100% | 97.1 |
| Trimer ($CHO^{FucT3}$, Endo H) | 57.99 | 79.25 | 72.97 | 104.1 | 78.15 | 68.31 | 96.57 | 78.76 | 100% | 78.3 |
| Trimer ($CHO^{FucT3}$) | 96.6 | 110.4 | 112.4 | 124.7 | 78.25 | 124.5 | 88.76 | 77.11 | 100% | 100.0 |
| Trimer (ExpiCHO) | 600 | 600 | 600 | 600 | 600 | 600 | 600 | 600 | 0% | 600.0 |

| W11 X2278 | M1 | M2 | M3 | M4 | M5 | M6 | M7 | M8 | RR | Geomean |
|---|---|---|---|---|---|---|---|---|---|---|
| I3-01v9 SApNP ($CHO^{FucT3}$, Endo H) | 221.4 | 270 | 246.8 | 600 | 297.9 | 600 | 194.8 | 246.5 | 75% | 305.6 |
| Trimer ($CHO^{FucT3}$, Endo H) | 171.2 | 246.3 | 292.2 | 297.7 | 208.7 | 232.9 | 600 | 238.2 | 88% | 266.5 |
| Trimer ($CHO^{FucT3}$) | 236.3 | 600 | 600 | 194.2 | 600 | 208.2 | 281.7 | 136.8 | 63% | 307.3 |
| Trimer (ExpiCHO) | 600 | 600 | 600 | 600 | 600 | 600 | 600 | 600 | 0% | 600.0 |

| W11 X1632 | M1 | M2 | M3 | M4 | M5 | M6 | M7 | M8 | RR | Geomean |
|---|---|---|---|---|---|---|---|---|---|---|
| I3-01v9 SApNP ($CHO^{FucT3}$, Endo H) | 600 | 282.3 | 600 | 600 | 600 | 179 | 190.3 | 600 | 38% | 406.6 |
| Trimer ($CHO^{FucT3}$, Endo H) | 600 | 600 | 600 | 600 | 600 | 600 | 600 | 600 | 0% | 600.0 |
| Trimer ($CHO^{FucT3}$) | 600 | 600 | 600 | 600 | 600 | 600 | 600 | 600 | 0% | 600.0 |
| Trimer (ExpiCHO) | 600 | 600 | 600 | 600 | 600 | 600 | 600 | 600 | 0% | 600.0 |

| W11 25710 | M1 | M2 | M3 | M4 | M5 | M6 | M7 | M8 | RR | Geomean |
|---|---|---|---|---|---|---|---|---|---|---|
| I3-01v9 SApNP ($CHO^{FucT3}$, Endo H) | 268.2 | 292.7 | 600 | 600 | 246 | 237.9 | 600 | 132.6 | 63% | 327.3 |
| Trimer ($CHO^{FucT3}$, Endo H) | 268.9 | 600 | 285.9 | 600 | 265.3 | 217.7 | 268.8 | 200.2 | 75% | 310.3 |
| Trimer ($CHO^{FucT3}$) | 297.2 | 600 | 600 | 186.3 | 600 | 267.7 | 204.3 | 184.2 | 63% | 323.7 |
| Trimer (ExpiCHO) | 600 | 600 | 600 | 600 | 600 | 600 | 600 | 600 | 0% | 600.0 |

| W11 BJOX2000 | M1 | M2 | M3 | M4 | M5 | M6 | M7 | M8 | RR | Geomean |
|---|---|---|---|---|---|---|---|---|---|---|
| I3-01v9 SApNP ($CHO^{FucT3}$, Endo H) | 600 | 249.4 | 211.6 | 224.4 | 135.8 | 165.8 | 211.3 | 137.3 | 88% | 215.4 |
| Trimer ($CHO^{FucT3}$, Endo H) | 191.3 | 249.7 | 252.5 | 600 | 245.1 | 201.3 | 600 | 295.5 | 75% | 298.7 |
| Trimer ($CHO^{FucT3}$) | 600 | 600 | 226 | 244 | 600 | 251.4 | 222.1 | 600 | 50% | 375.9 |
| Trimer (ExpiCHO) | 600 | 600 | 600 | 600 | 600 | 600 | 600 | 600 | 0% | 600.0 |

| W11 CNE8 | M1 | M2 | M3 | M4 | M5 | M6 | M7 | M8 | RR | Geomean |
|---|---|---|---|---|---|---|---|---|---|---|
| I3-01v9 SApNP ($CHO^{FucT3}$, Endo H) | 600 | 600 | 600 | 600 | 600 | 600 | 600 | 600 | 0% | 600.0 |
| Trimer ($CHO^{FucT3}$, Endo H) | 600 | 600 | 600 | 600 | 600 | 600 | 600 | 600 | 0% | 600.0 |
| Trimer ($CHO^{FucT3}$) | 600 | 600 | 600 | 600 | 600 | 600 | 600 | 600 | 0% | 600.0 |
| Trimer (ExpiCHO) | 600 | 600 | 600 | 600 | 600 | 600 | 600 | 600 | 0% | 600.0 |

### Fig. S5

**Fig. S5. Immunogenicity of $CHO^{FucT3}$-produced BG505 Env immunogens in wild-type mice.** (**a**) Serum IgG binding curves of mouse sera from $CHO^{FucT3}$-produced BG505 Env immunogen groups ($n$ = 8 mice per group) binding to ExpiCHO-produced BG505 UFO trimer. (**b**) (Top) Summary of geometric mean $EC_{50}$ titers measured for $CHO^{FucT3}$-produced BG505 Env immunogen groups. $EC_{50}$ titers were derived from serum IgG ELISA against ExpiCHO-produced BG505 UFO trimer. Color coding indicates $EC_{50}$ levels (green to red: low to high binding). (Bottom) Summary of statistical analysis performed for each timepoint. Note: $EC_{50}$ values at week 2 were derived by setting the minimum/maximum absorbance at 450 nm (A450) values to 0.0/2.5. (**c**) (Top) Summary of A450 values from serum IgE induced by $CHO^{FucT3}$-produced BG505 Env immunogens, measured against ExpiCHO-produced BG505 UFO trimer. (Bottom) Summary of statistical analysis performed for each timepoint. (**d**) (Top) Neutralization curves of week 11 serum from individual mice in $CHO^{FucT3}$-produced BG505 Env immunogen groups against the autologous tier 2 BG505.T332N pseudovirus and MLV pseudoparticles as a negative control. (Bottom 1) Summary of geometric mean $ID_{50}$ titers measured for $CHO^{FucT3}$-produced BG505 Env immunogen groups against BG505.T332N. Color coding: green to red (low to high neutralization). Note: ID50 values were calculated with neutralization constrained between 0 and 100%. (Bottom 2) Summary of statistical analysis performed for week 11. (**e**) Neutralization curves of purified week 11 IgG from individual mice immunized with $CHO^{FucT3}$-produced BG505 Env immunogens against a pseudovirus panel comprising the autologous tier 2 clade A BG505.T332N and six heterologous tier 2 strains: 398F1 (clade A), X2278 (clade B), X1632 (clade B), 25710 (clade C), BJOX2000 (CRF07_BC), and CNE8 (CRF01_AE). MLV pseudoparticles were included as a negative control. (**f**) Summary of neutralization $IC_{50}$ values of purified week 11 IgG. Color coding: green to red (low to high neutralization). Error bars represent the difference between duplicate measurements at each concentration tested for each sample. $EC_{50}$, $ID_{50}$, and $IC_{50}$ values were calculated using GraphPad Prism. Data were analyzed using one-way ANOVA, followed by Tukey's multiple comparisons post hoc test for each timepoint. For significance, ns (not significant), *$p < 0.05$, **$p < 0.01$, ***$p < 0.001$, and ****$p < 0.0001$.

### Fig. S6

**a** Serum IgG binding of individual humanized mice immunized with Hi5 and $CHO^{FucT3}$-produced BG505 UFO trimer vaccines

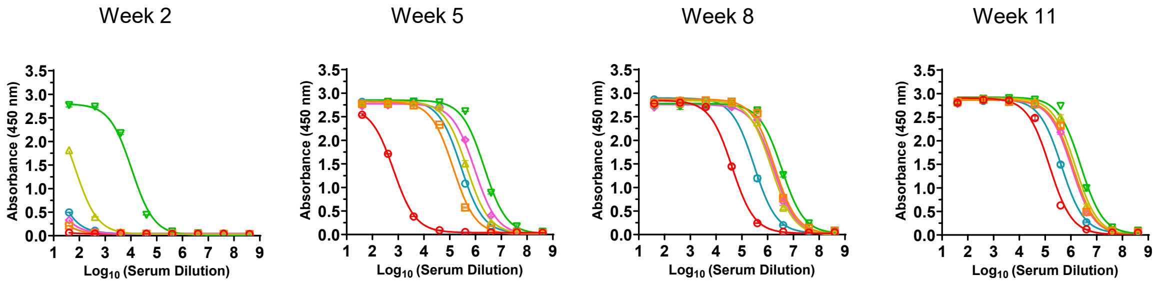


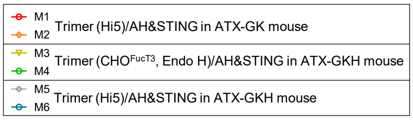


**b** Humanized mouse serum IgG ELISA $EC_{50}$ titers

| Antigen and mouse model | $EC_{50}$ titers (week 2) | | Geometric Mean |
|---|---|---|---|
| | M1 | M2 | |
| Trimer (Hi5)/AH&STING in ATX-GK mouse | 0.3 | 2.6 | 0.9 |
| Trimer ($CHO^{FucT3}$, Endo H)/AH&STING in ATX-GKH mouse | 69.5 | 10781.0 | 865.8 |
| Trimer (Hi5)/AH&STING in ATX-GKH mouse | 4.6 | 7.8 | 6.0 |

| Antigen and mouse model | $EC_{50}$ titers (week 5) | | Geometric Mean |
|---|---|---|---|
| | M1 | M2 | |
| Trimer (Hi5)/AH&STING in ATX-GK mouse | 656 | 135736 | 9439 |
| Trimer ($CHO^{FucT3}$, Endo H)/AH&STING in ATX-GKH mouse | 457659 | 2162820 | 994904 |
| Trimer (Hi5)/AH&STING in ATX-GKH mouse | 922347 | 271907 | 500792 |

| Antigen and mouse model | $EC_{50}$ titers (week 8) | | Geometric Mean |
|---|---|---|---|
| | M1 | M2 | |
| Trimer (Hi5)/AH&STING in ATX-GK mouse | 41474 | 1855694 | 277422 |
| Trimer ($CHO^{FucT3}$, Endo H)/AH&STING in ATX-GKH mouse | 1402653 | 3514587 | 2220303 |
| Trimer (Hi5)/AH&STING in ATX-GKH mouse | 1778375 | 299395 | 729683 |

| Antigen and mouse model | $EC_{50}$ titers (week 11) | | Geometric Mean |
|---|---|---|---|
| | M1 | M2 | |
| Trimer (Hi5)/AH&STING in ATX-GK mouse | 149115 | 1225066 | 427406 |
| Trimer ($CHO^{FucT3}$, Endo H)/AH&STING in ATX-GKH mouse | 1503403 | 2406935 | 1902260 |
| Trimer (Hi5)/AH&STING in ATX-GKH mouse | 1074532 | 434428 | 683233 |

**c** Humanized mouse serum IgE ELISA $OD_{450}$ values

| Antigen and mouse model | $OD_{450}$ values (week 2) | | Geometric Mean |
|---|---|---|---|
| | M1 | M2 | |
| Trimer (Hi5)/AH&STING in ATX-GK mouse | 0.046 | 0.050 | 0.048 |
| Trimer ($CHO^{FucT3}$, Endo H)/AH&STING in ATX-GKH mouse | 0.050 | 0.082 | 0.064 |
| Trimer (Hi5)/AH&STING in ATX-GKH mouse | 0.046 | 0.050 | 0.048 |

| Antigen and mouse model | $OD_{450}$ values (week 5) | | Geometric Mean |
|---|---|---|---|
| | M1 | M2 | |
| Trimer (Hi5)/AH&STING in ATX-GK mouse | 0.050 | 0.053 | 0.051 |
| Trimer ($CHO^{FucT3}$, Endo H)/AH&STING in ATX-GKH mouse | 0.048 | 0.050 | 0.049 |
| Trimer (Hi5)/AH&STING in ATX-GKH mouse | 0.067 | 0.050 | 0.058 |

| Antigen and mouse model | $OD_{450}$ values (week 8) | | Geometric Mean |
|---|---|---|---|
| | M1 | M2 | |
| Trimer (Hi5)/AH&STING in ATX-GK mouse | 0.051 | 0.060 | 0.055 |
| Trimer ($CHO^{FucT3}$, Endo H)/AH&STING in ATX-GKH mouse | 0.062 | 0.060 | 0.061 |
| Trimer (Hi5)/AH&STING in ATX-GKH mouse | 0.136 | 0.051 | 0.084 |

| Antigen and mouse model | $OD_{450}$ values (week 11) | | Geometric Mean |
|---|---|---|---|
| | M1 | M2 | |
| Trimer (Hi5)/AH&STING in ATX-GK mouse | 0.070 | 0.079 | 0.074 |
| Trimer ($CHO^{FucT3}$, Endo H)/AH&STING in ATX-GKH mouse | 0.125 | 0.123 | 0.124 |
| Trimer (Hi5)/AH&STING in ATX-GKH mouse | 1.013 | 0.056 | 0.237 |

### Fig. S6

**Fig. S6. Antibody responses induced by core α(1,3)-fucosylated BG505 UFO trimers in Alloy humanized mice.** (**a**) Serum IgG binding curves of Alloy humanized mouse sera elicited by Hi5- and $CHO^{FucT3}$-produced BG505 UFO trimers (n = 2 mice per group) against ExpiCHO-produced BG505 UFO trimer. (**b**) Summary of $EC_{50}$ titers derived from serum IgG ELISA against the ExpiCHO-produced BG505 UFO trimer. Color coding indicates $EC_{50}$ levels (green to red: low to high binding). Note: $EC_{50}$ values at week 2 were derived by setting the minimum/maximum A450 values to 0.0/2.8, respectively. (**c**) IgE responses in Alloy humanized mouse sera elicited by Hi5- and $CHO^{FucT3}$-produced BG505 UFO trimers (*n* = 2 mice per group) against ExpiCHO-produced BG505 UFO trimer. Given the low abundance of IgE antibodies in serum, we used A450 values to represent serum IgE binding to the ExpiCHO-produced BG505 UFO trimer.

**Fig. S7**

a

**Primers for heavy chain variable domain PCR1, PCR2, and sanger sequencing (ATX-Gx™ User Guide, page 22).**

| Primer | Forward/Reverse | Sequence | Description |
|---|---|---|---|
| **Heavy chain PCR1** | | | |
| **HB-006** | F(PCR1) | ACAGGTGCCCACTCCCAGGTGCAG | 5′ L-VH1 |
| **HB-007** | F(PCR1) | AAGGTGTCCAGTGTGARGTGCAG | 5′ L-VH3 |
| **HB-008** | F(PCR1) | CCCAGATGGGTCCTGTCCCAGGTGCAG | 5′ L-VH4/6 |
| **HB-009** | F(PCR1) | CAAGGAGTCTGTTCCGAGGTGCAG | 5′ L-VH5 |
| **HB-002** | R(PCR1) | TAGCCCTTGACCAGGCATCC | IgHG_R1 |
| **HB-003** | R(PCR1) | TAGCCTTTGACAAGGCATCC | IgHG_R2 |
| **Heavy chain PCR2** | | | |
| **HM-005** | F(PCR2) | GCATTAGTAACGAATTCACAGGTACAGCTGCAGCAGTCA | VHFR Forward1 |
| **HM-006** | F(PCR2) | GCATTAGTAACGAATTCACAGGTGCAGCTGCAGGAGTCSG | VHFR Forward2 |
| **HM-007** | F(PCR2) | GCATTAGTAACGAATTCACAGGTGCAGCTACAGCAGTGGG | VHFR Forward3 |
| **HM-008** | F(PCR2) | GCATTAGTAACGAATTCAGAGGTGCAGCTGKTGGAGWCC | VHFR Forward4 |
| **HM-009** | F(PCR2) | GCATTAGTAACGAATTCACAGRTCACCTTGAAGGAGTCTG | VHFR Forward5 |
| **HM-010** | F(PCR2) | GCATTAGTAACGAATTCASAGGTDCAGCTGGTGSARTCTGG | VHFR Forward6 |
| **HM-092** | R(PCR2) | TGAGGAGACRGTGACCAGGGTG | VHFR Reverse1 |
| **HM-093** | R(PCR2) | TGAGGAGACGGTGACCAGGGTT | VHFR Reverse2 |
| **HM-094** | R(PCR2) | TGAAGAGACGGTGACCATTGT | VHFR Reverse3 |
| **HM-095** | R(PCR2) | TGAGGAGACGGTGACCGTGGTCC | VHFR Reverse4 |
| **VHFseq** | Sequencing primer | GCATTAGTAACGAATTCASAG | Sequencing primer |

Heavy chain PCR1:
VH Forward Mix: **HB-006, HB-007, HB-008, HB-009**
VH Reverse Mix: **HB-002, HB-003**
Heavy chain PCR2:
VH Forward Mix: **HM-005, HM-006, HM-007, HM-008, HM-009, HM-010**
VH Reverse Mix: **HM-092, HM-093, HM-094, HM-095**

**Primers for VK PCR1, PCR2, and sanger sequencing (ATX-Gx™ User Guide, pages 22-23)**

| Primer | Forward/Reverse | Sequence | Description |
|---|---|---|---|
| **Kappa light chain PCR1** | | | |
| **HB-018** | F(PCR1) | ATGAGGSTCCCYGCTCAGCTGCTGG | 5′ LVκ 1/2 |
| **HB-019** | F(PCR1) | CTCTTCCTCCTGCTACTCTGGCTCCCAG | 5′ LVκ 3 |
| **HB-020** | F(PCR1) | ATTTCTCTGTTGCTCTGGATCTCTG | 5′ LVκ 4 |
| **HB-001** | R(PCR1) | CTGAGGCACCTCCAGTTGCTAACTGTTCCG | IgKC_R |
| **Kappa light chain PCR2** | | | |
| **HM-020** | F(PCR2) | GCATTAGTAACGAATTCAGACATCCRGDTGACCCAGTCTCC | VK FR Forward1 |
| **HM-021** | F(PCR2) | GCATTAGTAACGAATTCAGATATTGTGMTGACBCAGWCTCC | VK FR Forward2 |
| **HM-022** | F(PCR2) | GCATTAGTAACGAATTCAGAAACGACACTCACGCAGTCTC | VK FR Forward3 |
| **HM-023** | F(PCR2) | GCATTAGTAACGAATTCAGAAATTGTRWTGACRCAGTCTCC | VK FR Forward4 |
| **HM-024** | R(PCR2) | AGGGGCAGCTACCGTACGTTTGATTTCCACCTTGGTCC | VK FR Reverse1 |
| **HM-025** | R(PCR2) | AGGGGCAGCTACCGTACGTTTGATCTCCASCTTGGTCC | VK FR Reverse2 |
| **HM-026** | R(PCR2) | AGGGGCAGCTACCGTACGTTTGATATCCACTTTGGTCC | VK FR Reverse3 |
| **HM-027** | R(PCR2) | AGGGGCAGCTACCGTACGTTTAATCTCCAGTCGTGTCC | VK FR Reverse4 |
| **VKFseq** | Sequencing primer | GCATTAGTAACGAATTCAGAH | Sequencing primer |

Kapp chain PCR1:
VK Forward Mix: **HB-018, HB-19, HB-020**
VK Reverse: **HB-001**
Kapp chain PCR2:
VK Forward Mix: **HM-020, HM-021, HM-022, HM-023**
VK Reverse Mix: **HM-024, HM-025, HM-026, HM-027**

### Fig. S7

**b** Amino acid sequences of Env-specific antibodies isolated by single-cell sorting from G2M2 (Group 2, Alloy Mouse 2), Plate 2

```
>B2_HC
QVQLVQSGGGVVQPGRSLRLSCAASGFTFSGYGMHWVRQAPGKGLEWVAVIWYDGSNKYYADSVKGRFTISRDNSKNTLYLQMNSLRTEDTAVYYCARDRYCFGGTCSEGYFDYWGQGTPVTVSS
>B2_KC
EIVLTQSPLSLPVTLGQPASISCRSSQSLVHSDGYTYLNWFQQRPGQSPRRLIYKVSSRDSGVPDRFSGSGSGTDFTLKISRVEAEDVGVYYCMQGTHWITFGHGTRLEIK
>B6_HC
QVQLQESGPGLVKPSETLSLTCTVSGDSISSSSYSWGLIRQPPGKGLEWIGSIYYSGDTYYNPSLKSRVTITVDTSKNQFSLKLSSVTAADTAVYYCARRQFDYWGQGTLVTVSS
>B6_KC
EIVMTQTPATLSLSPGERATLSCRASQSISSYLVWYQQKPGQAPRLLIYDASNRATGIPARFSGSGSGTDFTLTISSLEPEDFAVYYCQQRNNWPLTFGGGTKVEIK
>B8_HC
QVQLQQWGAGLLKPSETLSLTCAVYGGSFSGYYWSWIRQSPGKGLEWIGEIDHSGSTNYNPSLKSRVTISVDTSKNQLSLKLSSATAADTAVFYCARGGDYYDNTGYYYWGQGTLVTVSS
>B8_KC
EIVMTQSPSSLSASVGDRVTITCQASQDISNYLNWYQQKPGKAPKLLIYDASILKAGFPSRFSGSGSGTDFTFTIGSLQPEDIATYYCQQYDSLPVTFGGGTKVEIK
>B10_HC
QVQLVESGGGLVKPGGSLRLSCAASGFSFSLYTVNWVRQAPGRGLDWVSSISGNSNHINYADSVKGRFAISRDNAKNSLYLQMNSLRAEDTAVYYCARELYGGNGDYWGQGTLVTVSS
>B10_KC
EIVMTQSPSSLSASVGDRVTITCQASQDIDKYLNWYQXXPGKAPKLLIYDASNLETEVPSRFSGSGSGTDFTFTIISLQPEDFATYYCQQYDNLPYTFGQGTKLEIK
>C3_HC
QVQLVESGGGLVKPGGSLRLSCAASGFIFSSYTMNWVRQAPGKGLEWVSSISSNSDYIYYEDSVKGRFTISRDNAKNSLYLQMHSLTVEDTAVYYCARDLVGATSYNWFDPWGQGTLVTVSS
>C3_KC
IVMTQTPSSLSASVGDRVNITCQASQDISNYLDWYQQKPGKAPKLLIYDASNLETGVPSRFSGSGSGTDFTFTISSLQPEDIATYYCQQYDNLPLTFGGGTKVEIK
>C5_HC
QVQLQESGGGVVQPGRSLRLSCAASGFNFRNYGMHWVRQAPGKGLEWVALIWFDGNKKHSANSVMDRFTISRDNSKNTLFLQMNSLRAEDTAVYYCARDGITGTMRYYYYYMDVWGKGTTVTVSS
>C5_KC
EIVLTQSPSLLPASVGDRVTITCRASQGISSYLAWYQQKPGKAPNLLIYAASTLQSGVPSRFSGSGSGTEFTLTISSLQSEDFATYYCQQLNNYPPGFGGGTKVDIK
>C8_HC
QVQLVQSGGGLVKPGGSLRLSCAASGFSFIYYTMNWVRQAPGKGLEWVSSISSRSSDIYYADSVRGRFTISRDNAKNSLFLQMNSLRADDTAVYFCARELYGGNGDYWGQGTLVTVSS
>C8_KC
EIVLTQTPSSLSASVGDRVTITCQASQYINSYLNWFQQKPGNAPKLLIYDESILETGVPSRFSGSGSGTDFTFTISSLQPEDIATYYCQQYDNIPYTFGQGTKVEIK
>C10_HC
QVQLQESGPGLVKPSETLSLTCSVSGDSINSSPYSWGWIRQPPGKGLEWIGSIYDNGDTYYNPSLKSRVTISVDTSKNQFSLMVNSVTAADTAVYYCARHQFDYWGQGTLVTVSS
>C10_KC
EIAMTQSPDTLSLSPGERATLSCRASQSTDSYLTWYQQKPGQAPRLLIYDASNRATGVPARFSGGGSGTDFTLTISNLEPEDFAVYYCQQRSTWPLTFGGGTKVEIK
>C11_HC
QVQLQQWGPGLVKPSETLSLTCTVSGASISGSSYYWGWIRQPPGKGLEWIGSIYYSGDTYYNPSLKSRVTISVDTSKNQFSLKLSSVTAADTAVYYCVRHQFDYWGQGTLVTVSSGTTVTVSS
>C11_KC
EIVMTQSPATLSLSPGERATLSCRASQSVSSYLVWYQQKPGQAPRLLIYDASNKATGIPARFSGSGPGTDFTLTISSLEPEDFAVYYCQQRNNWPLTFGGGTKVEIK
>D9_HC
QVQLVESGGGLVKPGGSLRLSCAASGFSFSLYTMNWVRQAPGKGLDWVSSISSTTSIYYADSVKGRFSISRDNAKNSLYLQMNSLRAEDTAVYYCARELYGGNGDYWGQGTLVTVSS
>D9_KC
EIVMTQSPSSLSASVGDRVTITCQASQDIDRYLNWYQQKPGKAPKLLIYDASNLEIGVPSRFSGSGSVTDFTFTIISLQPEDFATYYCQQFDNLPYTFGQGTKVEIK
>E3_HC
QVQLQESGGGLVKPGESLRLSCEASGLTLSSYSMSWVRQTPGKGLEWVSSITSSSFHIYYADSVKGRFTISRDNAKNSLYLQMSSLRAEDTALYYCARGWLQGIDYWGQGTLVTVSS
>E3_KC
EIVLTQSPGTLSLSPGYRATLSCRPSHSVGSSYLAWYQQKPGQAPRLLIYGASSRATGIPDRFSGSGSGTDFTLTISRLEPEDFAVYYCQQYGTSPWTFGQGTKVEIK
>E6_HC
QVQLQESGPGLVKPSETLSLTCTVSGASISSSSYYWGWIRQPPGKGLEWIGSIYYSGDTYYNPSLKSRVTISVDTSKNQFSLKLSSVTAADTAVYYCARHQFDYWGQGTLVTVSS
>E6_KC
EIVMTQSPATLSLSPGERATLSCRASQSVSSYLVWYQQKPGQAPRLLIYDASNGATGIPARFSGSGSGTDFTLTISSLEPEDFAVYYCQQRNNWPLTFGGGTKVEIK
>E11_HC
QVQLQQWGPGLVKPSQTLPLTCAISGFSVSSNTAAWNWIRQSPSRGLEWLGRTYYRSKWYNDYAVSVKSRITINPDTSKNQFSLQLNSVTPEDTAVYYCARDRVQVEPDAFDIWGQGTTVTVSS
>E11_KC
EIVMTQSPGTLSLSPGYRATLSCRPSHSVSSSYLAWYQQKPGQAPRLLIYGASSRATGIPDRFSGSGYGTDFTLTISRLEPEDFAVYYCQQYGTSPWTFGQGTKVEIK
>F5_HC
QVQLQESGGGLVQPGRSLRLSCAASGFTFDDYAMHWVRQAPGKGLEWVSGISWNSGSIGYADSVKGRFTISRDNAKNSLYLQMNSLRAEDTALYYCAKDRELRYFDFDYWGQGTLVTVSS
>F5_KC
EIALTQSPSSLSASVGDRVTITCRASQSISSYLNWYQQKPGKAPKLLIYAASSLQSGVPSRFSGSGSGTDFTLTINSLQPEDFATYYCQQSYSTPLTFGGGTKVDIK
>F8_HC
QVQLVQSGGGLVKPGGSLRLSCAASGFSFSYFTINWVRQAPGKGLEWVSSISGSTSNIYYTDSVQGRFTTSRDNAKNSLYLQMNTLRAEDTGVYYCAREVYGGNGDYWGQGTLVTVSS
>F8_KC
EIVLTQSPSSLSASVGDRVSITCQASQDIWKYLNWYQQKPGKAPKLLIYDASILETGVPSRFSGSGSGTDFTFIISSLQPEDFATYYCQQYDNLPYTFGLGTKVEIK
>F9_HC
QVQLXESGGGVVQPGRSLRLSCAASGFTFRNYGMHWVRQAPGKGLEWVAVIWFDGSIKHHVDSVKGRFTISRDNSKNTLYLQMNSLRAEDTALYYCARDGITGTTRYYYYYMDVWGEGTTVTVSS
>F9_KC
EIVLTQSPSFLSASVGDRVTITCRASQGISSYVAWYQQKPGKAPKLLIYAASILQSGVPSRFSGSGSGTEFTLTISSLQPEDFAIYYCQQLNSSPPTFGGGTKLEIK
>G4_HC
QVQLVESGGGVVQPGRSLRLSCAASGFTFSGYGMHWVRQAPGKGLEWVAVIWYDGSNKYYADSVKGRFTISRDNSKNTLYLQMNSLRTEDTAVYYCARDRYCFGGTCSEGYFDYWGQGTLVTVSS
>G4_KC
EIVLTQSPSFLSASVGDRVTITCRASQGISSYLAWYQQKPGKAPNLLIYAASTLQSGVPSRFSGSGSGTEFTLTISSLQPEDFATYYCQQLNSSPPTFGGGTKVDIK
>G10_HC
QVQLQESGGGVVQPGRSLRLSCAASGFTFRNYGMHWVRQAPGKGLEWVAVIWFDGGLRYYADSVKGRFTISRDNSKNTLFLQMDSLRAEDTAVYYCARDGITGTTRYYYYYMDVWGKGTTVTVSS
>G10_KC
EIVMTQSPSFLSASVGDRVTITCRASQGISSYLAWYQQKPGKAPNLLIYAASTLQSGVPSRFSGSGSGTEFTLTISSLQPEDFATYYCQQLNSSPPTFGGGTKVDIK
```

### Fig. S7

c ELISA binding of selected human antibodies (IgG format) to ExpiCHO-produced BG505 UFO trimer

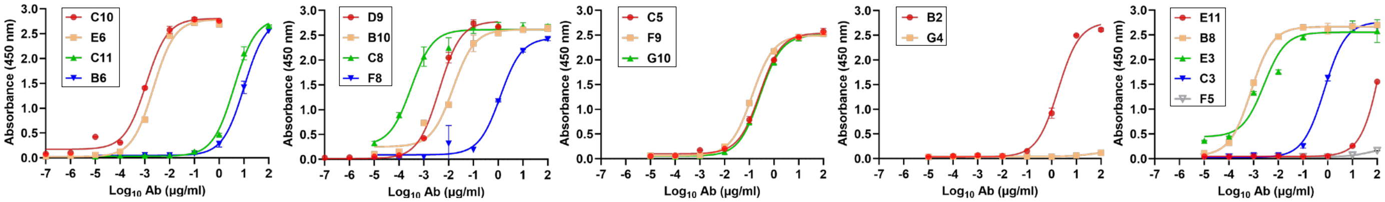


Coated with ExpiCHO-produced BG505 UFO trimer

| C10 | | E6 | | C11 | | B6 | | D9 | | B10 | |
|---|---|---|---|---|---|---|---|---|---|---|---|
| $EC_{50}$ | SD | $EC_{50}$ | SD | $EC_{50}$ | SD | $EC_{50}$ | SD | $EC_{50}$ | SD | $EC_{50}$ | SD |
| 0.001135451 | 1.05E-05 | 0.002206847 | 0.000026 | 4.075973749 | 0.4745 | 10.30149504 | 1.785 | 0.004189924 | 0.0004055 | 0.015285598 | 0.00151 |

| C8 | | F8 | | C5 | | F9 | | G10 | | B2 | |
|---|---|---|---|---|---|---|---|---|---|---|---|
| $EC_{50}$ | SD | $EC_{50}$ | SD | $EC_{50}$ | SD | $EC_{50}$ | SD | $EC_{50}$ | SD | $EC_{50}$ | SD |
| 0.000319203 | 0.00007865 | 1.226458316 | 0.079 | 0.266873566 | 0.02905 | 0.131444475 | 0.0082 | 0.274640784 | 0.00225 | 1.839751614 | 0.1175 |

| G4 | | E11 | | B8 | | E3 | | C3 | | F5 | |
|---|---|---|---|---|---|---|---|---|---|---|---|
| $EC_{50}$ | SD | $EC_{50}$ | SD | $EC_{50}$ | SD | $EC_{50}$ | SD | $EC_{50}$ | SD | $EC_{50}$ | SD |
| 47.78177058 | 5.07 | 217.7195444 | 19.6 | 0.000783901 | 0.000025 | 0.002846255 | 0.0012865 | 0.731533253 | 0.04555 | 50.44223231 | 5.535 |

d ELISA binding of selected human antibodies (Fab format) to ExpiCHO-produced BG505 UFO trimer

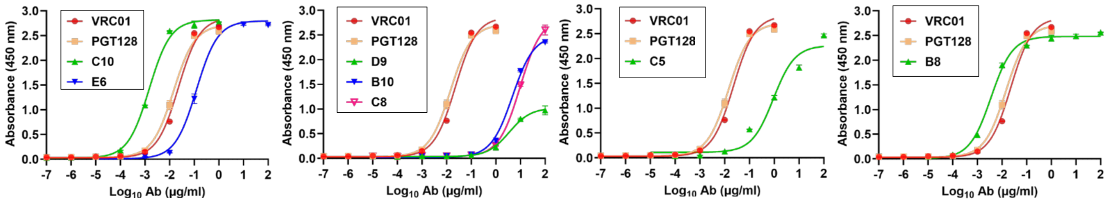


Coated with ExpiCHO-produced BG505 UFO trimer

| VRC01 | | PGT128 | | C10 | | E6 | | D9 | | B10 | |
|---|---|---|---|---|---|---|---|---|---|---|---|
| $EC_{50}$ | SD | $EC_{50}$ | SD | $EC_{50}$ | SD | $EC_{50}$ | SD | $EC_{50}$ | SD | $EC_{50}$ | SD |
| 0.022699029 | 0.00021 | 0.014402864 | 0.00176 | 0.001484655 | 3.2E-05 | 0.122989756 | 0.01065 | 3.515613176 | 0.1425 | 4.757106473 | 0.2045 |

| C8 | | C5 | | B8 | |
|---|---|---|---|---|---|
| $EC_{50}$ | SD | $EC_{50}$ | SD | $EC_{50}$ | SD |
| 9.64137179 | 0.8695 | 0.9194211 | 0.03905 | 0.00358874 | 0.000153 |

e **ExpiCHO-produced BG505 UFO trimer binding to selected human antibodies isolated from the sorting plate 2, Mouse 2 of Group 2 by BLI (AHC)**

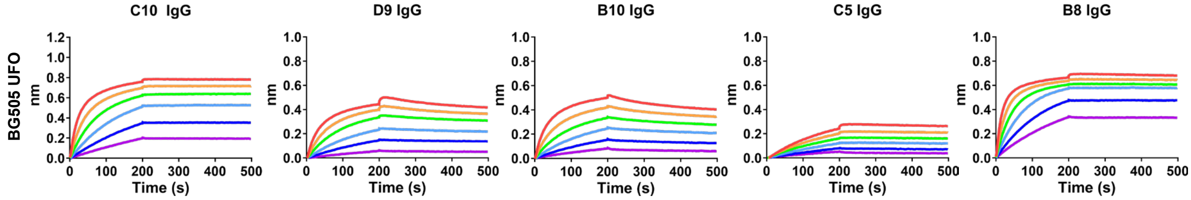

### Fig. S7

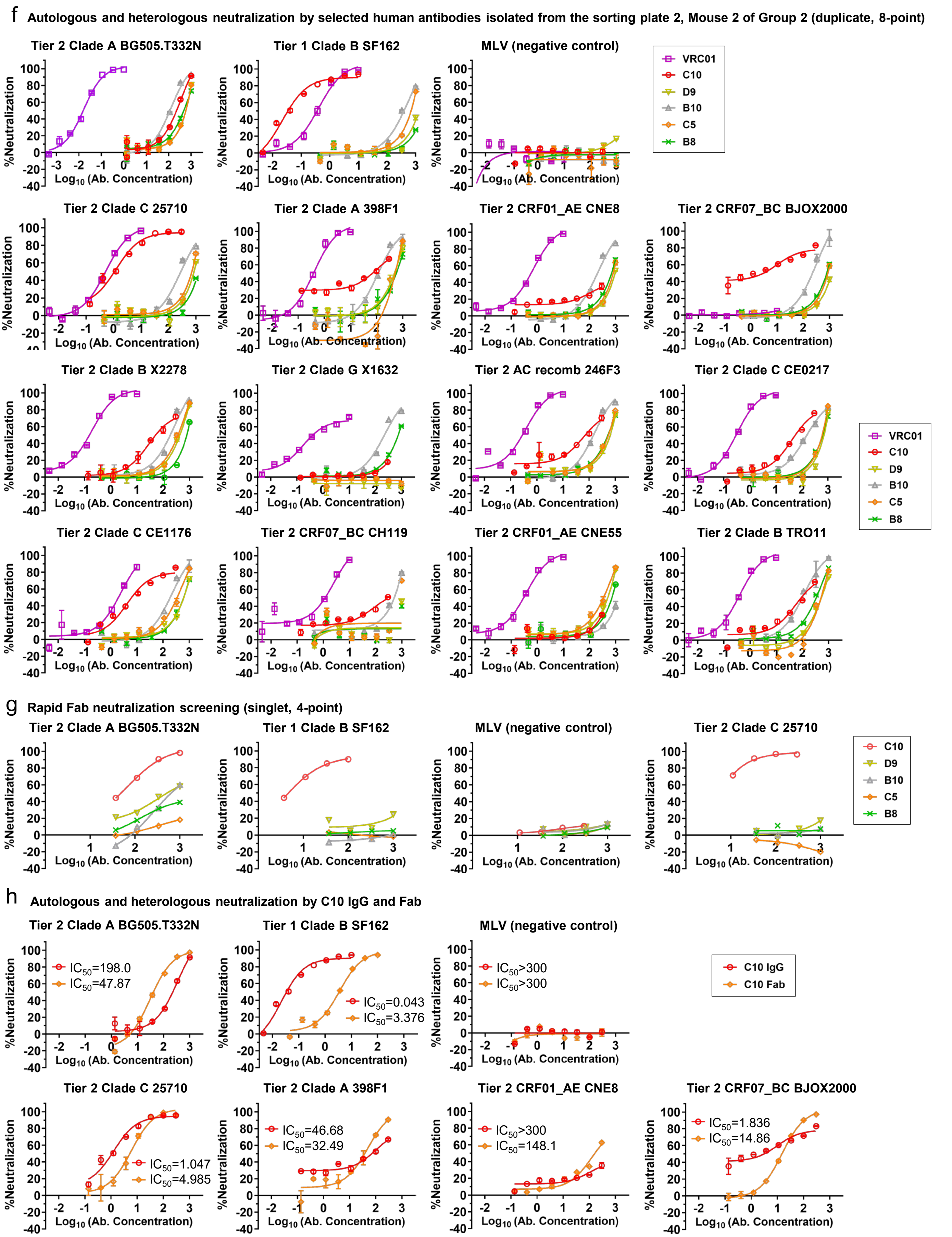
f Autologous and heterologous neutralization by selected human antibodies isolated from the sorting plate 2, Mouse 2 of Group 2 (duplicate, 8-point)
Tier 2 Clade A BG505.T332N
Tier 1 Clade B SF162
MLV (negative control)
VRC01
C10
D9
B10
C5
B8
Tier 2 Clade C 25710
Tier 2 Clade A 398F1
Tier 2 CRF01_AE CNE8
Tier 2 CRF07_BC BJOX2000
Tier 2 Clade B X2278
Tier 2 Clade G X1632
Tier 2 AC recomb 246F3
Tier 2 Clade C CE0217
Tier 2 Clade C CE1176
Tier 2 CRF07_BC CH119
Tier 2 CRF01_AE CNE55
Tier 2 Clade B TRO11
%Neutralization
Log10 (Ab. Concentration)
g Rapid Fab neutralization screening (singlet, 4-point)
Tier 2 Clade A BG505.T332N
Tier 1 Clade B SF162
MLV (negative control)
Tier 2 Clade C 25710
h Autologous and heterologous neutralization by C10 IgG and Fab
Tier 2 Clade A BG505.T332N
IC50=198.0
IC50=47.87
Tier 1 Clade B SF162
IC50=0.043
IC50=3.376
MLV (negative control)
IC50>300
IC50>300
C10 IgG
C10 Fab
Tier 2 Clade C 25710
IC50=1.047
IC50=4.985
Tier 2 Clade A 398F1
IC50=46.68
IC50=32.49
Tier 2 CRF01_AE CNE8
IC50>300
IC50=148.1
Tier 2 CRF07_BC BJOX2000
IC50=1.836
IC50=14.86

h (continued)

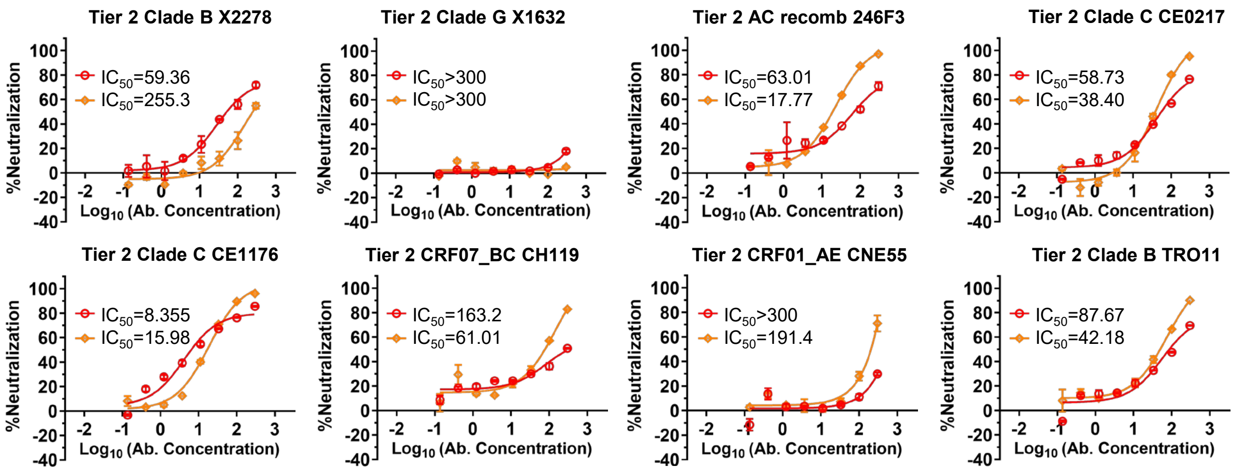


**Fig. S7. Genetic and functional characterization of single-cell-sorted human antibodies from an Alloy transgenic mouse with cross-clade serum neutralization.** (**a**) Heavy- and light-chain variable gene primers used in antibody cloning after single-cell sorting. (**b**) Amino acid sequences of BG505 UFO Env-specific antibodies isolated from mouse G2M2 (Group 2, Alloy mouse 2), Plate 2. (**c**) ELISA binding of selected human antibodies (IgG format) to ExpiCHO-produced BG505 UFO trimer. (**d**) ELISA binding of selected human antibodies (Fab format) to ExpiCHO-produced BG505 UFO trimer. In (**c**) and (**d**), binding curves are shown at the top, and corresponding $EC_{50}$ values with standard deviations are summarized in the tables below. Plates were coated with 0.1 μg of antigen, and antibodies were tested in a 10-fold dilution series starting at 1 μg ml$^{-1}$. Error bars represent the variation between duplicate measurements at each concentration tested for each sample. (**e**) Env binding by selected human antibodies isolated from mouse G2M2 (Group 2, Alloy mouse 2), Plate 2, measured by BLI using AHC analytical biosensors. A twofold concentration gradient of ExpiCHO-expressed BG505 UFO trimer, starting at 266 nM, was used in a six-point dilution series. (**f**) Autologous and heterologous neutralization by selected human antibodies isolated from mouse G2M2 (Group 2, Alloy mouse 2), Plate 2. For autologous tier 2 clade A BG505.T332N neutralization, tier 1 clade B SF162 and MLV were included as positive and negative controls, respectively. The tier 2 12-virus global panel was tested to evaluate cross-clade neutralization. Assays were performed in duplicate with an 8-point dilution series. (**g**) Rapid Fab neutralization screening against autologous tier 2 clade A BG505.T332N, tier 1 clade B SF162, and MLV as a negative control. Tier 2 clade C 25710 from the 12-virus panel was included as a representative heterologous isolate. Screening assays were performed in singlet with a 4-point dilution series. (**h**) Autologous and heterologous neutralization by C10 IgG and Fab. The same viruses as in (f) were tested to evaluate C10 Fab potency and breadth.

### Fig. S8

**a** **Simulated 30-Å EM density maps of BG505 UFO trimer models in complex with representative bNAb and RM20A3 Fabs**

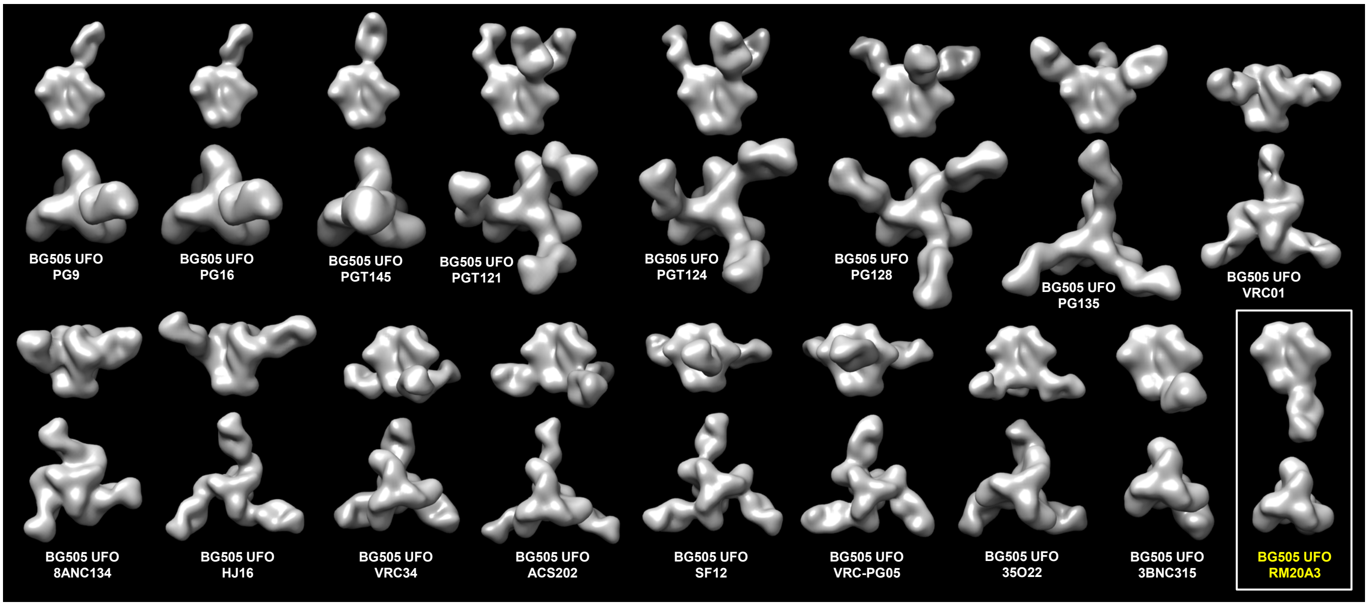


**b**

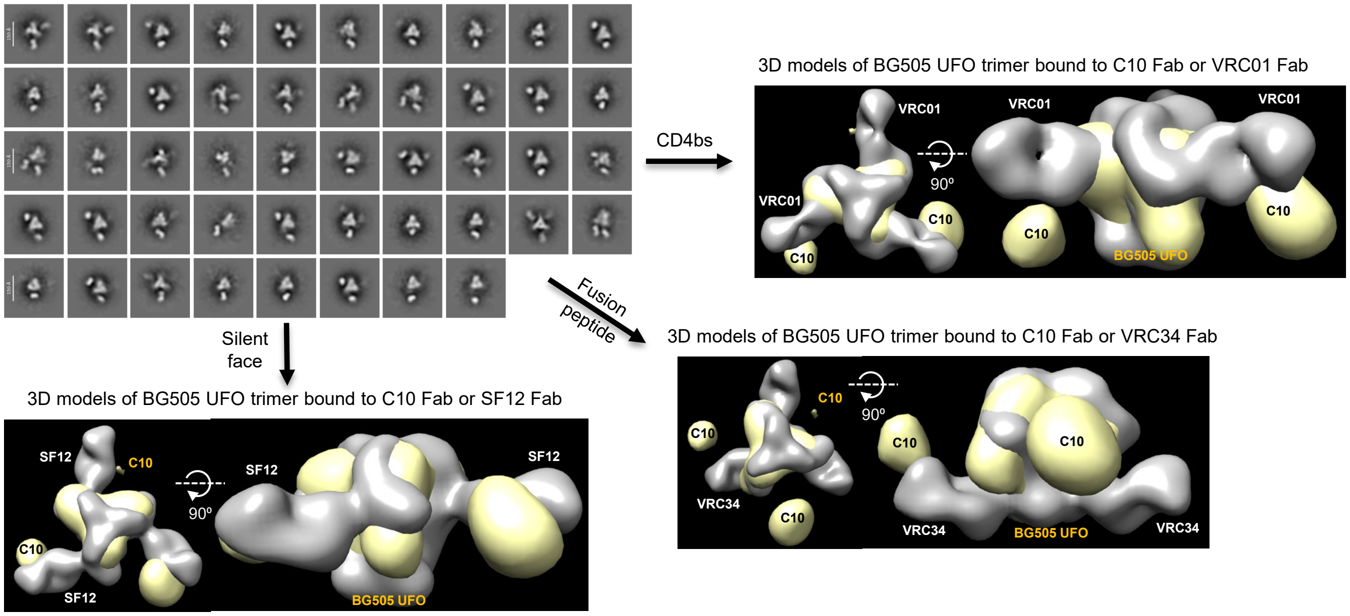


**c** Additional 2D and 3D analysis of CHO$^{FucT3}$-produced Endo H-trimmed BG505 UFO trimer bound to B10 Fab (**SEC-purified**)

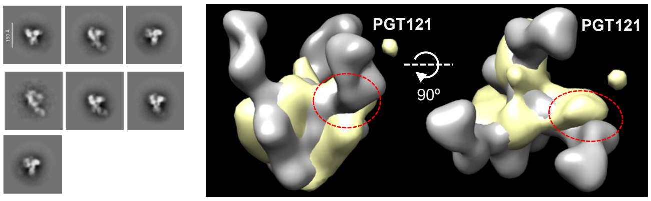


Additional 2D and 3D analysis of CHO$^{FucT3}$-produced Endo H-trimmed BG505 UFO trimer bound to B10 Fab (**without SEC**)

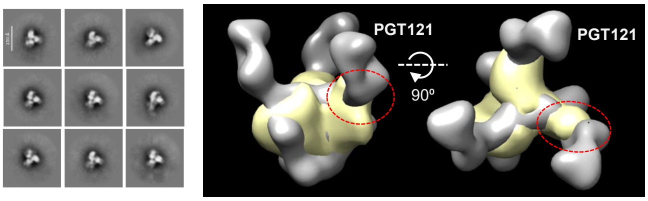

### Fig. S8

d **Antibodyomics pipeline for AVITI sequencing data and downstream bioinformatics analysis**

**Step 1**
**Fastq Data Preparation and Cleanup**

Raw AVITI sequencing data are initially processed to standardize read headers and assign a unique sequence index (e.g., index=12345) to each read. Due to the single-end sequencing configuration, reads extending beyond the platform-defined read boundaries contain terminal poly-C padding. These trailing poly-C sequences are removed to retain only antibody variable region sequences for subsequent analysis steps.

**Step 2**
**V(D)J Segment Identification and Somatic Divergence Quantification**

High-quality preprocessed reads are aligned against standard reference databases to identify and assign the closest corresponding germline variable (V), diversity (D), and joining (J) gene segments. A stringent alignment significance threshold (E-value cutoff) are applied to establish initial germline gene boundaries and distinguish functional from nonfunctional sequences.

**Step 3**
**Indel Error Correction and Amino Acid Translation**

To distinguish low-frequency sequencing artifacts from genuine biological mutations, an alignment-based filter is implemented to scan the assigned V and J genes. Sequences displaying frameshift-inducing insertions or deletions, particularly within homopolymer tracts, are flagged as probable platform-specific artifacts and evaluated to determine whether indel correction improved germline sequence identity. The corrected nucleotide sequences are then translated into functional amino acid sequences.

**Step 4**
**Sequence Identity Comparison Against User-Specified Antibody Panels**

Pairwise sequence identity is calculated between the corrected nucleotide and amino acid sequences and a user-defined panel of reference antibody chains, typically isolated by experimental approaches such as single-cell sorting. These comparisons enable direct identification of variants according to their homology to the target antibody panel.

**Step 5**
**CDR3 Determination, Comparison, and Variable Domain Extraction**

The final step of the pipeline determines the boundaries of complementarity-determining region 3 (CDR3) and extracts the complete variable domain. CDR3 sequences and variable domains are systematically compared with a user-defined panel of reference sequences using local sequence alignment. These comparisons enable identification and clonal tracking of target antibody variants across the processed repertoire.

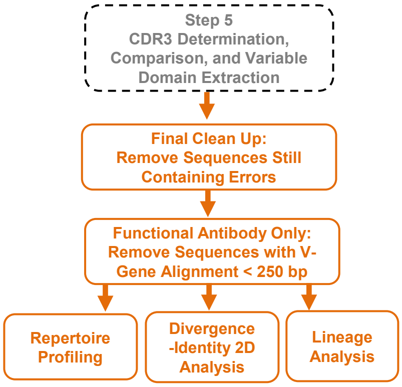


e **Antibodyomics pipeline processing of AVITI sequencing data for G2M1 and G2M2 splenic B-cell populations.** [a,b]

| Animal | Chain | $N_{Step1}$ | <Length> (nt) | $N_{Step2}$ | $N_{step3}$ | $N_{step4}$ | $N_{step5}$ | $N_{Final}$ | $N_{Functional}$ |
|---|---|---|---|---|---|---|---|---|---|
| G2M1 | H | 10,996,368 | 511.8 | 10,958,598 | 10,958,598 | 10,958,598 | 10,486,075 | 8,442,669 | 8,431,222 |
| | κ | 10,824,951 | 511.7 | 10,770,321 | 10,770,321 | 10,770,321 | 10,455,427 | 8,596,039 | 8,582,206 |
| G2M2 | H | 12,236,956 | 511.8 | 12,200,525 | 12,200,525 | 12,200,525 | 11,675,322 | 9,642,717 | 9,626,715 |
| | κ | 8,084,088 | 511.7 | 8,055,888 | 8,055,888 | 8,055,888 | 7,853,117 | 7,199,377 | 7,188,210 |

[a] Primers used to prepare antibody chain libraries are described in Fig. S7.
[b] Listed items include animal name, chain type, number of reads at Step 1, average read length after removal of terminal poly-C at Step 1 (in principle, 512 or $2^9$ bp), number of reads at Step 2 that could be reliably assigned to a heavy- or light-chain germline V gene using an E-value cutoff of $10^{-3}$, number of reads at Steps 3 and 4 (identical to that at Step 2), number of reads at Step 5 after removal of sequences lacking complete variable domains, number of reads after final Step 5 processing to remove sequences with translation errors or abnormalities, and number of usable antibody chains after removal of sequences with a germline V-gene alignment of less than 250 bp.

f **Heavy chain germline V-gene usage**

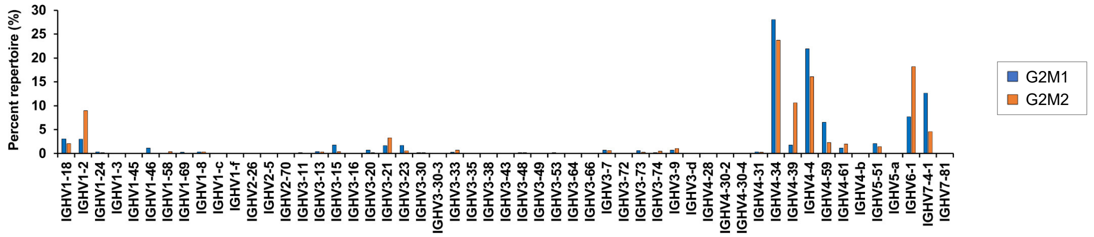


**Light (κ) chain germline V-gene usage**

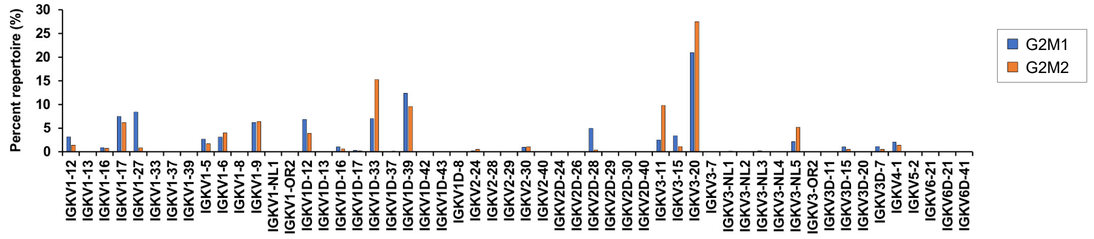


g **Somatic variants of C10, B10, C5, and B8 antibody lineages and their frequencies within the respective germline families.** [a,b]

| Antibody | | VH/K family | | Lineage | Percentage | | VH/K family | | Lineage | Percentage |
|---|---|---|---|---|---|---|---|---|---|---|
| C10 | | VH4-39 | 1,020,118 | 400,614 | 39.3% | | VH4-39 | 153,329 | 61 | 0.04% |
| | | VK3-11 | 708,842 | 8,971 | 1.3% | | VK3-11 | 214,580 | 111 | 0.05% |
| B10 | | VH3-21 | 318,421 | 84,830 | 26.6% | | VH3-21 | 137,506 | 27 | 0.02% |
| | **G2M2** | VK1/D-33 | 1,099,869 | 475,732 | 43.3% | **G2M1** | VK1/D-33 | 606,017 | 35,669 | 5.9% |
| C5 | | VH3-33 | 66,917 | 546 | 0.8% | | VH3-33 | 21,235 | 1 | 0.005% |
| | | VK1-9 | 462,180 | 38 | 0.008% | | VK1-9 | 530,812 | 0 | 0% |
| B8 | | VH4-34 | 2,285,704 | 17,034 | 0.7% | | VH4-34 | 2,364,433 | 3 | 0% |
| | | VK1/D-33 | 1,099,869 | 12,119 | 1.1% | | VK1/D-33 | 606,017 | 1033 | 0.17% |

[a] Heavy- and light-chain libraries were generated and sequenced separately and were not paired; therefore, the numbers of heavy and light chains within each antibody lineage may not match.
[b] The four antibodies were isolated from G2M2 and are shown on the left. Analysis of G2M1 sequences was performed to identify similar antibody chains and determine whether similar sequences may arise in a different animal immunized with the same vaccine.

#### h C10 and B10 heavy-chain putative germline precursors and somatic or high-identity variants identified in G2M2 and G2M1

```
>C10-GH (C10 germline VH, v-gen mismatch=0)
QLQLQESGPGLVKPSETLSLTCTVSGGSISSSSYYWGWIRQPPGKGLEWIGSIYYSGSTYYNPSLKSRVTISVDTSKNQFSLKLSSVTAADTAVYYCARHQFDYWGQGTLVTVSS

>B10-GH (B10 germline VH, v-gen mismatch=0)
EVQLVESGGGLVKPGGSLRLSCAASGFTFSSYSMNWVRQAPGKGLEWVSSISSSSSYIYYADSVKGRFTISRDNAKNSLYLQMNSLRAEDTAVYYCARELYGGNGDYWGQGTLVTVSS
```

---

```
>C10-VH1 (G2M2, index=16039 v-mismatch=31 seqid_aa=82.6 cdrh3_sid_nt=100.0)
QVQLQQWGAGLLKPSETLSLTCAVSGDSISGSPYSWGWLRQPPGKGLEWIGSIYYDGTAHYNPSLKSRVSMSVDTSKNQFSLRLRSVTVADTAIYYCARHQFDYWGQGTLVTVSS

>C10-VH2 (G2M2, index=1220498 v-mismatch=36 seqid_aa=84.3 cdrh3_sid_nt=100.0)
EVQLVETGPGLVKSSGTLSLTCSVSGDSISGSPYSWGWLRQPPGKDLEWIGSIYYDGDTHYNPSLKSRVSMSVDTSKNQFSLRLNSVTVADTAIYYCARHQFDYWGQGTLVTVSS

>C10-VH3 (G2M2, index=923299 v-mismatch=43 seqid_nt=84.1 cdrh3_sid_nt=100.0)
QVQLQQSGPGLVKPSQTLSLTCAISGDSVSSNSAAWNWIRQSPGKGLEWIGSIYDSGSTYYNPSLKSRVTISVDTSKNQFSLTLRSVTAADTAVYYCARHQFDYWGQGTLVTVSS

>C10-VH4 (G2M2, index=11264719 v-mismatch=47 seqid_aa=80.9 cdrh3_sid_nt=100.0)
QVQLQQSGPGLVKPSQTLSLTCAISGDSVSSNSSCYWSWIRQPPGKGLEWIGAINHSGSTKYSPSLKSRLTISLDTSKNQFSLKLRSVTAADTAVYYCARHQFDYWGQGTLVTVSS

>C10-VH5 (G2M2, index=4046246 v-mismatch=51 seqid_aa=80.9 cdrh3_sid_nt=100.0)
QVQLQQSGLGLVKPSQTLSLTCAISGDSVSNNSAAWNWIRQSPSRGLEWIGSIYDSGSTYYNPSLKSRVTISVDTSKNQFSLKLSSVTATDTAVYYCARHQFDYWGQGTLVTVSS

>B10-VH6 (G2M2, index=359166 v-mismatch=22 seqid_aa=83.9 cdrh3_sid_nt=96.3
EVHLVESGEGLVKPGGSLRLSCGASGFTFSRYTIHWVRQAPGKGLEWVSSITGSSTYIYYADSVKGRFTISRDNAKNALYLQMNSLRVEDTAVYYCAREIYGGNGDYWGQGTLVTVSS

>B10-VH7 (G2M2, index=294377 v-mismatch=28 seqid_aa=83.9 cdrh3_sid_nt=100.0
QVTLKESGGDLVKPGGSPRLSCVASGFIFSNAWMSWVRQAPGKGLEWVSAISGSGASTNYADSVKGRFAISRDNAKNSLYLQMNSLRAEDTAVYYCARELYGGNGDYWGQGTLVTVSS

>B10-VH8 (G2M2, index=2033858 v-mismatch=34 seqid_aa=81.4 cdrh3_sid_nt=100.0
EVQLVESGPGLVKPSESLSLTCTVSGGSISGSSYYWGWVRQAPGKGLDWVSSISSSSTYIYYADSVKGRFTISRDNAKNSLYLQMNSLRAEDTAVYYCARELYGGNGDYWAQGTLVTVSS
```

---

```
>C10-VH1p (G2M1, putative variant, index=356363 v-mismatch=25 seqid_aa=85.2 cdrh3_sid_nt=100.0)
QVQLQQWGPGPVKPSETLSLTCTVSGDSIRDSIYSWGWIRQPPGKGLEWIGSIYDDGSTNYNPSLKSRVTMSVDTSKNQFSLKLSSVTATDTADYYCARHQFDYWDQGTLVTVSS

>B10-VH2p (G2M1, putative variant, index=5667635 v-mismatch=17 seqid_aa=86.4 cdrh3_sid_nt=100.0)
EVQLVESGGGLVKPGGSLRLSCAASGFTFSSYTMNWVRQAPGRGLEWVSSISSSSSFVYYADSVKGRFTISRDSAENSLYLQMHSLRAEDTAMYYCARELYGGNGDYWGQGTLVTVSS
```

## i

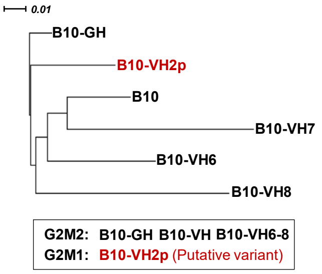


Autologous and heterologous HIV-1 neutralization by B10 germline and NGS-derived variants ($IC_{50}$, μg/mL) [a]

| Virus panel | | | G2M2 B10 germline and somatic variants | | | | G2M1 |
|---|---|---|---|---|---|---|---|
| Tier | Subtype | Virus | B10-GH | B10-VH6 | B10-VH7 | B10-VH8 | B10-VH2p |
| Tier 2 | A | BG505.T332N | >300 | 133.9 | >300 | 190.8 | >300 |
| Tier 1 | B | SF162 | >300 | >300 | >300 | >300 | >300 |
| — | — | MLV | >300 | >300 | >300 | >300 | >300 |
| Tier 2 | C | 25710 | >300 | >300 | >300 | >300 | >300 |

[a] Due to the lower potency of wild-type B10 and the low yields of the B10 germline and somatic variants, heterologous neutralization was evaluated against a single representative virus from the global panel, the tier 2 clade C virus 25710.

## j

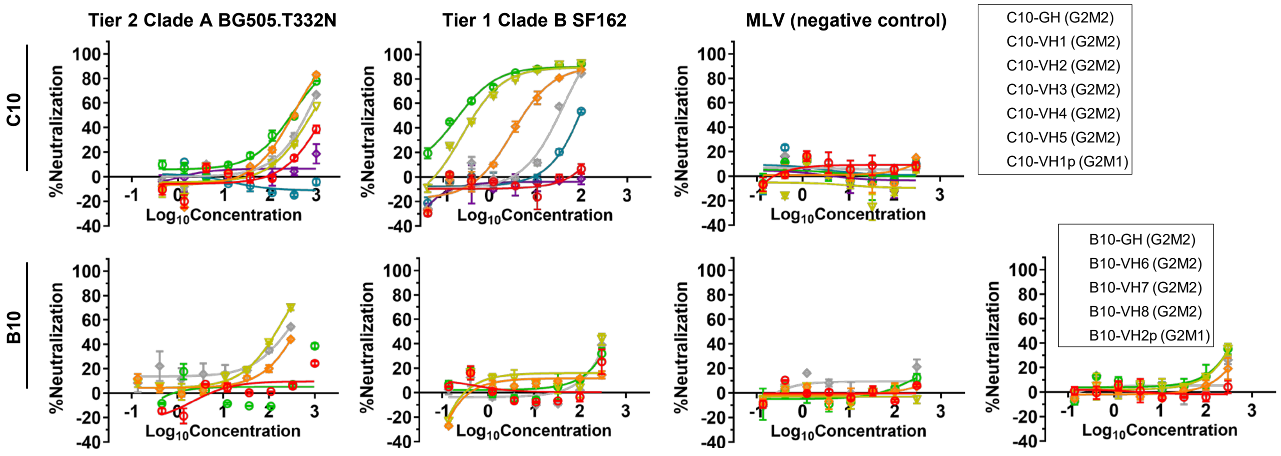

### Fig. S8

j (Continued)

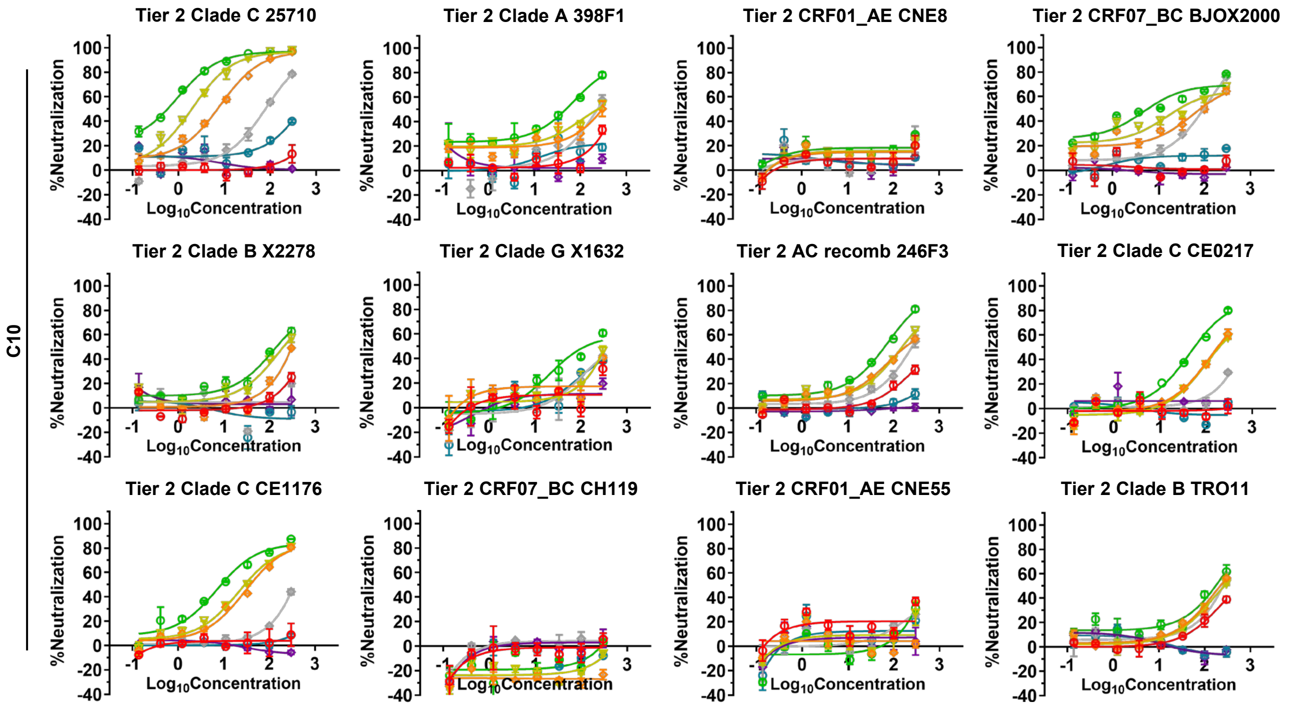


**Fig. S8. Epitope mapping and B-cell repertoire analysis of Alloy-derived human antibodies C10, B10, C5, and B8.** **(a)** Simulated 30-Å EM density maps of BG505 UFO trimer models in complex with Fabs of 16 representative bNAbs and the base-binding RM20A3. bNAbs target the V2 apex (PG9, PG16, PGT145), the N332 supersite (PGT121, PGT124, PGT128, PGT135), the CD4bs (VRC01, 8ANC134, HJ16), the FP (VRC34, ACS202), the SF (SF12, VRC-PG05), and the gp120/gp41 interface (35O22, 3BC315). The density maps were generated using pdb2mrc in EMAN2 and shown as gray surfaces. **(b)** Representative 2D class averages of Fab C10 binding to the ExpiCHO-produced BG505 UFO trimer (left) and its 3D reconstruction overlaid with the simulated maps of the VRC01, VRC34, and SF12 complexes (right and bottom). The C10/trimer complex density is shown as yellow surfaces, whereas the simulated bNAb/trimer complex density is shown as gray surfaces. **(c)** Representative 2D class averages and 3D reconstructions of Fab B10 binding to the $CHO^{FucT3}$-produced, Endo H-trimmed BG505 UFO trimer, with SEC purification (left) and without SEC purification (right) following complex formation. The 3D reconstructions of the B10/trimer complex are overlaid with the simulated map of PGT121 Fab bound to the trimer. The B10/trimer complex density is shown as yellow surfaces, whereas the simulated PGT121/trimer complex density is shown as gray surfaces. The EM density potentially corresponding to the B10-Env interaction site is highlighted with red circles. Of note, the C10/trimer and B10/trimer complex models in (b) and (c) were generated without enforcing C3 symmetry. **(d)** Modified Antibodyomics pipeline for AVITI sequencing data and downstream bioinformatics analysis. The pipeline flowchart is shown on the left, with key technical details shown next to the textboxes. Post-pipeline analysis steps are shown in orange on the right. **(e)** Summary of results obtained from the pipeline processing of AVITI sequencing data for G2M1 and G2M2 splenic B-cell populations. **(f)** Sub-family-level germline V-gene distributions for G2M1 and G2M2 heavy chains (HC, top) and κ-light chains (KC, bottom). **(g)** Summary of results from lineage analysis, listing somatic variants of the C10, B10, C5, and B8 antibody lineages and their frequencies within the respective germline families. **(h)** Amino acid sequences of inferred C10 and B10 HC precursors and somatic or high-identity variants identified in G2M2 and G2M1. The number of nucleotide mismatches in the V-gene alignment (`v-mismatch`), variable domain sequence identity to the parental antibody at the amino acid level, and HCDR3 sequence identity to that of the parental antibody at the nucleotide level are shown for each sequence. **(i)** Phylogenetic tree of B10 heavy-chain precursor and somatic variants, with neutralization potency reported as $IC_{50}$ values (µg/mL). **(j)** Neutralization curves of reconstituted C10 and B10 heavy-chain precursors and somatic variants. In (i) and (j), autologous tier 2 clade A BG505.T332N and tier 1 clade B SF162 were tested, with MLV serving as a negative control. The 12-virus global panel was then tested to evaluate the neutralization breadth of the C10 variants, whereas the tier 2 Clade C virus 25710 was used as a representative heterologous isolate for the B10 variants. The assays were performed in duplicate using an 8-point dilution series. Of note, two B10 variants, B10-GH and B10-VH7, were produced at sufficient yields to permit testing in the BG505.T332N neutralization assay at a starting concentration of 1 mg/mL. For all other TZM-bl assays involving B10 variants, the starting concentration was 300 µg/mL.
